# Glycolysis revisited: from steady state growth to glucose pulses

**DOI:** 10.1101/2022.06.22.497165

**Authors:** David Lao-Martil, Joep P.J. Schmitz, Bas Teusink, Natal A.W. van Riel

**Affiliations:** Department of Biomedical Engineering, Eindhoven University of Technology, Eindhoven, Noord-Brabant, 5612AE, The Netherlands; DSM Biotechnology Center, Delft, Zuid-Holland, 2613AX, The Netherlands; Systems Biology Lab, Vrije Universiteit Amsterdam, Amsterdam, Noord-Holland, 1081HZ, The Netherlands; Amsterdam University Medical Center, University of Amsterdam, Amsterdam, Noord-Holland, 1105AZ, The Netherlands

**Keywords:** Glycolysis, Growing Cell, Kinetic Modeling, Parameter Estimation, *Sacchamoyces cerevisiae*

## Abstract

Kinetic metabolic models of central metabolism have been proposed to understand how *Saccharomyces cerevisiae* navigates through nutrient perturbations. Yet, these models lacked important variables that constrain metabolism under relevant physiological conditions and thus have limited operational use such as in optimization of industrial fermentations. In this work, we developed a physiologically informed kinetic model of yeast glycolysis connected to central carbon metabolism by including the effect of anabolic reactions precursors, mitochondria and the trehalose cycle. A parameter estimation pipeline was developed, consisting of a divide and conquer approach, supplemented with regularization and global optimization. We show how this first mechanistic description of a growing yeast cell captures experimental dynamics at different growth rates and under a strong glucose perturbation, is robust to parametric uncertainty and explains the contribution of the different pathways in the network. Our work suggests that by combining multiple types of data and computational methods, complex but physiologically representative and robust models can be achieved.

## Introduction

*Saccharomyces cerevisiae*, commonly known as baker’s yeast, is a prominent cell factory for the biotechnology industry (Nielsen *et al*, 2013). In large-scale fermentations, dynamic gradients expose yeast cells to rapid nutrient changes in their extracellular environment, which in turn will impact intracellular metabolic regulation (Enfors *et al*, 2001; Haringa *et al*, 2016). To understand this dynamic stress response process, mechanistic models representing intracellular carbon metabolism kinetics have been developed (van Eunen *et al*, 2012; Smallbone *et al*, 2013; van Heerden *et al*, 2014). However, upscaling these models from lab to industry has proven challenging and restricted to simplified mechanistic growth models with little information on intracellular dynamics (Wang *et al*, 2020; Sarkizi Shams Hajian *et al*, 2020).

This could be partly explained because models lacked important physiological information. In the fields of pharmacology and medicine, validation of physiological realism is a necessary step in development of credible mechanistic model (Pruett *et al*, 2020). To develop physiologically representative yeast models, several aspects are to be considered. Current state of the art models study glycolysis in isolation (van Eunen *et al*, 2012; Smallbone *et al*, 2013; van Heerden *et al*, 2014), but should consider the complex cellular context it interacts with, for example by representing the trehalose and tricarboxylic acid (TCA) cycles.

The situation regarding biomass synthesis is an important example. Whereas maximization of biomass synthesis is the physiological function to which genome scale models (GSM) are optimized (Yasemi and Jolicoeur, 2021), it has been neglected in yeast kinetic networks, resulting in models that cannot represent a growing cell and thus poorly represent an important physiological state that comes with importable demands (van Heerden *et al*, 2015). Considering sink reactions for glycolytic intermediates is of upmost importance, as they were shown to be central in achieving the steady state (Teusink *et al*, 2000).

Moreover, variables alien to carbon flux regulate glycolysis as well. For instance, oxygen concentration determines if respiration or fermentation is performed (Otterstedt *et al*, 2004) and is routinely quantified in both lab and industrial scale bioreactors (Canelas *et al*, 2011; Haringa *et al*, 2016), but has not been used to constrain dynamic models. Furthermore, cofactors NAD and ATP are required to carry out multiple glycolytic reactions, but their availability cannot be taken for granted. ATP homeostasis is challenged under strong dynamic perturbations, where the inosine salvage pathway is used as transient store (Walther *et al*, 2010). In addition, ATP required for maintenance changes under different growth conditions (Chen and Nielsen, 2019). Still, a yeast model considering the effect of these variables does not exist.

Recently published data and tools make implementation of yeast models in a physiological context more feasible. The first perturbations studies in yeast were restricted to a single glucose pulse perturbation of 1 g L^-1^ (Theobald *et al*, 1997), but experimental quantification has been extended to larger perturbations that result in profound intracellular dynamics, including possible nonlinear effects of stress response (Walther *et al*, 2010; van Heerden *et al*, 2014). Furthermore, data on different growth steady states (SS) (van Eunen *et al*, 2012; Canelas *et al*, 2011) and industrially relevant feast/famine (FF) regimes (Suarez-Mendez *et al*, 2014) has also been generated. New experimental variables have been quantified, such as most metabolic species in central carbon metabolism (CCM) (Canelas *et al*, 2011), dynamic flux profiles (Suarez-Mendez *et al*, 2017), and proteome composition (Chen and Nielsen, 2019; Elsemman *et al*, 2022).

These data can now be used to quantify model parameters within a physiological range. Previous work showed that kinetic constants measured *in vitro* in isolated enzymes assays might not be representative of *in vivo* behavior (Teusink *et al*, 2000; Davidi and Milo, 2017). To overcome this issue, model parameters can be directly estimated to fit *in vivo* metabolomic and fluxomic data (Davidi and Milo, 2017). Few works estimated part of the yeast kinetic model parameter set in this way (Smallbone *et al*, 2013; Pritchard and Kell, 2002; Van Riel *et al*, 1998), but the data available has notably increased since then.

To address the abovementioned challenges, here we conceived a kinetic model of yeast glycolysis and trehalose cycle that considers the physiological effect of growth rate, gas exchange, ATP synthesis and maintenance, by developing a state-of-the-art approach for parameter identification. We show how this first mechanistic description of a growing yeast cell captures experimental dynamics at different growth rates and under a strong glucose perturbation, is robust to parametric uncertainty and explains the contribution of the different pathways in the network. Our work suggests that combining multiple types of data and computational methods, complex but physiologically representative and robust models can be achieved, and simultaneously points at specific locations in glycolytic models where regulation mechanisms are missing.

## Results

### An integrated, physiologically informed kinetic model of yeast glycolysis

Existing yeast glycolysis models studied this pathway in isolation (Smallbone *et al*, 2013; van Heerden *et al*, 2014). To transit from pathways in isolation, to pathways embedded in a growing cell, inclusion of physiological variables informative of the bioreactor environment is crucial (Lao-Martil *et al*, 2022). The required data to implement these variables is now available for *S. cerevisiae*. Carbon flux distribution in central metabolism and secretion of *O*_2_ and *CO*_2_ were determined at different dilution rates in chemostat (Canelas *et al*, 2011). Additionally, growth and non-growth -associated maintenance (GAM and NGAM, respectively) were determined and extensively implemented in GSMs and can be used to estimate the ATPase activity (Beeftink *et al*, 1990; Lu *et al*, 2019).

These variables have been included in the model developed in this work. Carbon metabolism includes glycolysis, trehalose cycle, glycerol branch and sink reactions to biomass and other by products (Fig 1). Production of CO2 (*q*_*CO*2_) consisted of the sum of what is released in the pyruvate decarboxylase (PDC) reaction and all the carbon moles that are processed in the mitochondria. The uptake of oxygen (*q*_*O*2_) was estimated using the *q*_*CO*2_ and the experimental respiratory quotient (RQ) value. Finally, the synthesis of ATP (*q*_*AT P*_) consisted of glycolytic production and mitochondrial contribution, estimated considering *q*_*O*2_ and P/O ratio (Verduyn *et al*, 1991). The maintenance consumption of ATP (*m*_*AT P*_) was estimated using the GAM and NGAM determined in GSMs (Lu *et al*, 2019).

**FIGURE 1.**
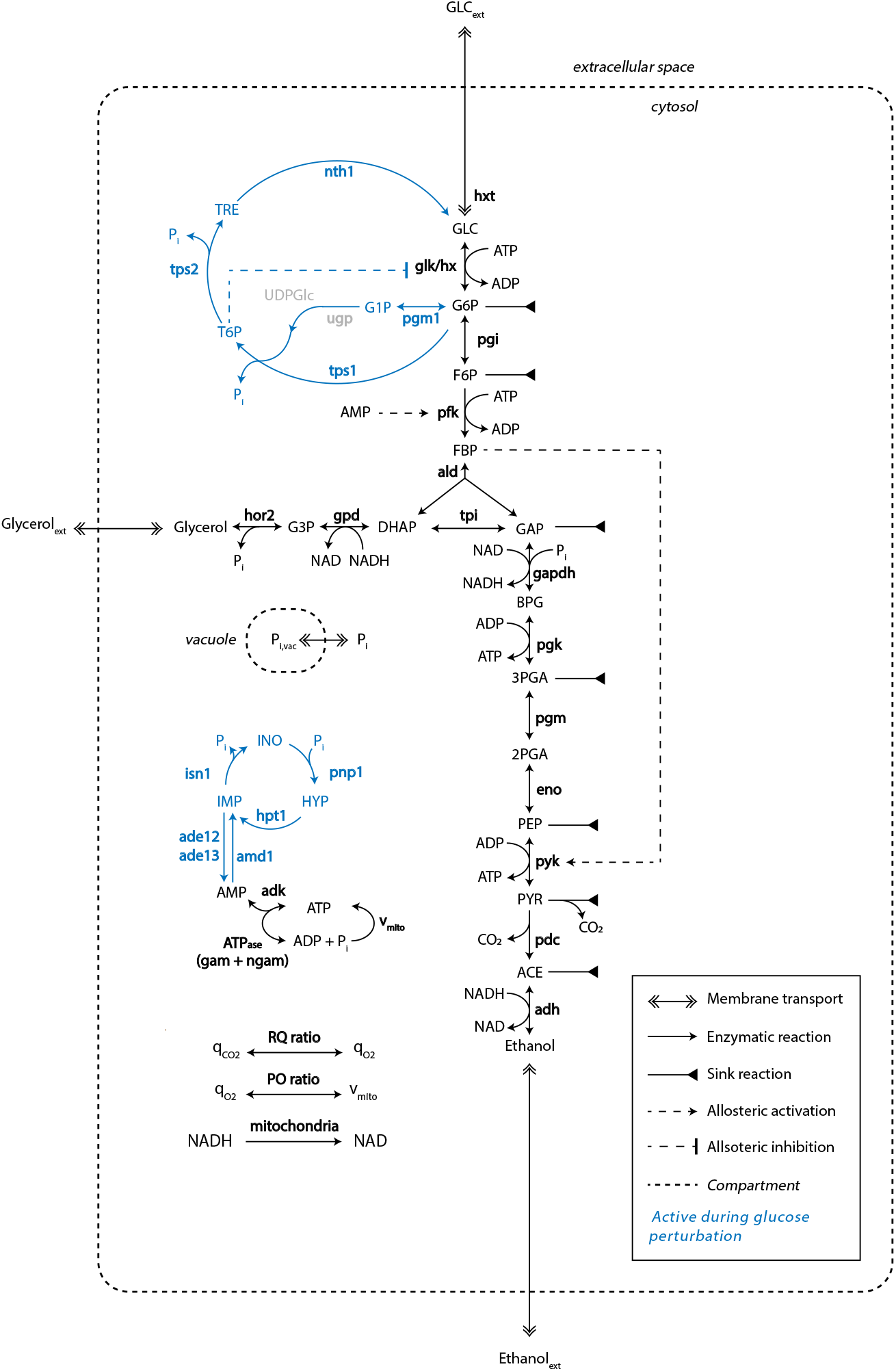
An updated central carbon metabolism *S. cerevisiae* model considering variables of physiological relevance. Black colored reactions are active for all simulations, blue only during the glucose perturbation and gray are lumped. Metabolites are connected by the reactions in the model (continuous lines), which are catalysed by enzymes (bold). Allosteric regulation is shown by semi-continuous arrows and compartments by semi-continuous lines.

The model consists of a series of nonlinear ordinary differential equations (ODE), where each mass balance describes a metabolite concentration, that changes over time according to reaction rates. Reactions that cannot be described by a single enzyme, such as the rates in which glycolytic intermediates are taken up to form biomass, are described by phenomenological expressions that closely resemble experimental data. The change in enzyme concentrations at different dilution rates found in (van Hoek *et al*, 2000) is adjusted for each enzyme. Model parameters were estimated to fit experimental data (see later sections). For further details on model implementation, parameters and kinetic expressions, see the materials and methods and supporting information.

### The model reproduced physiological properties of a growing cell in a bioreactor

The resulting model predicted experimental dynamics, differentiating respiratory and respirofermentative metabolism. At growth rates below 0.2 h^-1^, carbon uptake was directed to the sink reactions, which account for biomass synthesis and TCA cycle (Fig 2B). Above this threshold, metabolism gradually shifted to ethanol production which eventually became the main carbon product. In line with this, *q*_*CO*2_ closely resembled *q*_*O*2_ at low growth rates but became predominant above 0.2 h^-1^, discerning respiratory from respirofermentative metabolism (Fig 2C). Likewise, ATP was produced by respiratory metabolism at low growth rates and partially fermentative at high growth rates (Fig 2D). Maintenance reaction rate was close to a GAM of 40 mmol of ATP per g of biomass during the respiratory state but increased above this level as fermentation became predominant. A mismatch occurred at 0.2 h^-1^, when carbon flux was notable underestimated. We will discuss this later.

**FIGURE 2.**
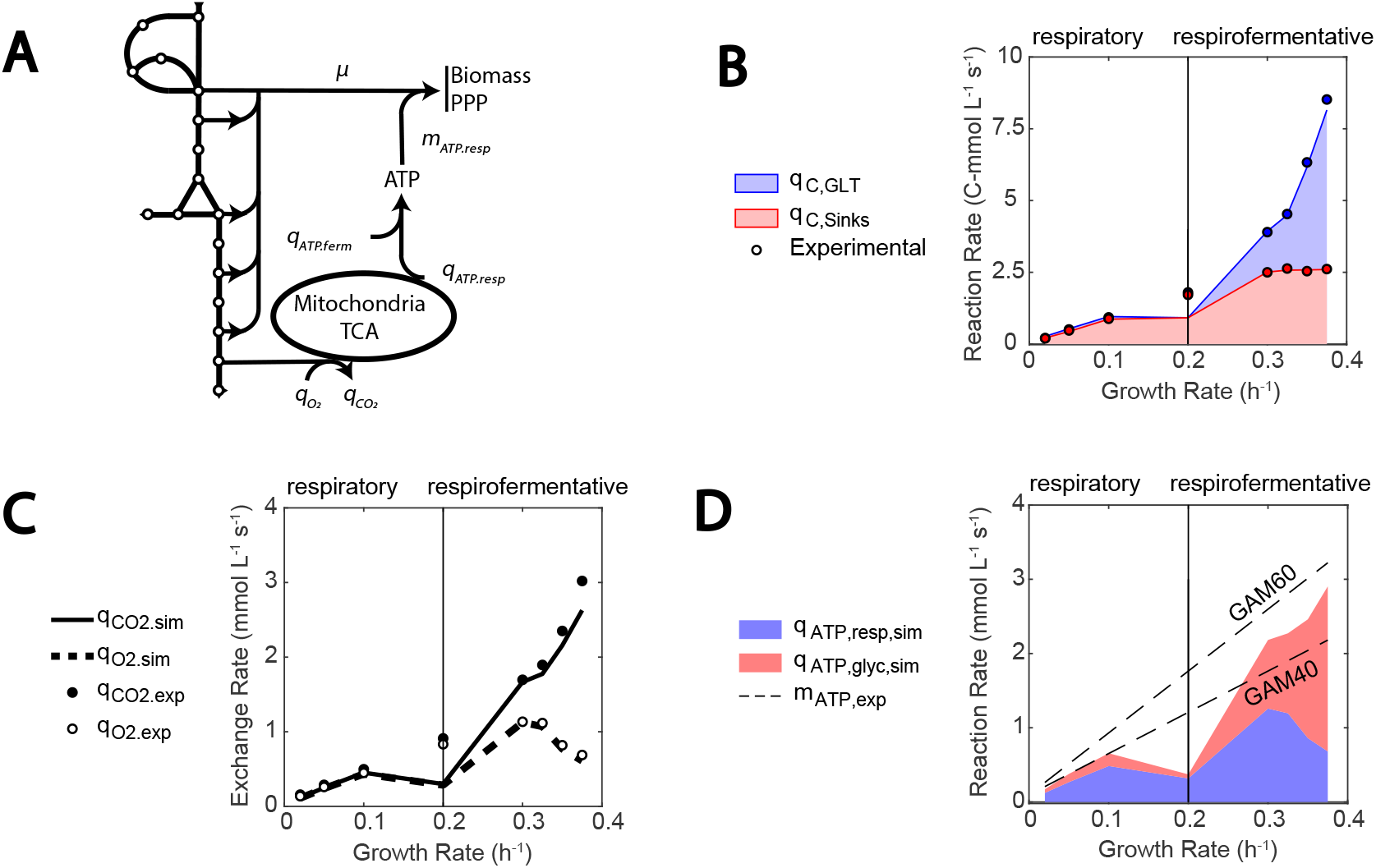
The model reproduces physiological properties of a growing cell in a bioreactor. (A) Simplified model scheme highlighting the role of sinks for anabolic reaction precursors, PPP and mitochondria (g6p, f6p, gap, pep, pyr, ace). (B) Carbon flux entering the sink reactions (red) compared to the total carbon uptake (blue). (C) Simulated exchange rate of *CO*_2_ (continuous line) and *O*_2_ (semi-continuous line). Experimental data shown as bullet points. (D) ATP produced by respiration (blue area) and fermentation (red area). Semi continuous lines point at the theoretical values calculated using the NGAM and GAM (40 and 60 mmol g_DW_^-1^) found in genome scale representations (Lu *et al*, 2019).

In conclusion, the agreement between model simulations and experimental data suggests that the model can work within the physiologically relevant region studied here. The implementation of the growth rate dependency seems essential. For instance, if only the ATP produced by glycolysis is considered, the overall maintenance needs cannot be matched at the different growth rates. This opens the possibility for kinetic models to be used in new setups, such as different growth rates or respiration/fermentation regimes, by using variables such as gas exchange rate as constraints, rather than as validation.

### Parameter estimation: A problem decomposition approach supplemented by regularization, parameter space sampling and cost function balancing

Quantification of the parameter set was performed by optimizing the model fit to multiple experimental setups and data types (van Heerden *et al*, 2014; Canelas *et al*, 2011; van Hoek *et al*, 2000). Nonetheless, the high number of unknown kinetic parameters made this task challenging due to multiple local optimum and ill-conditioning. To deal with non-identifiability problems in large kinetic models, multiple optimization methods have been proposed and evaluated (Villaverde *et al*, 2019). For instance, the so-called divide and conquer method aimed to analyze undetermined biochemical networks despite parameter uncertainty (Kotte and Heinemann, 2009). This problem decomposition approach splits the model in smaller parts where parameters are estimated and later assembled in a complete network.

To develop the S. cerevisiae model in this study, this problem decomposition method was adapted and supplemented with global optimization and regularization to improve convergence, akin to the approach described in (Gábor and Banga, 2015). Global optimization algorithms meet the danger presented by local minima by starting parameter estimation from a set of initial parameter guesses (Moles *et al*, 2003). Regularization methods reduce ill-conditioning by introducing prior knowledge on the parameters (Engl *et al*, 2009). Additionally, to control that overfitting did not occur, weighting factors were used to leverage the components of the cost function.

The parameter estimation pipeline is illustrated for the well-studied enzyme enolase (ENO) in (Fig 3). In the first stage, the divide and conquer problem decomposition approach was applied to glycolytic enzymes. As suggested in (Kotte and Heinemann, 2009), steady state data was used (Canelas *et al*, 2011; van Hoek *et al*, 2000). A regularization factor was added in the cost function, so that parameters resembled prior knowledge, provided that the model error was not increased (Fig 3A). The benefits of global optimization and regularization can be appreciated for parameters *V*_*max,E N O*_ and *K*_*M, F* 6*P, P F K*_ in Fig 3B and 3C, respectively. A parameter for phosphofructokinase (PFK) is shown since this is the most complex enzyme in the pipeline, and results differed according to enzyme kinetics complexity. While for ENO most initial parameter samples ended in the same parameter value as the literature one, the regularization factor brought estimated paraemters closer to literature values (Fig 3B). For PFK, global sampling revealed the existence of parameter dependencies (Fig 3C), and convergence towards literature values was achieved upon regularization. Since enzymatic kinetic constants vary within a large range, the parameters estimated were transformed from linear to logarithmic scale, as this has shown to be more reliable, also with global optimization methods (Kreutz, 2016; Villaverde *et al*, 2019).

**FIGURE 3.**
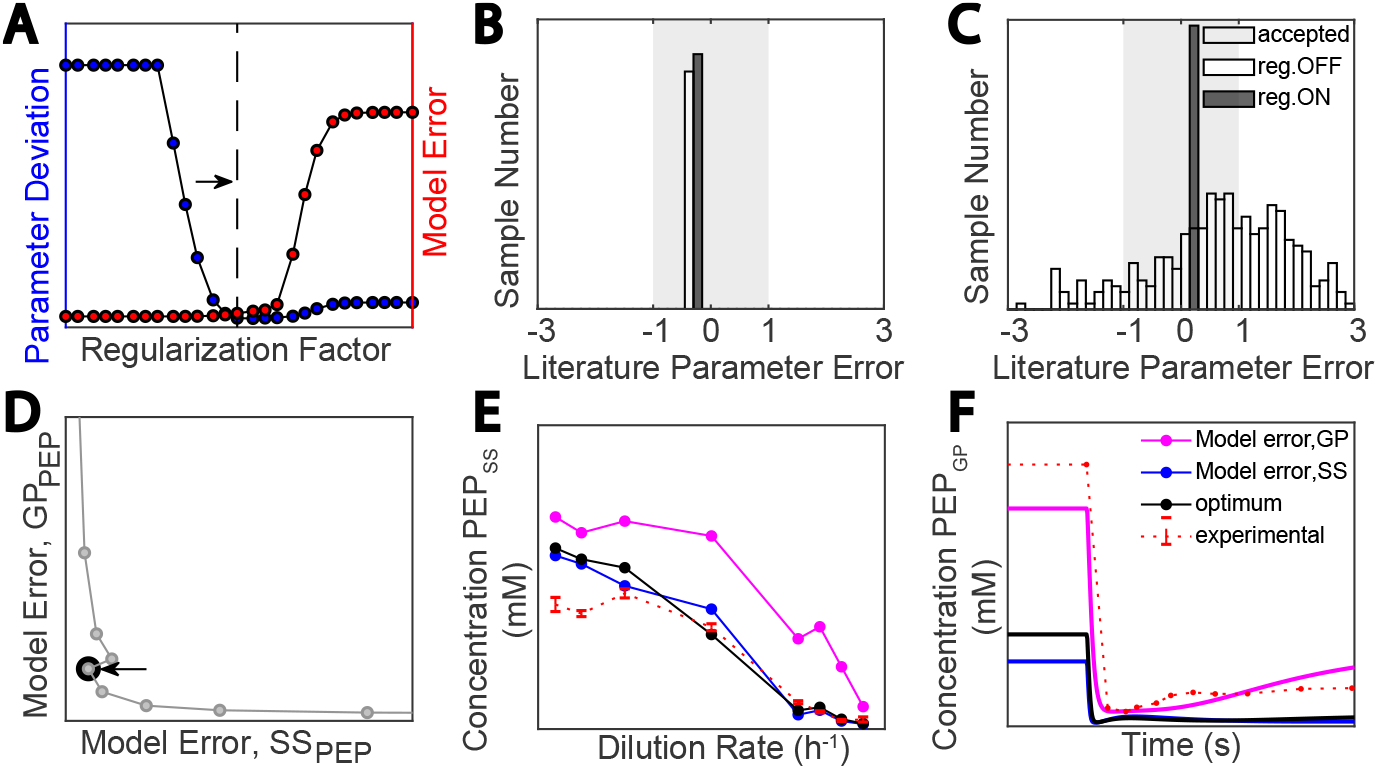
A problem decomposition approach supplemented by regularization, parameter sampling space and weighting data types. (A) Regularization approach: the regularization factor (x-axis), literature parameter deviation (blue, left y-axis) and model error (red, right y-axis) (B and C) Parameter estimation case for an identifiable (Enolase, *K*_*cat*_) and weakly identifiable parameter (Phosphofructokinase, *K*_*F* 6*P*_), respectively. Histogram plot showing the estimated parameter deviation from the literature value (x-axis) and the number of initial parameter samples in the y-axis. Non-regularized and regularized cases in white and dark bars, respectively. The shaded area covers the range of one order of magnitude of deviation from the literature parameter value. (D) Pareto front commonly obtained when fitting two types of data simultaneously: case of PEP concentration. Model error for steady state and glucose perturbation concentrations of PEP (x-axis and y-axis, respectively). (E and F) Model error, PEP steady state and glucose perturbation concentrations, respectively.

In the second stage, the developed models were assembled in a complete glycolysis model with trehalose cycle and cofactor metabolism. The weighting factors in the cost function were balanced ad hoc, until model fit for multiple data could not be further improved. This resulted in pareto fronts, such as the one for ENO (Fig 3D), where the fit for phosphoenolpyruvate (PEP) concentration in steady state and the 110 Mm glucose pulse was balanced (Fig 3E and 3F, respectively). See the supplementary information for an overview of the divide and conquer parameter estimation results, enzyme by enzyme.

### Combining different data types was central for physiological system identification

Determining a model parameter set, or system identification, is a data-intensive process (Almquist *et al*, 2014). To identify the model properties, excitation experiments such as impulse or step response and time variant and invariant data are used. Considering different experimental data types when building a kinetic metabolic model contributes to an accurate physiological description of the system (Peskov *et al*, 2012). In the case of yeast fermentations, early studies focused on the transient metabolic response to a glucose pulse (Theobald *et al*, 1997), but implementations have broadened to study the role of the trehalose cycle during higher residual glucose concentrations (van Heerden *et al*, 2014) and steady state dynamics were analyzed at different growth rates (Canelas *et al*, 2011). In the process, data types have also increased from describing metabolomics to fluxomics and even enzymatic constraints (van Hoek *et al*, 2000; Elsemman *et al*, 2022).

Therefore, the benefits of combining data types were examined for the developed model, in the areas highlighted in (Fig 4A). Simulations were compared when changing enzyme concentrations were considered as shown in (van Hoek *et al*, 2000) (data displayed in the supporting information), to when they were fixed at the dilution rate value of h^-1^ (Fig 4B). This analysis showed that inclusion of proteome changes improved the predicted fluxes substantially, especially at higher dilution rates, as the glycolytic flux is increased. This is supported by the relevance attributed to the proteome at different growth rates using GSMs (Sánchez *et al*, 2017; Elsemman *et al*, 2022).

**FIGURE 4.**
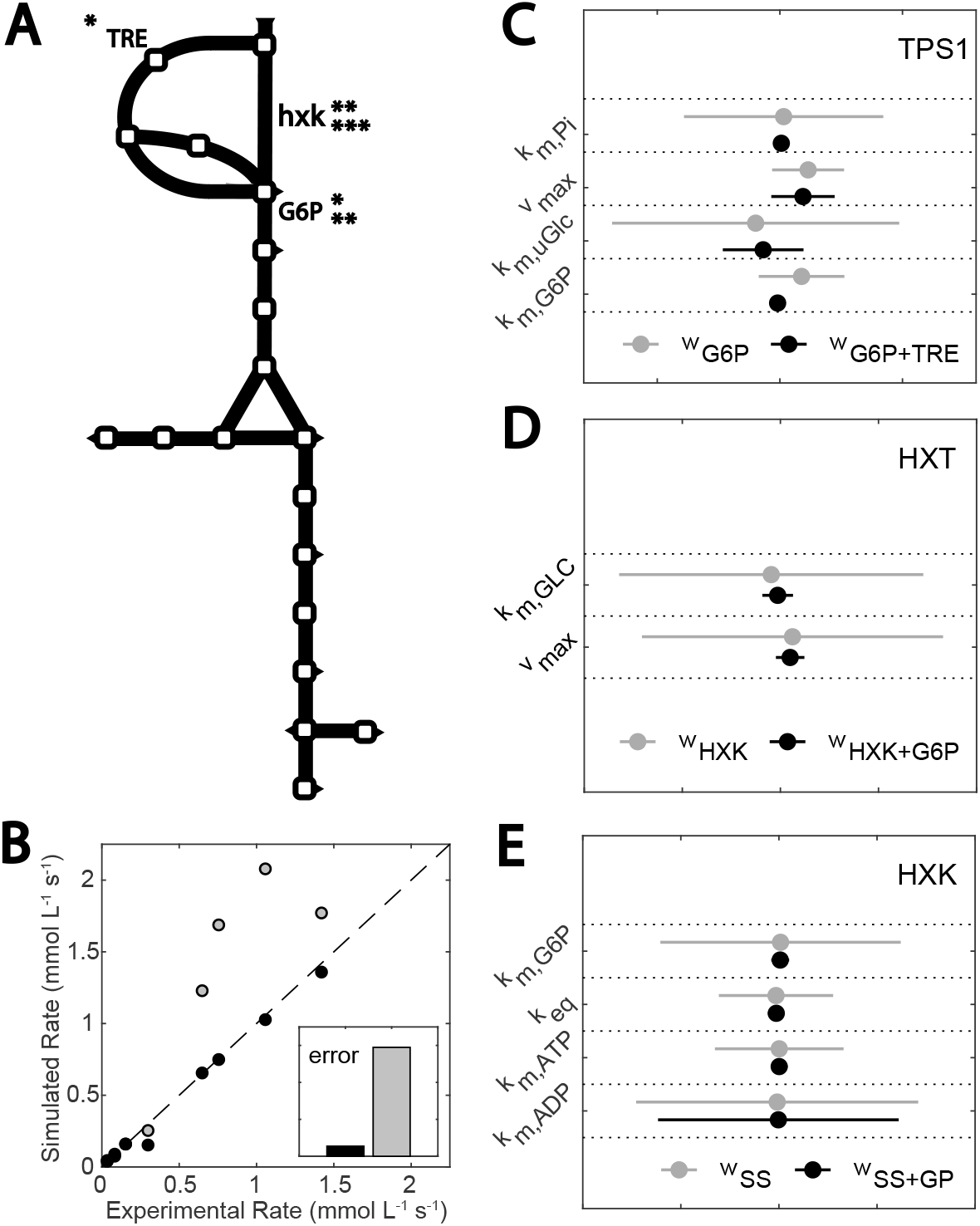
Contribution of different data sets to model identification. (A) Simplified model diagram: variables included in the TPS1 (*), HXT (**) and HXK (***) parameter estimation examples. (B) Simulated HXT reaction rates plotted against experimental rates. Including dilution rate dependency in enzyme concentrations (black) or keeping them constant at the experimental value found for 0.1 h-1. Confidence intervals obtained for the parameter estimates when: (C) G6P or the combination G6P+trehalose was used in TPS1 parameter estimation, (D) reaction rates (HXK) or the combination reaction rate and metabolite concentration (G6P) was used in HXT parameter estimation and (E) SS or the combination SS+GP data was included in HXK parameter estimation.

Furthermore, parameters were estimated to fit a single or combination of experimental variables, to show the benefits of using multiple data types when fitting the model. For instance, reduced confidence intervals were obtained for TPS1 kinetic constants if trehalose concentrations were used in combination with glucose 6-phosphate (G6P) (Fig 4C), and similar results were obtained combining metabolomic and fluxomic, or steady state and glucose perturbation data (Fig 4D and 4E, respectively). This tendency highlights how the identified system becomes more accurate, on top of more representative of cell dynamics. This is relevant, given that kinetic yeast CCM models have often been only developed with a single type of data, most usually a glucose perturbation for which only some metabolites were measured.

### Ensemble modelling suggested that the model was robust to parameter heterogeneity

Kinetic metabolic models are generally designed as deterministic descriptions of an average cell in the bioreactor, represented by a unique parameter set (Almquist *et al*, 2014). Nonetheless, heterogeneity between cells of the same population or between strains implies a certain degree of metabolic diversity (van Heerden *et al*, 2014; Nidelet *et al*, 2016; Rodrigues *et al*, 2021). These Bayesian dynamics can be represented by means of ensemble modelling, in which parameters are sampled from a variability distribution and are often used to account for uncertainty (Tan *et al*, 2011; Jia *et al*, 2012; Tiemann *et al*, 2013). In this work, a Monte Carlo approach was taken to assess how robust were the modelled glucose perturbation response and steady state dynamics. The models were generated by adding a 10% random noise to the parameter set.

Model simulations were found to be notably robust within the conditions tested. Predictions showed certain quantitative but little qualitative deviation (Fig 5). Steady state concentrations and fluxes were especially robust, as it has been found for other kinetic networks (Tran *et al*, 2008; Tan *et al*, 2011; Christodoulou *et al*, 2018). Variability was more pronounced during the glucose perturbation, which could be explained by the notably higher extracellular glucose concentrations (110 mM) experienced during the strong perturbation, when compared to the steady state range (0.13-3.30 mM).

**FIGURE 5.**
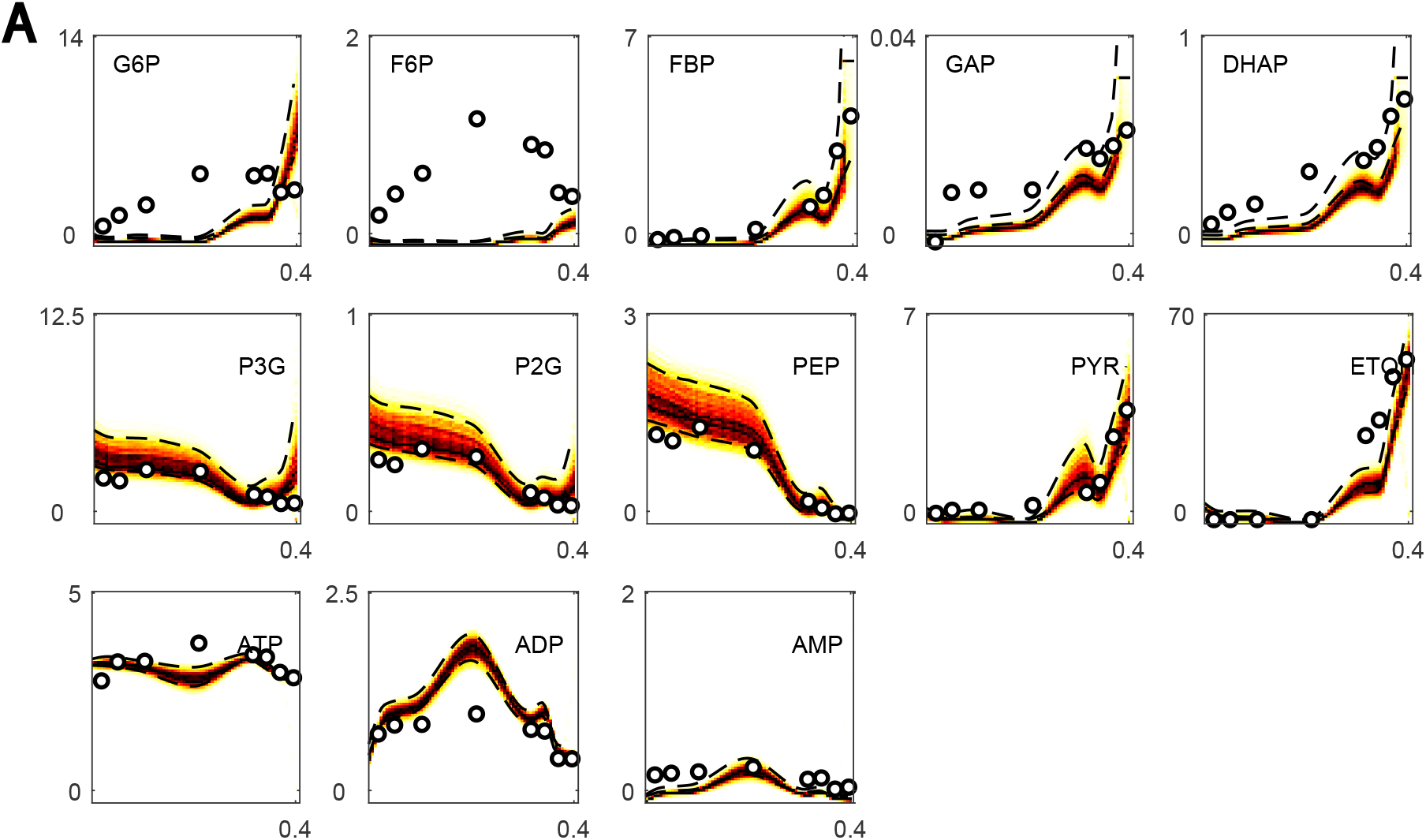
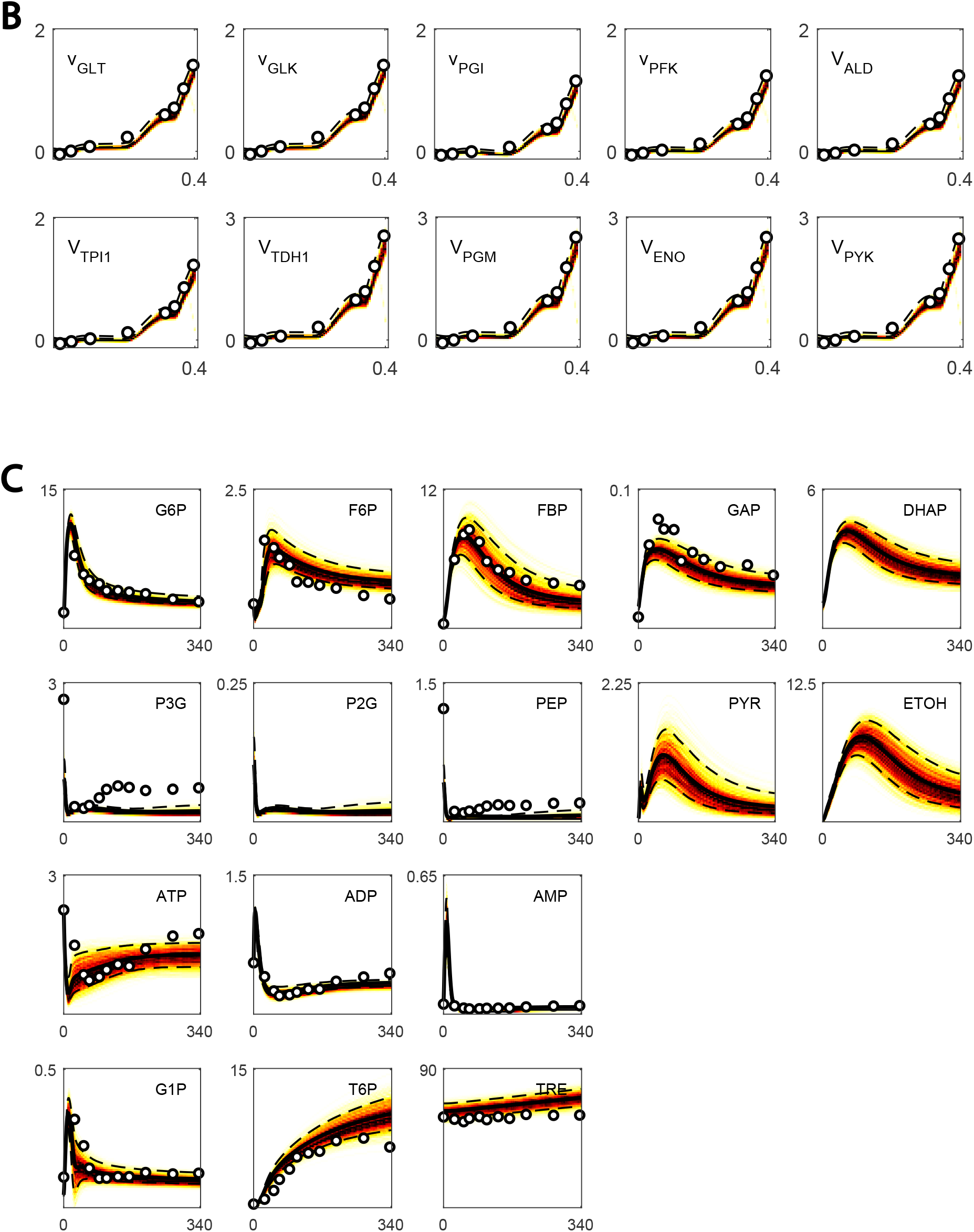
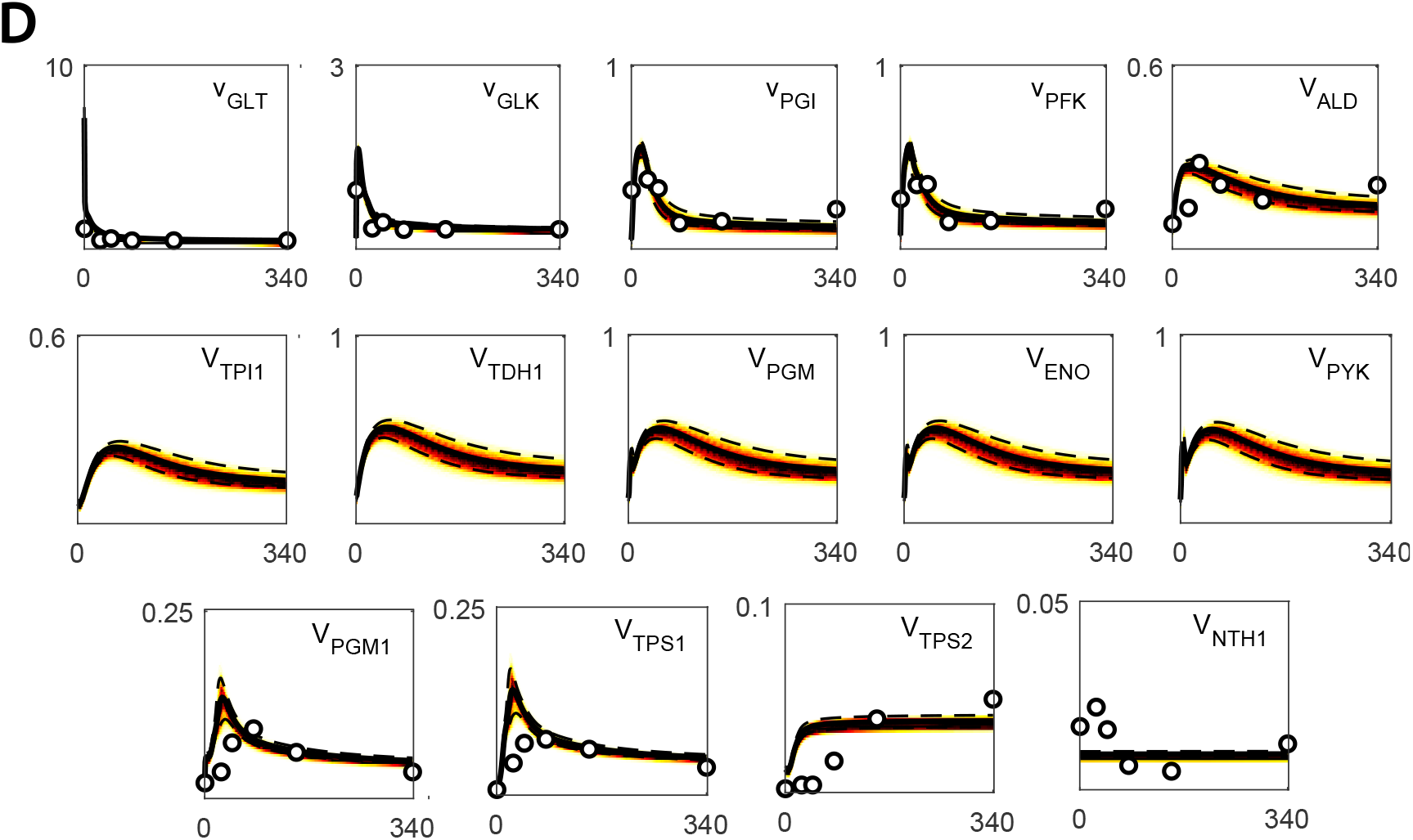
The model was robust to parameter heterogeneity. Simulations with the ensemble of models and the experimental data. Random noise was added to generate up to 1000 models in the ensemble. This noise was sampled from a random distribution within 10% of the value of each parameter. The model ensemble is shown as heat maps, where darker regions indicate higher simulation agreement. Experimental data is shown is bullet points. The semi-continuous lines indicate the region where 90% of simulations were found. (A) Steady state concentrations (mM) and dilution rate (h^-1^) in the Y. and X-axes, respectively. (B) Steady state reaction rates (mM s^-1^) and dilution rate (h^-1^) in the Y. and X-axes, respectively. (C) Glucose perturbation concentrations (mM) over time (s) in the Y. and X-axes, respectively. (D) Glucose perturbation reaction rates (mM s^-1^) over time (s) in the Y. and X-axes, respectively. To see the simulations of each metabolite and reaction rate in the model, see supplementary materials.

### Problem decomposition revealed a mismatch between literature and *in vivo* parameters

A concern in the literature is the accuracy of the estimated enzymatic kinetic constants. Previous work showed that kinetic constants measured *in vitro* in isolated enzymes assays might not be representative of *in vivo* behavior (Teusink *et al*, 2000; Davidi and Milo, 2017). This was improved by standardizing *in vivo*-like assay conditions, but only some glycolytic parameters were updated (Van Eunen *et al*, 2010; Smallbone *et al*, 2013). In this work, the abovementioned problem decomposition approach was used to assess this experimental bias and to adjust the parameters to fit the data.

Simulating the experimental steady state reaction rates with the models generated in the problem decomposition step revealed a notable bias if the parameter values found in the literature were used, even if *in vivo*-like experimental parameters were considered (Fig 6). These fits were improved using the parameter estimation pipeline described in (Fig 3). In line with (Davidi and Milo, 2017), this highlights the need of cross validating experimental data generated in individual enzyme assays with *in vivo* data and suggest that even with improved media conditions, an impactful bias exists. See the supplementary information for a complete comparison between the estimated and the literature parameter set.

**FIGURE 6.**
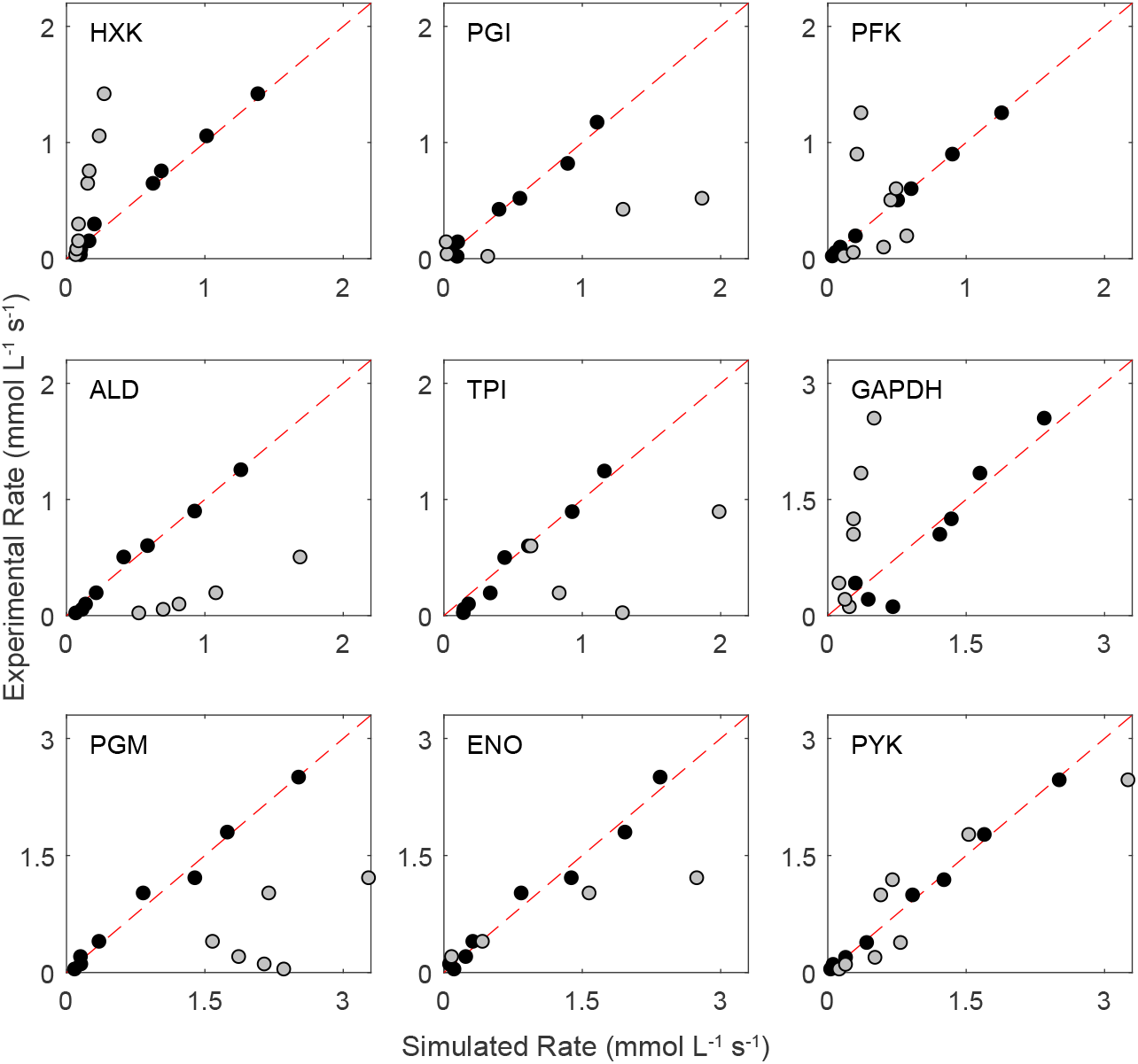
Problem decomposition reveals a mismatch between literature and *in vivo* parameters. Experimental reaction rates plotted against simulated reaction rates. Gray dots show the model fit using literature parameters. Black dots show the fit with the developed models. The red dashed line shows the location of a perfect fit. Each plot shows a different enzyme in glycolysis. Upper and lower glycolysis in top and bottom row, respectively.

### Large deviations from literature parameters suggested missing knowledge around PFK kinetics

When the estimated parameters were compared to the literature, most deviated by less than 0.5-fold from the literature value. Nonetheless, the kinetic constants which deviated the most belonged to PFK kinetics or the two enzymes found before and after this reaction in glycolysis, phosphoglucose isomerase (PGI) or aldolase (ALD) (Table 1). Furthermore, the model fit agreed for most experimental variables, except for the SS concentrations of metabolites G6P and fructose 6-phosphate (F6P), for which simulations disagreed both qualitatively and quantitatively. This could be solved if PFK kinetics parameters were changed in a growth rate dependent manner, for instance, by adding a ‘fudge factor’ to the maximum reaction rate (see supplementary information), which could suggest that there is a missing regulation on PFK kinetics. Even though well studied, the kinetics of this enzyme are known to be notable complex (Teusink *et al*, 2000; Gustavsson *et al*, 2014). For instance, the concentrations of F26bP, a modulator of PFK was not present in the data and a constant value was assumed in our simulations.

**TABLE 1.** Largest deviations between estimated and literature values are found in PGI, PFK and ALD enzymes. Only deviations above 0.75-fold change from the literature value are displayed. Fold change defined as: Fold change = estimated parameter value / literature parameter value.

| Parameter | Enzyme | Fold Change | Reference |
| --- | --- | --- | --- |
| $K_{m,DHAP}$ | ALD | -1.90 | (Teusink *et al*, 2000) |
| $K_{m,F6P}$ | PGI | 1.37 | (Smallbone *et al*, 2013) |
| $K_{AMP}$ | PFK | -0.98 | (Teusink *et al*, 2000) |
| $K_{m,G6P}$ | PGI | 0.89 | (Smallbone *et al*, 2013) |
| $K_{m,ATP}$ | PFK | 0.88 | (Teusink *et al*, 2000) |
| $K_{i,FBP}$ | PFK | 0.77 | (Teusink *et al*, 2000) |
| $K_{cat}$ | PGI | -0.76 | (van Eunen *et al*, 2012) |

### Dilution rate dependent glucose transport kinetics

Transmembrane glucose transport kinetics are known to change with dilution rate. For instance, changing affinities had to be considered to model different dilution rates (Postma *et al*, 1989). Thereafter, multiple transporters (HXT1-7) were found to be notably expressed at different growth rates (Diderich *et al*, 1999). The kinetics of these transporters were found to be divergent and isoenzyme dependent, both in terms of affinity constants (*K*_*m*_) and maximum reaction rates (*V*_*max*_) (Bosdriesz *et al*, 2018; Maier *et al*, 2002; Reifenberger *et al*, 1997). Therefore, we investigated whether a unique parameter set could explain the entire data, or dilution rate dependency had to be considered on the kinetic constants.

The simulations with the unique parameter set fit the data to a notable degree, but with certain deviation (Fig 7A). Uptake was overestimated at dilution rates below 0.2 h^-1^, resulting in fermentative excess where residual glucose concentrations remained under 0.20 mM (Fig 7B) (Postma *et al*, 1989). Then, this fit could be improved by allowing HXT *V*_*max*_ to change in a dilution rate dependent manner (Fig 7A). As expected, *V*_*max*_ was found to decrease and increase at low and high dilution rates, respectively (Fig 7C). Nonetheless, the change at low growth rates could also be explained by a decreased enzyme affinity (see supplementary information) or a combination of both *V*_*max*_ and *K*_*cat*_ changes, in line with (Diderich *et al*, 1999). Interestingly, a data point that could not be fit was the dilution rate h^-1^, given the low residual glucose concentration. As suggested in (Postma *et al*, 1989), this could be caused by intracellular changes on the driving force.

**FIGURE 7.**
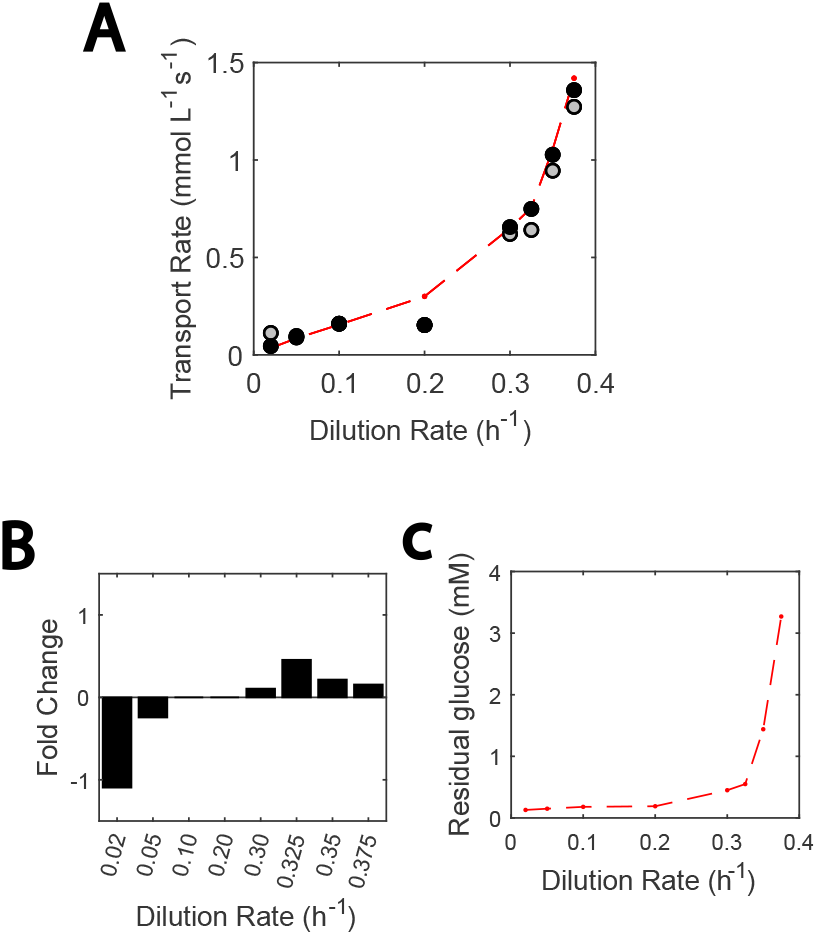
Adjustments in the HXT *V*_*max*_ can describe most growth rate-dependent transporter kinetics. (A) Reaction rates plotted against dilution rates. Gray markers show the model fit if a single set of kinetic constants for the hexose transporter is used. Black markers highlight how this fit is improved if the maximum reaction rate is fit for every growth rate. Experimental data point are shown in red markers, united by the semi-continuous red line. (B) Residual glucose concentration (mM) plotted against dilution rate, and (C) Fold change required for *V*_*max*_ at each dilution rate.

### Glycolysis was regulated by other pathways in central carbon metabolism

Glycolysis is a pathway that does not act in isolation but that is influenced by other metabolic routes in CCM. Therefore, here we investigated how the different pathways in the model regulate glycolysis. Flux towards the different glycolytic intermediate sinks was dilution rate dependent. At low growth rates, most carbon entered the sink reactions, while fermentation was predominant at higher growth rates (Fig 8A). Nonetheless, the model did overestimate fermentative flux at lower growth rate, due to small deviations of the already low glucose transport flux.

**FIGURE 8.**
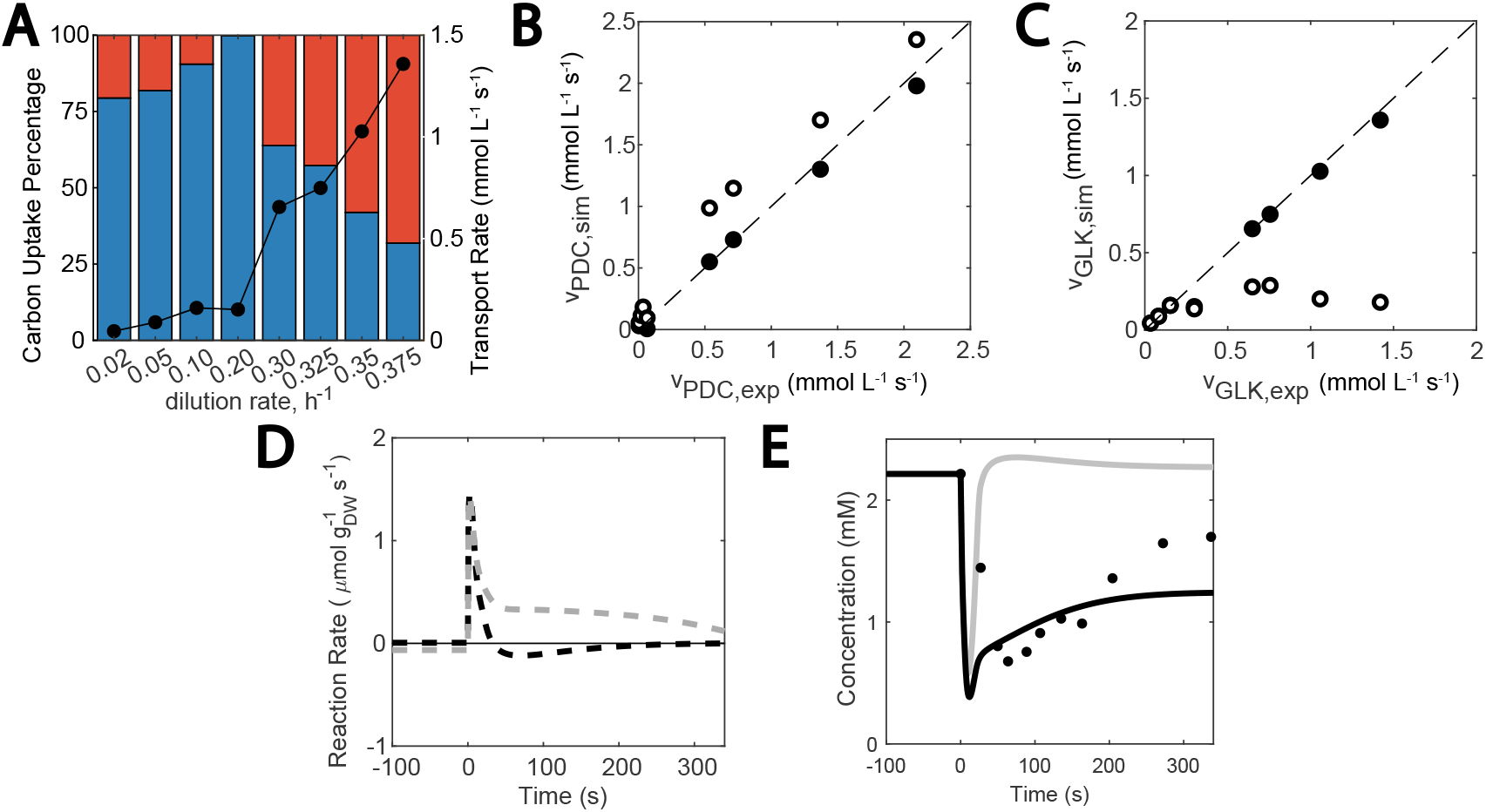
Contribution of different modules to glycolytic regulation. (A) Carbon flux distribution at different growth rates: flux towards sink reactions (blue), and towards ethanol and glycerol synthesis (red). The black dots show the simulated glucose uptake rate. (B) Effect of knocking out mitochondrial flux: Simulated PDC reaction rate plotted against experimental PDC reaction rate. Reference model (black) and effect of knockout carbon flux to respiration (white). (C) Effect of no oxygen: Simulated HXK reaction rate plotted against experimental HXK reaction rate. Reference model (black) and effect of knocking out NADH mitochondrial respiration (white). (D) Imbalance between upper and lower glycolysis (calculated as vGLK – 2*vGAPDH) occurring if the trehalose cycle is knocked out. The black and gray dashed lines show the *WT* and Trehalose cycle-knocked out strain, respectively. (E) Sensitivity of the ATP paradox to inosine salvage pathway. When the inosine salvage pathway is active (black), a decay in ATP concentrations can be observed, while knocking the cycle (gray) leads to a conservation of ATP steady state concentration.

Furthermore, the effect of knock outs resulted in expected dynamics. Simulating the lack of oxygen by knocking out mitochondrial reactions resulted in an increase in fermentation and less NADH mitochondrial respiration lowered glycolytic intake, since the network compensated by over-activation of the glycerol branch (Fig 8B and 8C). Then, knocking out the trehalose cycle during glucose perturbation resulted in an imbalance between upper and lower glycolysis reaction rates, in line with (van Heerden *et al*, 2014) (Fig 8D) and a knock out of the inosine salvage pathway meant that no ATP paradox would be observed (Fig 8E).

### Missing regulation could explain glycolytic response under a 20 g L-1 glucose pulse

At this point, we investigated how our model could simultaneously explain the response to a strong glucose perturbation of a 20 g L^-1^ glucose pulse experiment (van Heerden *et al*, 2014), the moderate perturbations of 0.4 g L^-1^ from FF experiments (Suarez-Mendez *et al*, 2017), and the steady state data from (Canelas *et al*, 2011).

We identified two type of dynamics depending on if PFK and ALD reaction rates balanced each other (Fig 9A). By analyzing the experimental reaction rate data, we could see that during the moderate glucose perturbations in (Suarez-Mendez *et al*, 2017), the glycolytic response of PFK and ALD occurred hand in hand, resulting in a transient increase of ATP (Fig 9B). Meanwhile, during the stronger perturbation in (van Heerden *et al*, 2014) the response of ALD was slower (Fig 9C), leading to more ATP being consumed in upper glycolysis than recycled in lower glycolysis, and a consequent temporary decrease in ATP.

**FIGURE 9.**
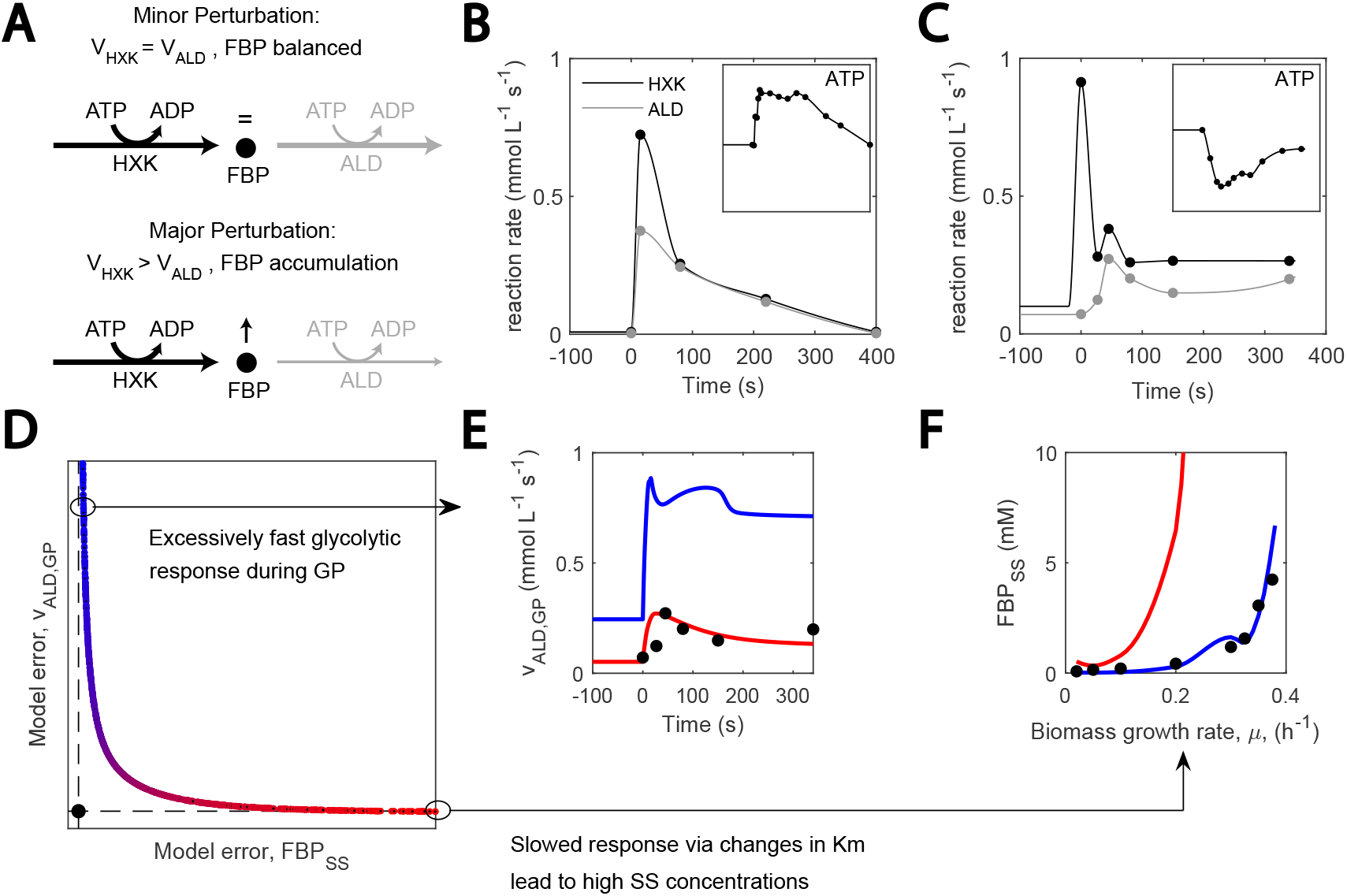
A transient imbalance between ALB and FBP can be explained by missing regulation. (A) A large glucose perturbation triggers FBP accumulation, (B-C) HXK and ALD reaction rate and ATP concentration upon glucose perturbation under feast famine regime, 0.4 mM (Suárez-Mendez et al., 2014), and single, 110 mM glucose perturbation (van Heerden et al., 2014), (D) Pareto front observed when fitting ALD reaction rate during GP and FBP SS concentrations. Red and blue colors show the zone with lower model error for ALD and FBP, respectively. (E-F) Model fits for ALD reaction rate during GP and FBP SS concentrations, respectively. Colors correspond with (D).

To understand what differed between the two conditions, we modelled both and found that they could not be simultaneously understood with a single parameter set. A pareto analysis was performed to find out which parameters needed to change to fit one and the other dataset. This resulted in the pareto front described in (Fig 9D). If the steady state fructose 1,6-bisPhosphate (FBP) concentrations were reproduced, ALD flux rapidly increased in response to the glucose perturbation. Another solution along the Pareto front was obtained for which the increase in ALD flux was slower, resembling experimental data, but then, the fit for FBP was not achieved (Fig 9E and 9F, respectively). Interestingly, this slower response resulted from apparent substrate affinity constants in upper glycolysis strongly deviating from the literature value and de facto caused enzymes to require a higher buildup of substrate before carrying out their reaction.

Altogether, this could imply that a mechanism is missing that regulates glycolytic response under strong perturbations by, for instance, transiently decreasing ALD reaction rates. Since parameter values are closer to the literature when the SS data is fit, which also results in a fast ALD response during FF, this seems to suggest that our model is best suited to fit physiology from SS and FF regime.

## Discussion

How cells respond and adapt to environmental changes while balancing internal needs prevails as a fundamental question in biology. Mechanistic models based on enzyme kinetics are important to cope with the complexity of the underlying molecular circuitry. For these models to be useful, however, they require a proper intracellular embedding of central carbon metabolism in the context of the larger metabolic system and should not be studies in isolation.

Hence, a physiologically informed kinetic model of this pathway was developed. It contained detailed descriptions of the enzymatic mechanisms that compose glycolysis, trehalose cycle and glycerol branch and extended this knowledge with coarse-grained and growth rate-dependent phenomenological descriptions of glycolytic intermediate demand, gas exchange, mitochondrial activity and ATPase activity (Fig 1). This more was then used to characterize metabolic dynamics under both a 110 mM glucose pulse and at different growth rates at steady state (Fig 2). Smaller models exist for *Escherichia coli* and *Penicillium chrysogenum* metabolic models (Chassagnole *et al*, 2002; Tang *et al*, 2017), but to our knowledge this is the first time a kinetic CCM model can realistically describe a growing *S. cerevisiae* cell. Therefore, new circumstances are now open for exploration with a unique model. For instance, by constraining the model to anaerobic conditions or by studying the shift between respiratory and respirofermentative metabolism and glucose perturbations at different growth rates (Wiebe *et al*, 2008; Ishtar Snoek and Yde Steensma, 2007; van Eunen *et al*, 2012).

One of the core developments of this work is a parameter estimation pipeline suited to identify complex systems and using large data sets (Fig 3). This approach made use of different resources: (i) divide and conquer model decomposition resulted in smaller scale systems easier to identify that the full system, (ii) global sampling determined whether multiple local minima existed, (iii) regularization improved convergence when parameters were still underdetermined and (iv) leveraging the weights of the components of the cost function to avoid overfitting. As a result, this method increases parameter identifiability, and we believe that it is well-suited for implementation in other cell factory models. Compared to other state of the art parameter estimation methods (Gábor and Banga, 2015; Villaverde *et al*, 2019; Smallbone *et al*, 2013; Raue *et al*, 2015; Steiert *et al*, 2016), this pipeline makes use of the divide and conquer approach, combined with regularization for parameters where physiological values are known. Nonetheless, limitations of this approach are the lack of reliable parameter estimates confidence intervals, given that the regularization bias added is affected by experimental noise, and the need for leveraging the weights of these data in the cost function.

In order to develop this complex and physiologically representative model, multiple data types needed to be combined. A collection of datasets containing metabolomic, fluxomic and proteomic data at steady state and during glucose perturbations was considered to make the complete model identifiable (van Hoek *et al*, 2000; Canelas *et al*, 2011; van Heerden *et al*, 2014). Considering this data was required and increased the accuracy of the parameter estimates (Fig 4). We therefore suggest making use of growingly available proteomics data to constrain the parameter solution space, as it is the case for GSM (Chan *et al*, 2017; Sánchez *et al*, 2017; Elsemman *et al*, 2022).

Ensemble modelling suggested that the network dynamics are robust to parameter heterogeneity. A modelling approach based on Monte Carlo sampling of parameter values was used to represent the heterogeneity in a cell population. This showed how dynamics and steady states were resilient to parameter value deviations (Fig 5). Assays such as this start with the premise that phenotypes arise from model structure and are performed to represent population heterogeneity (Gunawardena, 2010; Oguz *et al*, 2017; Nijhout *et al*, 2019). Therefore, given the heterogeneity in the industrial fermenter (Haringa *et al*, 2016), we propose that this ensemble modelling perspective is a more physiologically realistic representation than a single parameter set. Nonetheless, one limitation of this approach is the range to which parameters are disturbed, given that cell-specific measurements in a population are very limited (Newman *et al*, 2006).

Furthermore, we advocate to estimate parameters and supplement with literature knowledge for determination of *in vivo* enzymatic kinetic constants. Enzymatic kinetic constants were traditionally estimated *in vitro* for isolated enzymes (Teusink *et al*, 2000; van Eunen *et al*, 2012; Smallbone *et al*, 2013). However, using these to simulate experimental data showed how a notable bias still exists (Fig 6). Hence, we propose to re-estimate parameters to fit the *in vivo* metabolomic and fluxomic data, in line with (Davidi and Milo, 2017), and use the *in vitro* determined constants as reference to which the cost function is regularized. The resulting improved data fit with generally little deviation from the literature parameters implies that, even though *in vitro* studies are good estimators, they need adjustment to be implemented in a full-scale model.

When estimated parameters widely deviated from the literature values or when steady state data was not properly fit, we identified uncertainty in our model that reflected possible unknown biology. The estimated parameters that deviated the most from the literature value were from PFK or surrounding enzymes PGI and ALD (Table 1). Together with the fact that steady state concentrations of G6P and F6P are only fit in the model decomposition simulations (Fig 6) and not in the complete model simulations (Fig 5), this hints at uncertainty surrounding PFK reaction kinetics, not covered in the complex regulation already considered for this reaction (Teusink *et al*, 2000; Gustavsson *et al*, 2014). Furthermore, another area where steady state data is not perfectly fit is the HXT reaction. HXT kinetics could be explained by changes in enzyme concentration and isoenzyme-specific affinity constants, in line with (Reifenberger *et al*, 1997; Diderich *et al*, 1999). Nonetheless, the consistent lack of fit at 0.2 h^-1^ points at a missing mechanism acting intracellularly (Postma *et al*, 1989).

Finally, missing regulation could explain the transient imbalance occurring between PFK and ALD upon strong glucose perturbation (Fig 9). When explaining this imbalance with the same model structure, we found that a subset of parameters had to be adjusted, leading to a lower ALD reaction rate. Nonetheless, in the industrial bioreactor, prolonged exposure to glucose concentrations higher than 100 mM as in (van Heerden *et al*, 2014) is rare (Haringa *et al*, 2017). A more realistic representation is the FF setup, in which this imbalance did not take place under moderate perturbations in FF experiments. This could hint at influence by glucose level (Suarez-Mendez *et al*, 2014) and suggest that there is a missing phenomenon in the model, which becomes active upon large glucose perturbations. We believe that this could be due to either a regulation cascade, such as post translation modifications or cAMP/PKA pathway (Tripodi *et al*, 2015; Tamaki, 2007), given that cAMP buildup was detected during perturbation with high residual concentration but not for low (Botman *et al*, 2019; Suarez-Mendez *et al*, 2014). Consequently, the synthesis of fructose-2,6-bisphosphate by the activated PKA pathway could be regulating could be regulating PFK activity. In addition, the pH decay observed during the glucose perturbation (van Heerden *et al*, 2014) could also be relevant, given the sensitivity of glycolytic enzyme constants to this variable (Van Leemputte *et al*, 2020; Luzia *et al*, 2022). Nonetheless, to further proof these hypothesis, more experimental testing would be needed.

Here, we presented a model that describes yeast glycolysis as part of the more comprehensive central carbon metabolism, and not in isolation. This model was built using data from the metabolome, fluxome and proteome; to which extent gene regulation influenced the metabolic dynamics here studied remains an open question. At this point, gradients in the industrial bioreactor can be considered thanks to the transport rates and yields included in the model (Haringa *et al*, 2016; Nadal-Rey *et al*, 2021), bringing it closer to industrial applications. We believe that complex intracellular models like the hereby presented could soon be linked to bioreactor dynamics simulations, an area that has been so far restricted to simplified cell models (Tang *et al*, 2017; Sarkizi Shams Hajian *et al*, 2020).

## Materials and methods

### Experimental data sets used to develop the model

Three experimental datasets were used in model development: (i) steady state metabolic concentrations and fluxes at different chemostat dilution rates (0.025 - 0.375 h^-1^) (Canelas *et al*, 2011), (ii) concentrations and fluxes during a single glucose perturbation of 20 g L^-1^ (van Heerden et al., 2014) and (iii) glycolytic enzymes activity also at several chemostat dilution rates (van Hoek *et al*, 2000). Data in (Canelas *et al*, 2011; van Heerden *et al*, 2014) was obtained with the haploid yeast Saccharomyces cerevisiae CEN PK 113-7D strain, while (van Hoek *et al*, 2000) used the strain DS28911.

### Model description

The kinetic metabolic model in this work included individual enzyme reaction descriptions for glycolysis, glycerol branch and trehalose cycle, whose kinetics were taken from (van Heerden *et al*, 2014), (Smallbone *et al*, 2011) and (Cronwright *et al*, 2002), respectively. The reaction of UDP-glucose phosphorylase in the trehalose cycle was lumped due to lack of experimental data. Cofactor metabolism reactions and sinks for anabolic precursors were lumped, and the sink reactions and inosine salvage pathway were adapted from (Chassagnole *et al*, 2002) and (Walther *et al*, 2010), respectively.

### System of ordinary differential equations and reaction rate equations

The model consisted of a series of ordinary differential equations representing the mass balances for each metabolite modelled in the cytosol, except glucose and inorganic phosphate, also modelled in the extracellular space and vacuole. Moiety conservations were used for the total sum of adenosine and nicotinamide adenine nucleotides (ATP + ADP + AMP and NAD + NADH, respectively) as in (Smallbone *et al*, 2013). The ATP + ADP + AMP moiety conservation was not considered during the single glucose perturbation response, where the inosine salvage pathway was included as a pool. When available, the dilution rate-dependent protein activity change (van Hoek *et al*, 2000) was considered by adjusting the *V*_*max*_.

Enzymatic reaction rate kinetics were obtained from previous works, where reversible Michaelis Menten kinetics dominated (van Heerden *et al*, 2014; Smallbone *et al*, 2011; Cronwright *et al*, 2002). Exceptions to this were hill type kinetics for pyruvate kinase and decarboxylase (van Eunen *et al*, 2012), phosphofructokinase and facilitated diffusion for membrane transport (Teusink *et al*, 2000). Allosteric regulation acted both by activation and (competitive) inhibition. Hydrolysis reactions were modelled as irreversible. Sink reactions were modelled by phenomenological expressions that closely resembled experimental data in (Canelas *et al*, 2011). Reaction rates were expressed in (mM s^-1^). For a detailed description, see supplementary information.

### Simulation setup

Simulations were performed in agreement with the experimental setup. To confirm simulation stability, the model was first simulated for 3000 seconds using the experimental concentrations at 0.1 h^-1^ dilution rate. Then, the steady states at different dilution rates were modelled in parallel simulations where the residual glucose concentration was changed to the value in (Canelas *et al*, 2011), for 3000 seconds. The trehalose cycle was not modelled in steady state. Anabolic sink reaction rates, ATP maintenance and the mitochondrial activity were adjusted in a dilution rate-dependent manner as well. For the single glucose perturbation, residual glucose was increased to 110 mM (van Heerden *et al*, 2014) and the inosine salvage pathway was made active in a step manner. This simulation lasted 340 seconds. Matlab version 9.3.0.713579 (R2017b) was used. A summary of the differences between simulating steady states at different growth rates of the 110 mM single glucose perturbation can be seen in Table 2. Detailed expplanation on these can be found in the supplementary information.

**TABLE 2.**
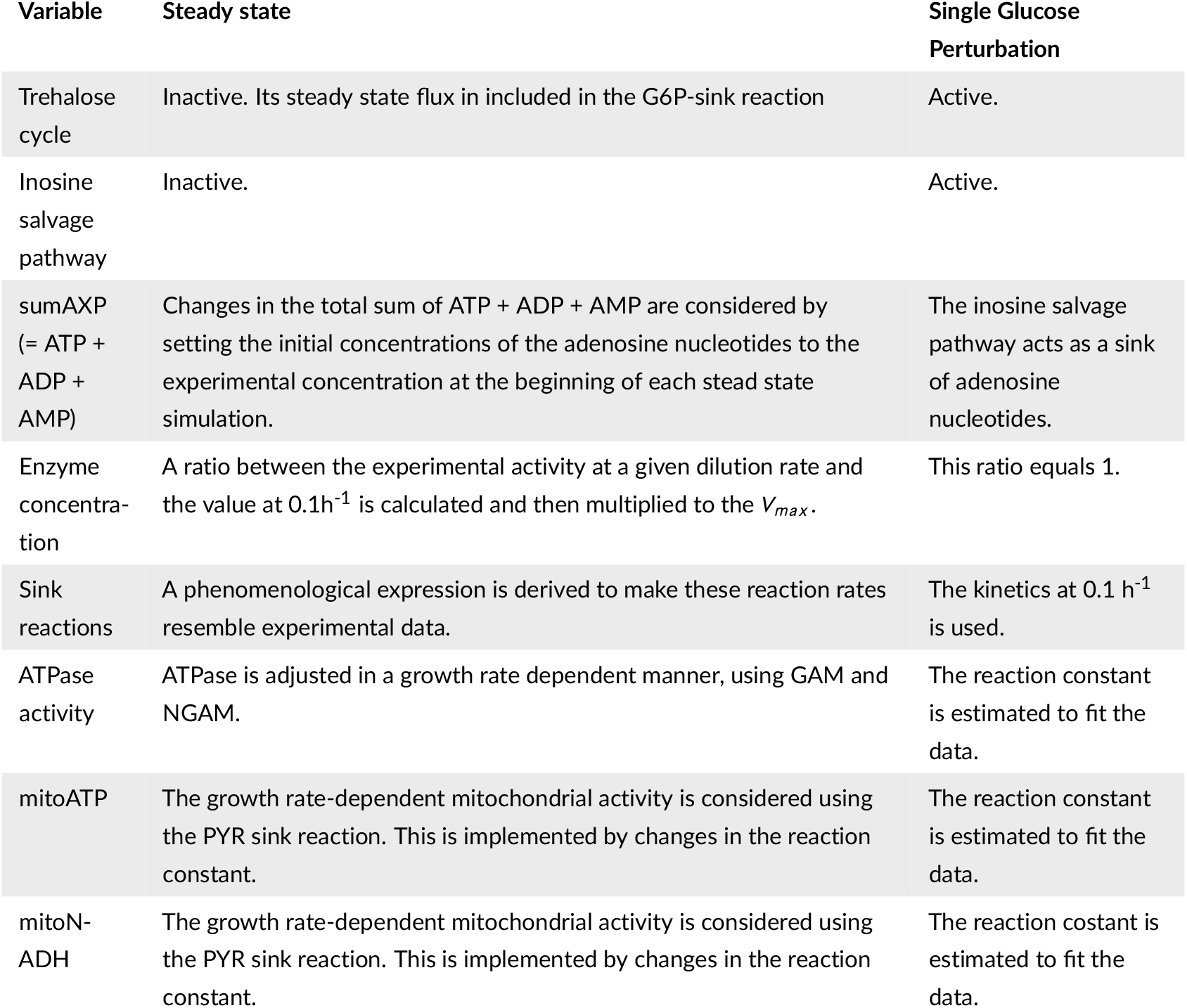
Summary of main differences in the simulation setup of the steady state and the single glucose perturbation.

### Literature parameter values

Initial estimates of kinetic constants were obtained from the literature. For glycolytic enzymes, (van Eunen *et al*, 2012) and (Smallbone *et al*, 2013) were the most recent estimates of *V*_*max*_ and *K*_*m*_. If a parameter was not available, it was taken from (Teusink *et al*, 2000). Glycerol branch and trehalose cycle parameters were retrieved from (Cronwright *et al*, 2002) and (Smallbone *et al*, 2011), respectively. For the specific parameter values used, see supplementary materials.

### In vivo parameter estimation and cost function development

Parameters in the model were estimated to fit the in vivo experimental data. The lsqnonlin solver in the Matlab optimization toolbox (version 9.3.0.713579, R2017b), which uses an interior reflective Newton method (Coleman and Li, 1996), was used to minimize the error between experimental and simulated data. To overcome the effects of ill conditioning and parameter dependencies in this large parameter set (Gábor and Banga, 2015), a model decomposition approach, also known as divide-and-conquer, was used (Kotte and Heinemann, 2009). This was first implemented to individual reactions in glycolysis. Once parameters were estimated for them, parameters were estimated for other pathways. The global solution space was explored by means of multi-start deterministic local searches (Villaverde *et al*, 2019). This model fit was supplemented with L1-type regularization (Steiert *et al*, 2016; Dolejsch *et al*, 2019), where kinetic constants were biased to resemble the experimental measurement, as long as data fit was adequate. Steady state data (Canelas *et al*, 2011) was fit first and, afterwards, single glucose perturbation (van Heerden *et al*, 2014).

## Abbreviations

[E]]: enzyme concentration
ADP: adenosine diphosphate
ALD: aldolase
AMP: adenosine monophosphate
ATP: adenoside triphosphate
CCM: central carbon metabolism
CFD: computational fluid dynamics
*CO*_2_: carbon dioxide
*E.coli*: *Escherichia coli*
ENO: enolase
F6P: fructose 6-phosphate
FF: feast famine
G6P: glucose 6-phosphate
GP: glucose pulse
GSM: genome-scale model
*K*_*cat*_: catalytic constant
*K*_*m, mAT P*_: ATP maintenance rate michaelis constant
NAD: nicotinamide-adenine-dinucleotide
*O*_2_: oxygen
ODE: ordinary differential equation
*P.chrysogenum*: *Penicillium chrysogenum*
P*/*O ratio: phosphate/oxygen ratio
PDC: pyruvate decarboxylase
PEP: phosphoenolpyruvate
PFK: phosphofructokinase
PGI: phosphoglucoisomerase
PPP: pentose phosphate pathway
PTM: post-translational modifications
*q*_*AT P*_: ATP production rate
*q*_*CO2*_: carbon dioxide transport rate
*q*_*O2*_: oxygen transport rate
RQ: respiratory quotient
*S.cerevisiae*: *Saccharomyces cerevisiae*
SS: steady state
*sum*_*AXP*_: sum of ATP, ADP and AMP
TCA cycle: Tricarboxylic acid cycle
TPS1: trehalose phosphate synthase 1
*V*_*max*_: maximum reaction rate
VVUQ: verification, validation and uncertainty quantification.

## Acknowledgements

We thank our colleagues from the Yeast 3M team, S.Aljoscha Wahl, Koen J.A. Verhagen, Laura Guilherme Luzia, Evelina Tutucci and Johann van Heerden, for the insightful discussions.

## Author contributions

**David Lao-Martil:** Conceptualization; Validation; Investigation; Methodology; Writing - originak draft; Writing - review and editing. **Joep Schmitz:** Conceptualization; Investigation; Methodology; Writing - review and editing. **Bas Teusink:** Conceptualization; Writing - review and editing. **Natal van Riel:** Conceptualization; Writing; Methodology - review and editing.

## Data, software and model availability

This work was developed in MATLAB and model implementations are also available in Python language and SBML format. The data used and model developed in this work are available in the github repository from the author (github.com/DavidLaoM/y3m1_ss_gp) and the CBio group (github.com/Computational-Biology-TUe).

