## Supplementary material for "Glycolysis revisited: from steady state growth to glucose pulses": SM_DandC_ALD.pdf

### Aldolase

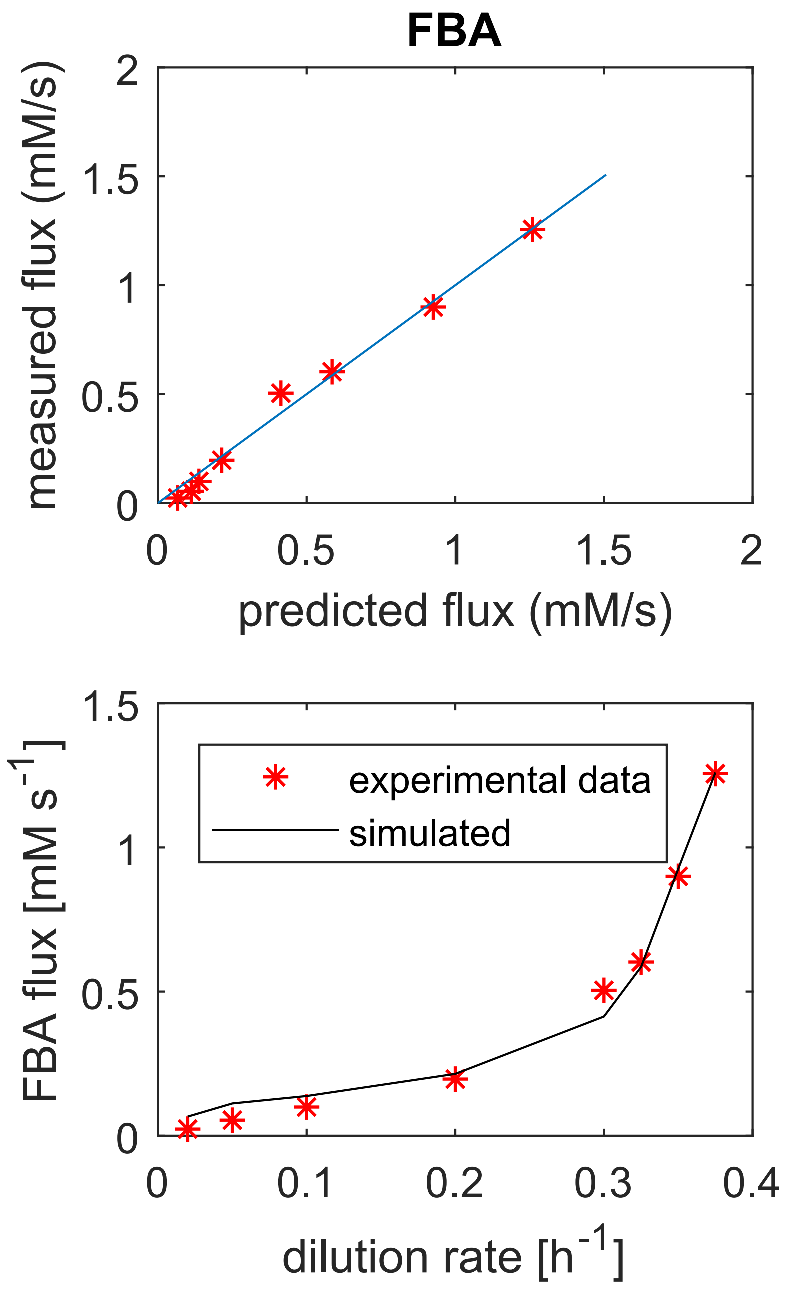


**FIGURE 1** Aldolase. Model fit.

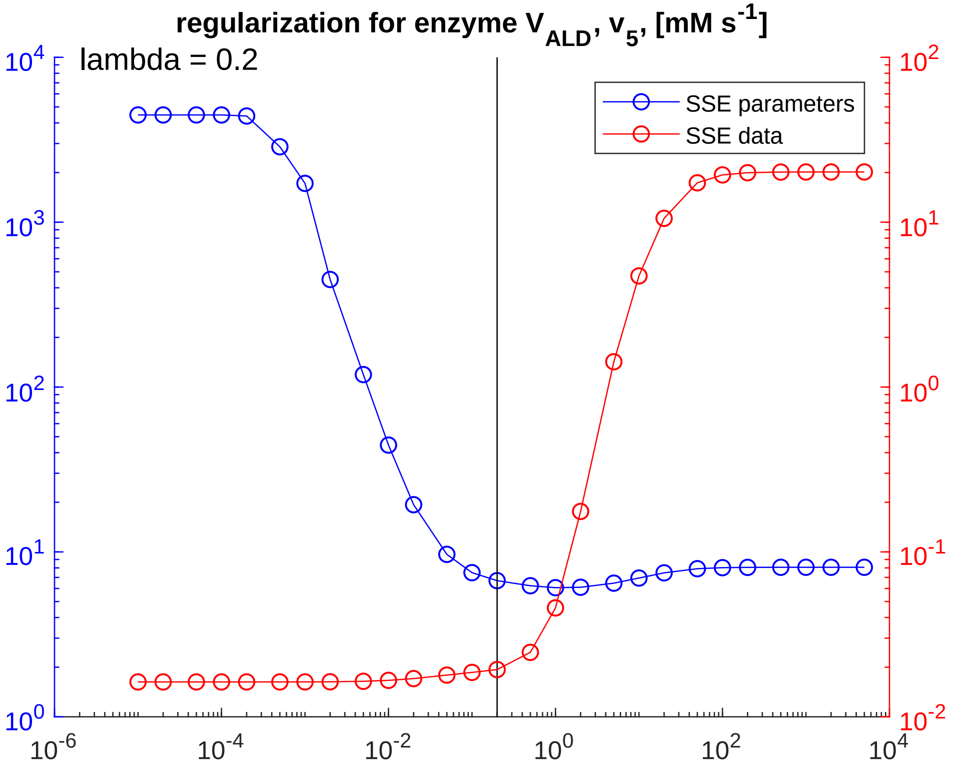


**FIGURE 2** **Aldolase. Regularization.**

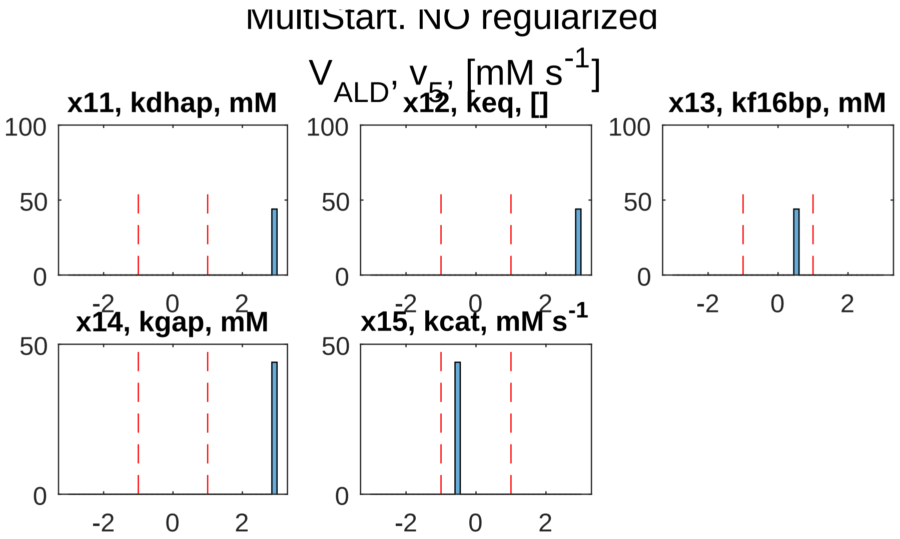


**FIGURE 3** Aldolase. Global sampling, non-regularized.

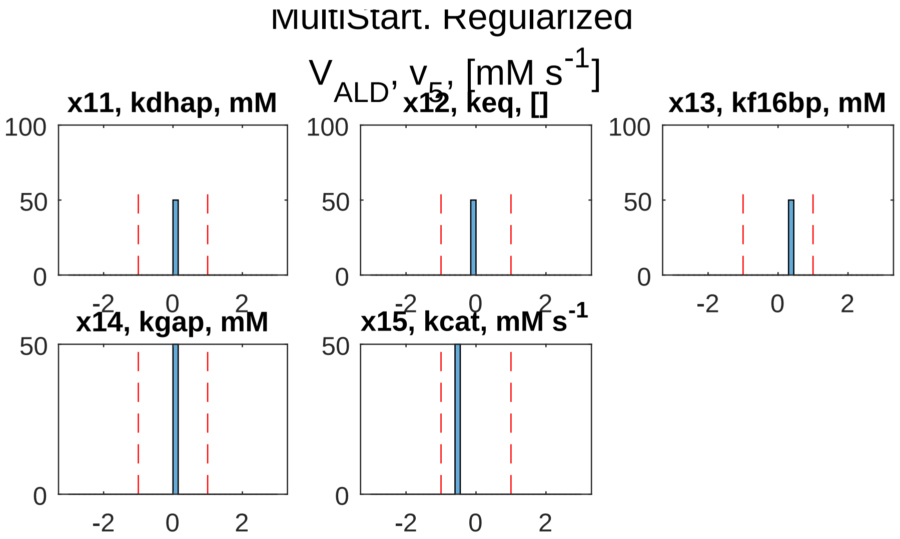


**FIGURE 4** Aldolase. Global sampling, regularized.

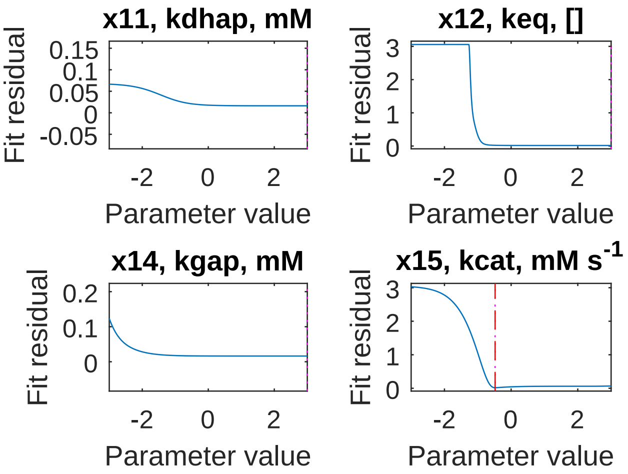


**FIGURE 5** Aldolase. PLA, non-regularized.

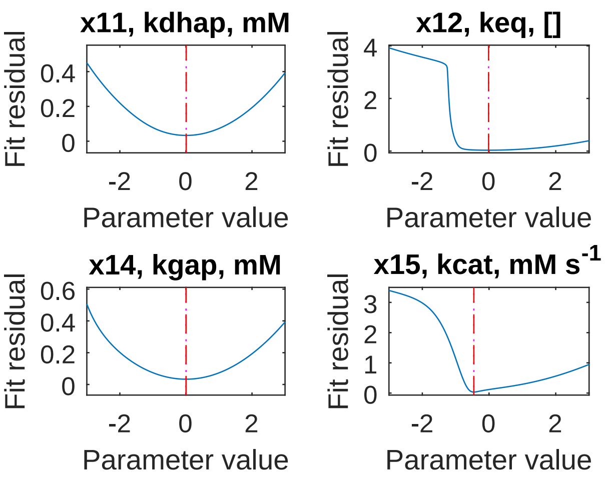


**FIGURE 6** Aldolase. PLA, regularized.
