## Supplementary material for "Glycolysis revisited: from steady state growth to glucose pulses": SM_DandC_ENO.pdf

### Enolase

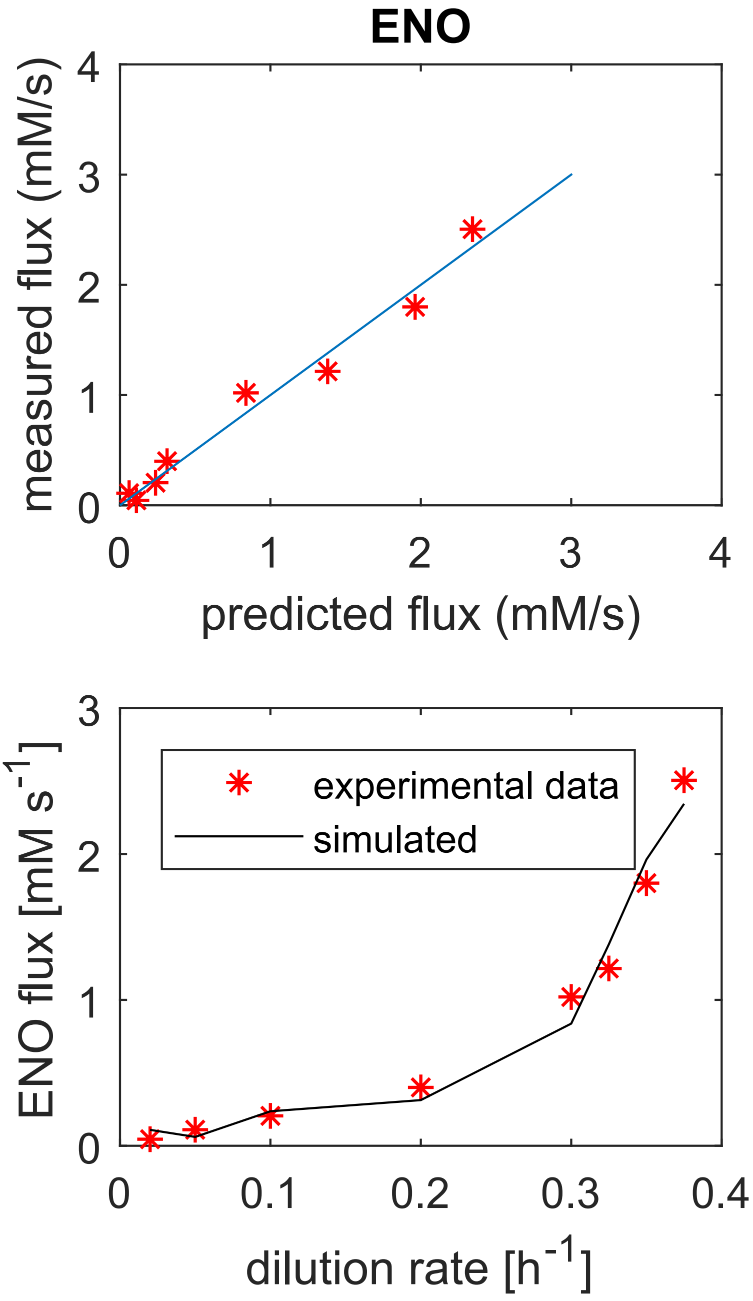


**FIGURE 1** Enolase. Model fit.

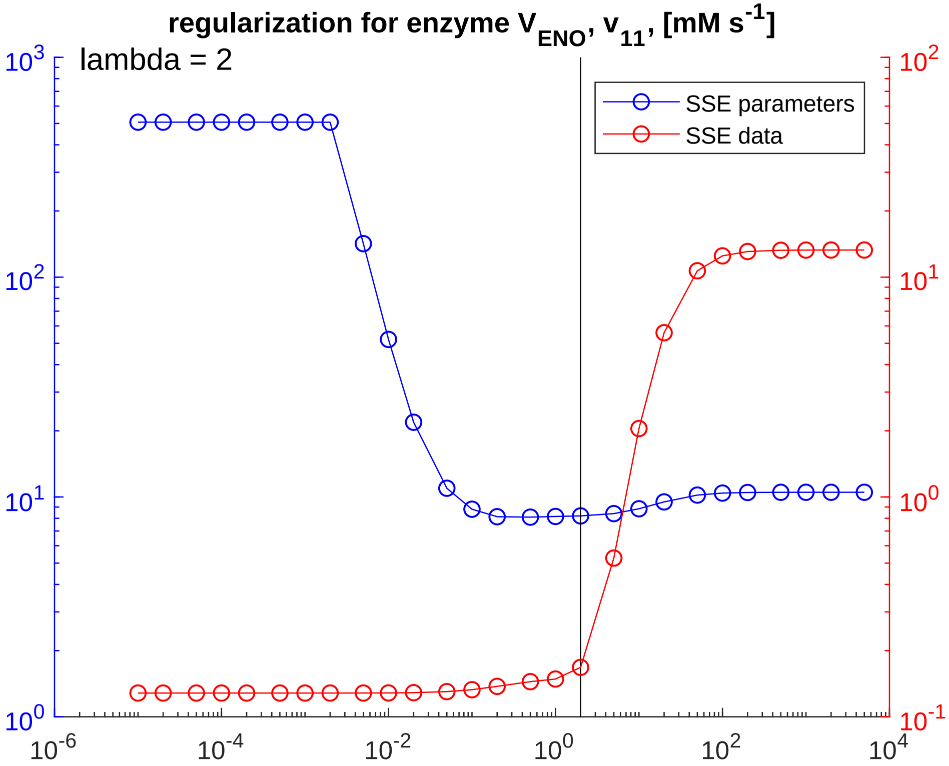


**FIGURE 2** **Enolase. Regularization.**

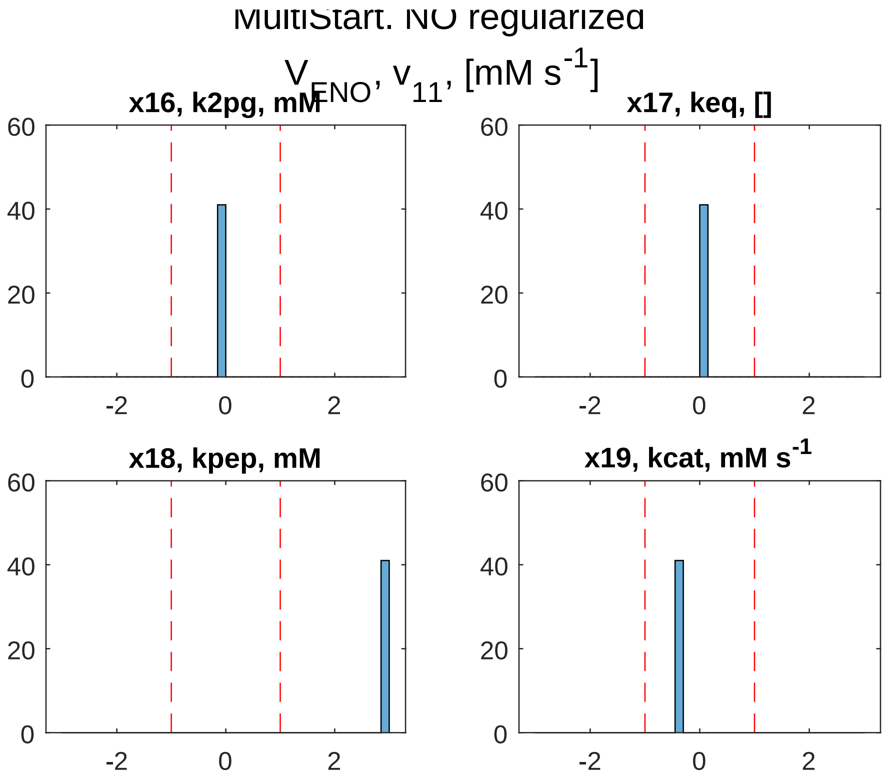


**FIGURE 3** Enolase. Global sampling, non-regularized.

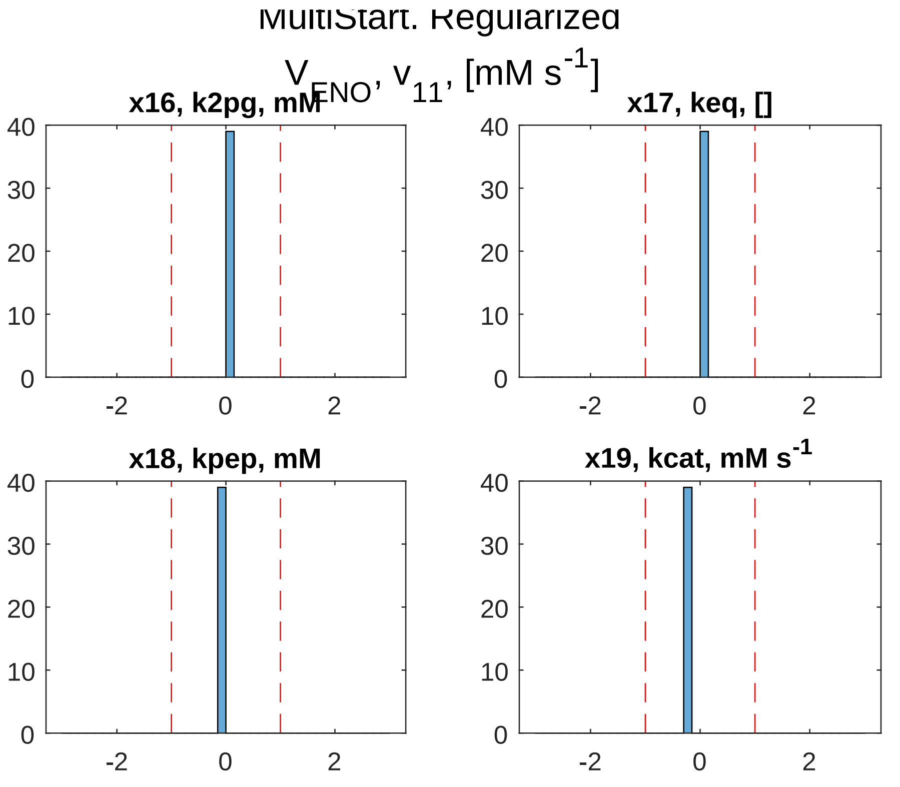


**FIGURE 4** **Enolase. Global sampling, regularized.**

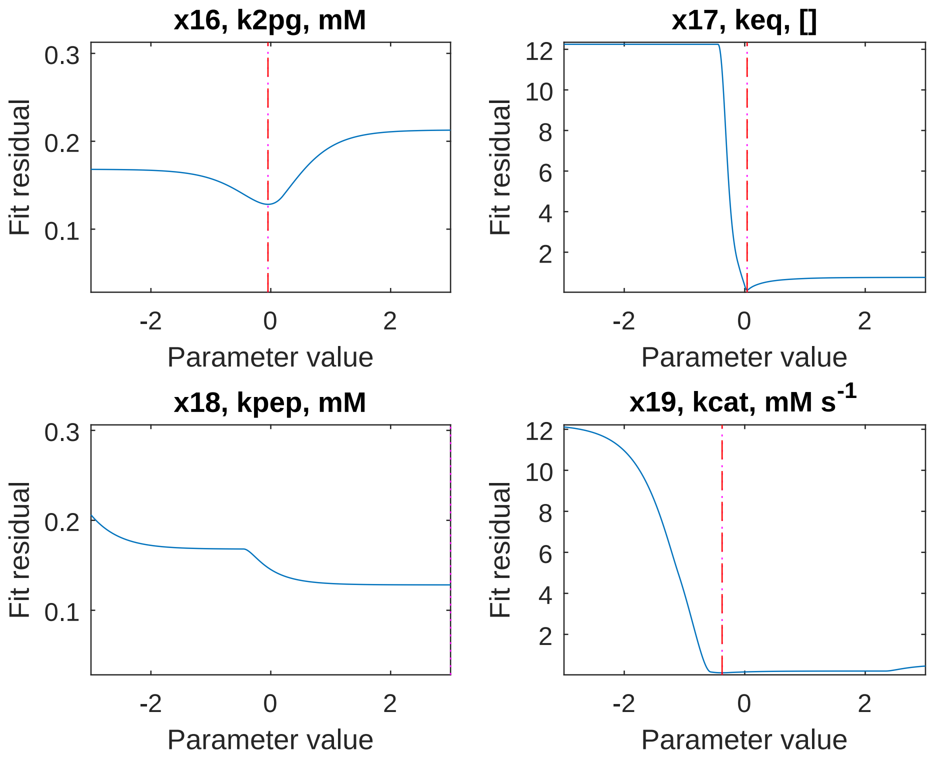


**FIGURE 5** **Enolase. PLA, non-regularized.**

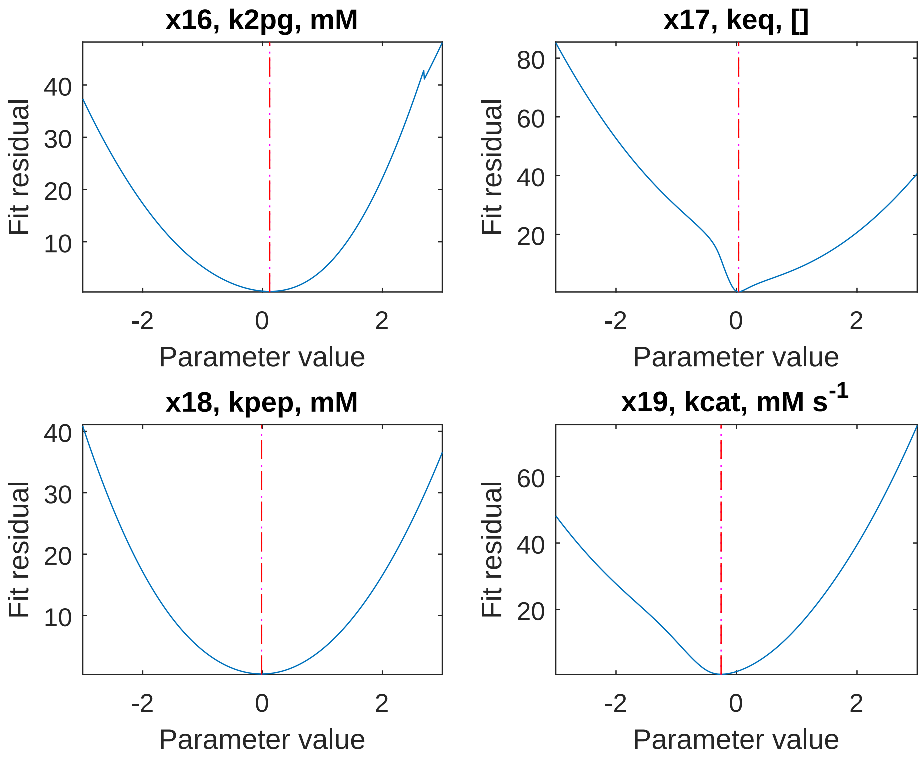


**FIGURE 6** **Enolase. PLA, regularized.**
