## Supplementary material for "Glycolysis revisited: from steady state growth to glucose pulses": SM_DandC_GAPDH_PGK.pdf

supplementary/divide_and_conquer/SM_DandC_GAPDH_PGK_SScosfunNonreg.pdf

**FIGURE 1** **GAPDH + PGK. Model fit.**

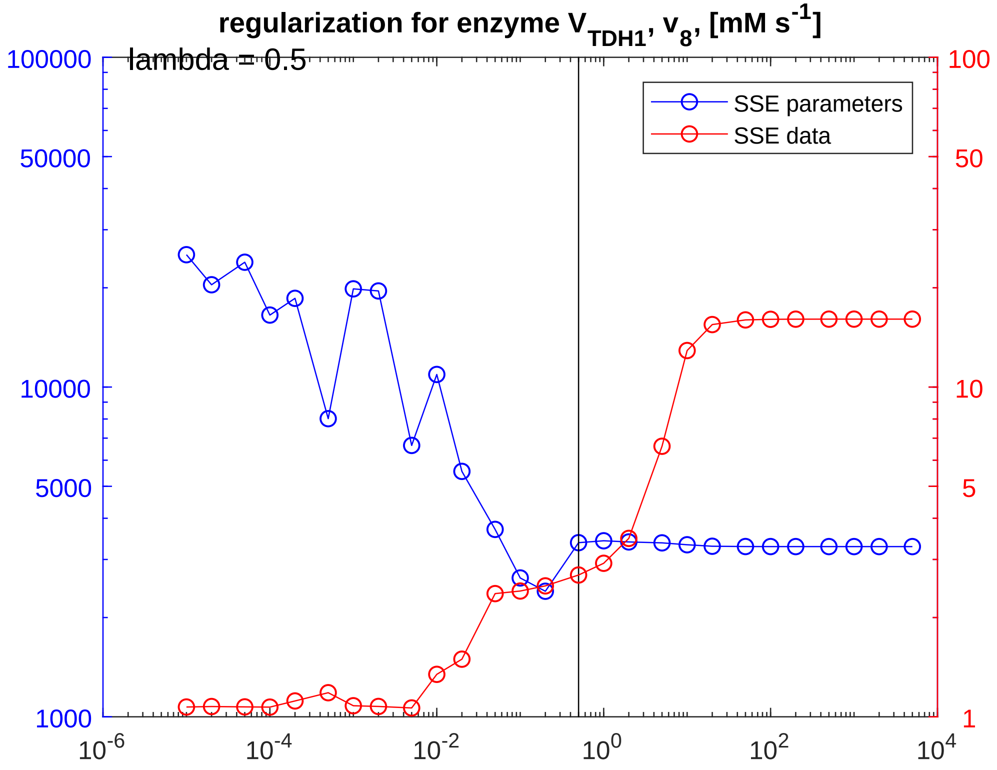


**FIGURE 2** **GAPDH + PGK. Regularization.**

supplementary/divide_and_conquer/SM_DandC_GAPDH_PGK_SSMSNonreg.pdf

**FIGURE 3** **GAPDH + PGK. Global sampling, non-regularized.**

supplementary/divide_and_conquer/SM_DandC_GAPDH_PGK_SSMSreg.pdf

**FIGURE 4** **GAPDH + PGK. Global sampling, regularized.**

supplementary/divide_and_conquer/SM_DandC_GAPDH_PGK_PLAnonreg.pdf

**FIGURE 5** **GAPDH + PGK. PLA, non-regularized.**

supplementary/divide_and_conquer/SM_DandC_GAPDH_PGK_PLAreg.pdf

**FIGURE 6** GAPDH + PGK. PLA, regularized.
