## Supplementary material for "Glycolysis revisited: from steady state growth to glucose pulses": SM_DandC_GLT_HXK.pdf

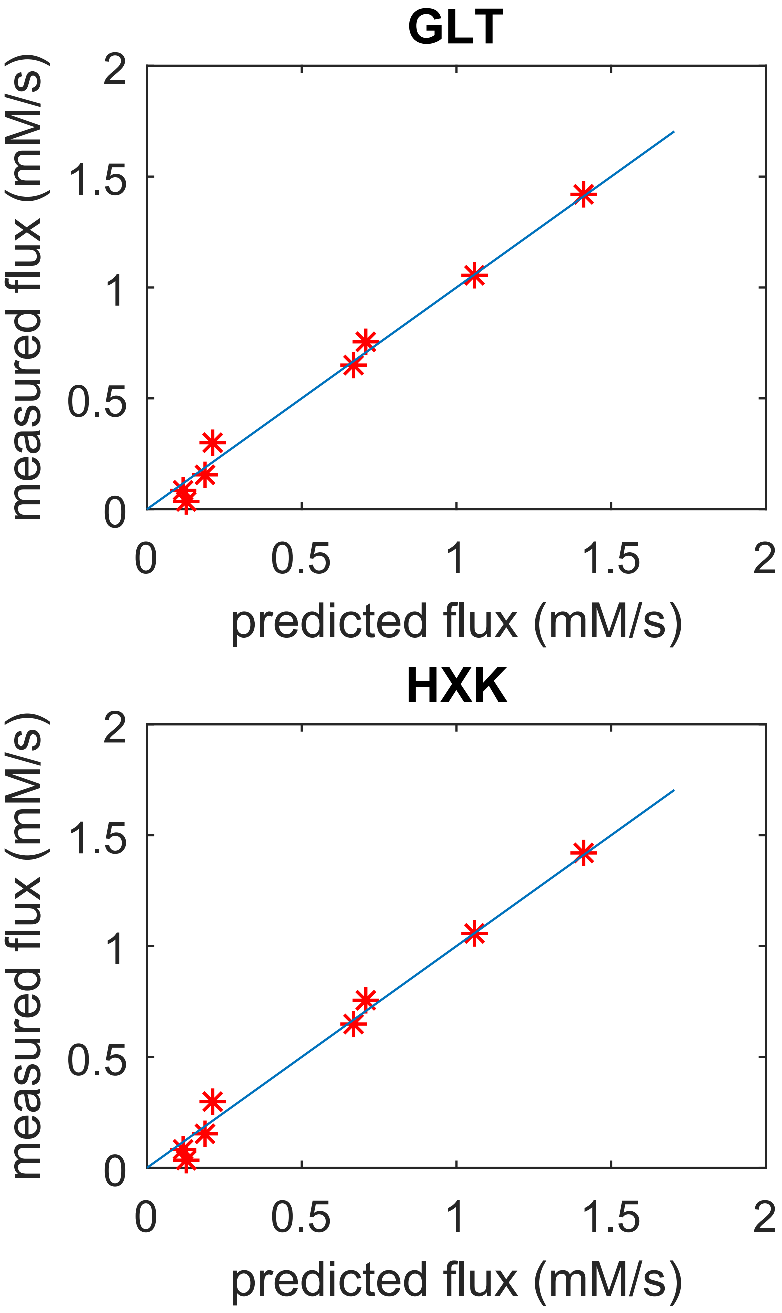


**FIGURE 1** GLT + HXK. Model fit.

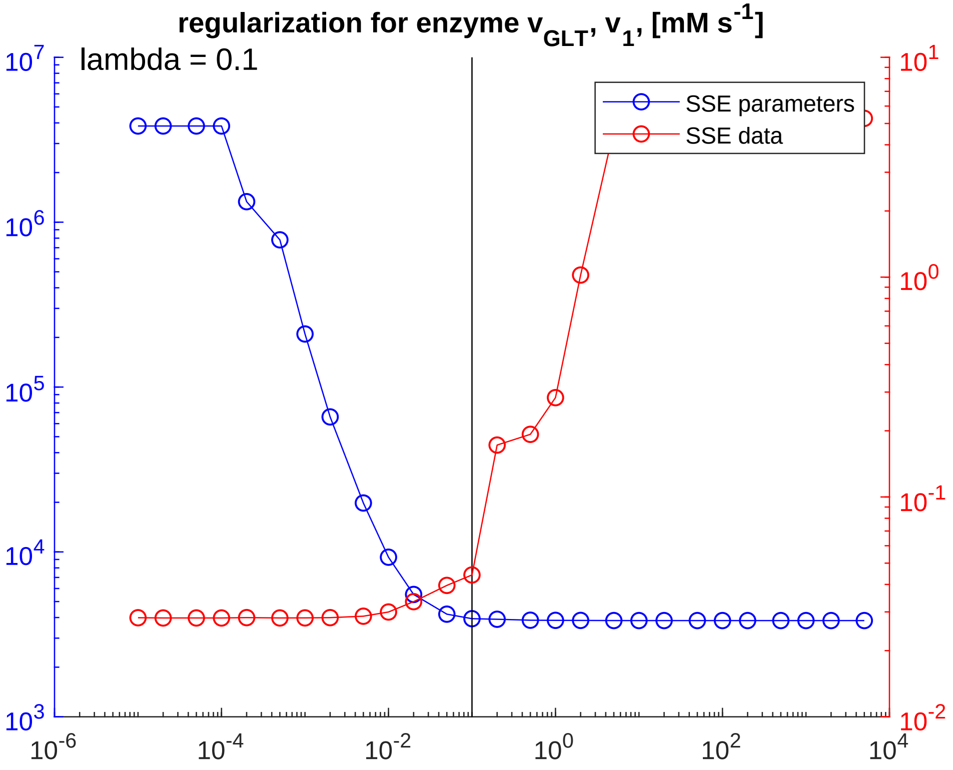


**FIGURE 2** **GLT + HXK. Regularization.**

supplementary/divide_and_conquer/SM_DandC_GLT_HXK_SSMSNonreg.pdf

**FIGURE 3 GLT + HXK. Global sampling, non-regularized.**

supplementary/divide_and_conquer/SM_DandC_GLT_HXK_SSMSreg.pdf

**FIGURE 4** **GLT + HXK. Global sampling, regularized.**

supplementary/divide_and_conquer/SM_DandC_GLT_HXK_PLAnonreg.pdf

**FIGURE 5** **GLT + HXK. PLA, non-regularized.**

supplementary/divide_and_conquer/SM_DandC_GLT_HXK_PLAreg.pdf

**FIGURE 6** **GLT + HXK. PLA, regularized.**
