## Supplementary material for "Glycolysis revisited: from steady state growth to glucose pulses": SM_DandC_GPD.pdf

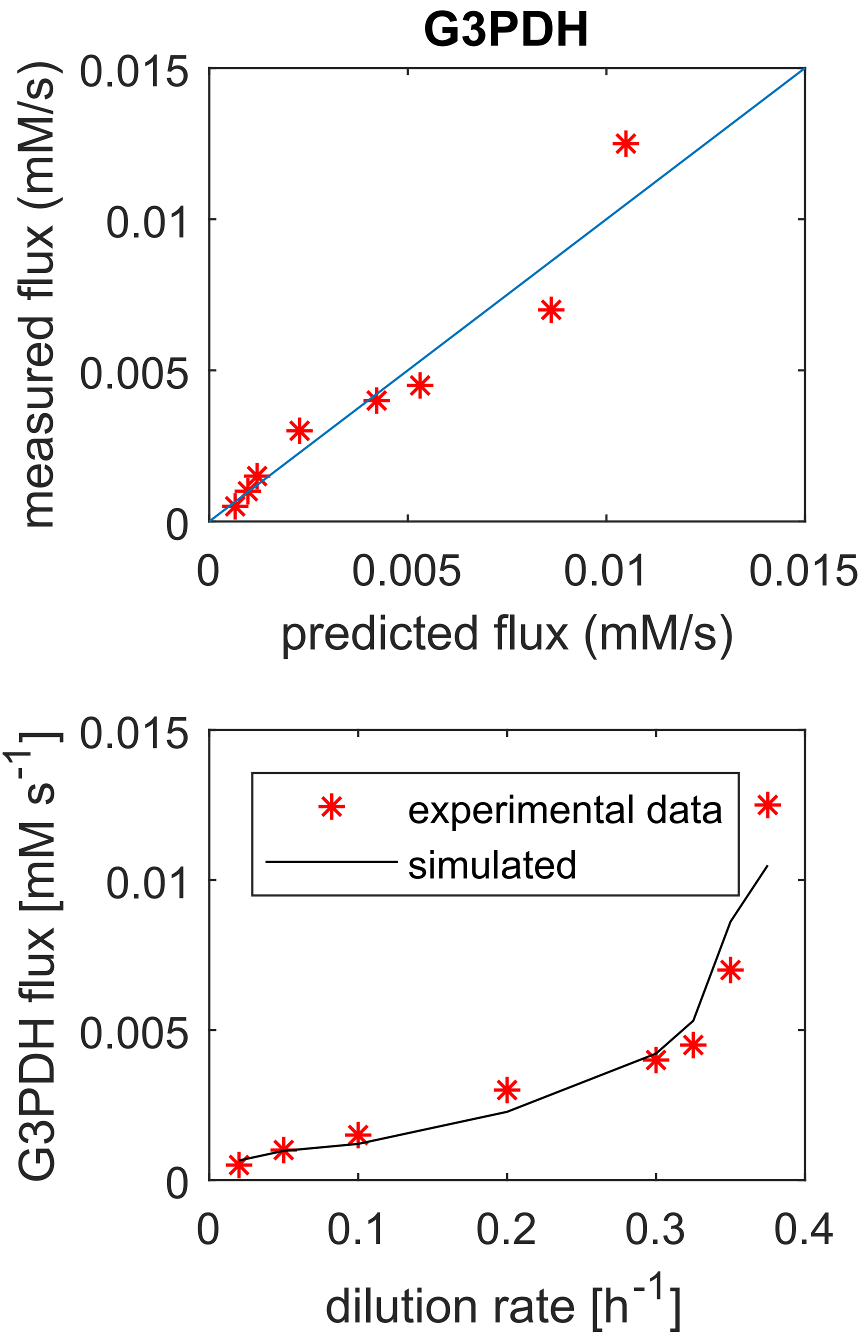


**FIGURE 1** GPD. Model fit.

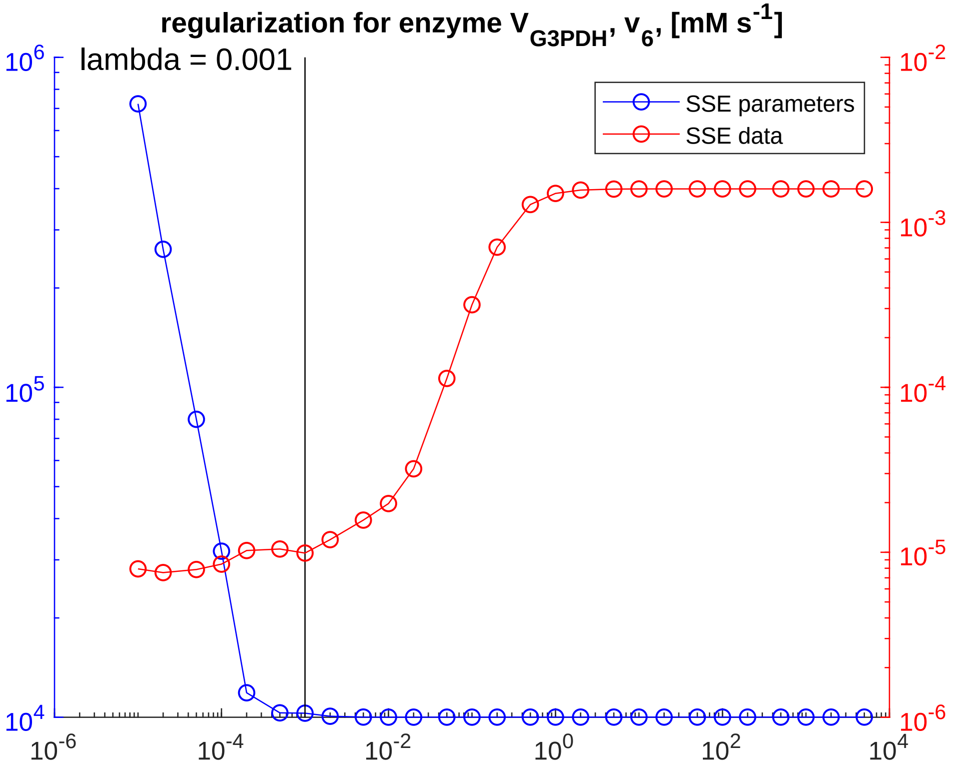


**FIGURE 2** **GPD. Regularization.**

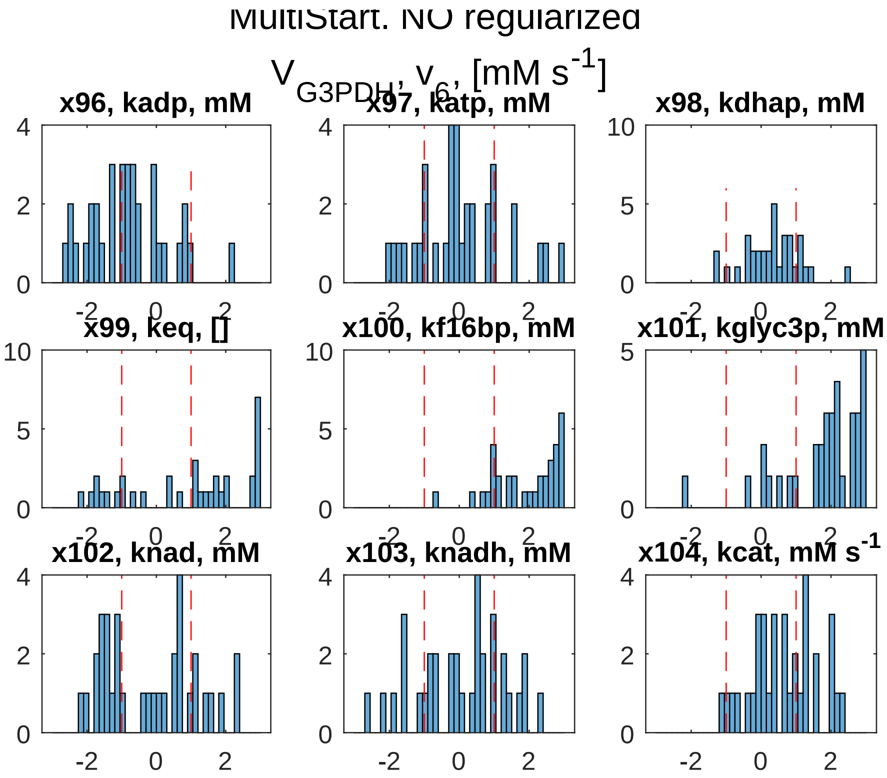


**FIGURE 3** GPD. Global sampling, non-regularized.

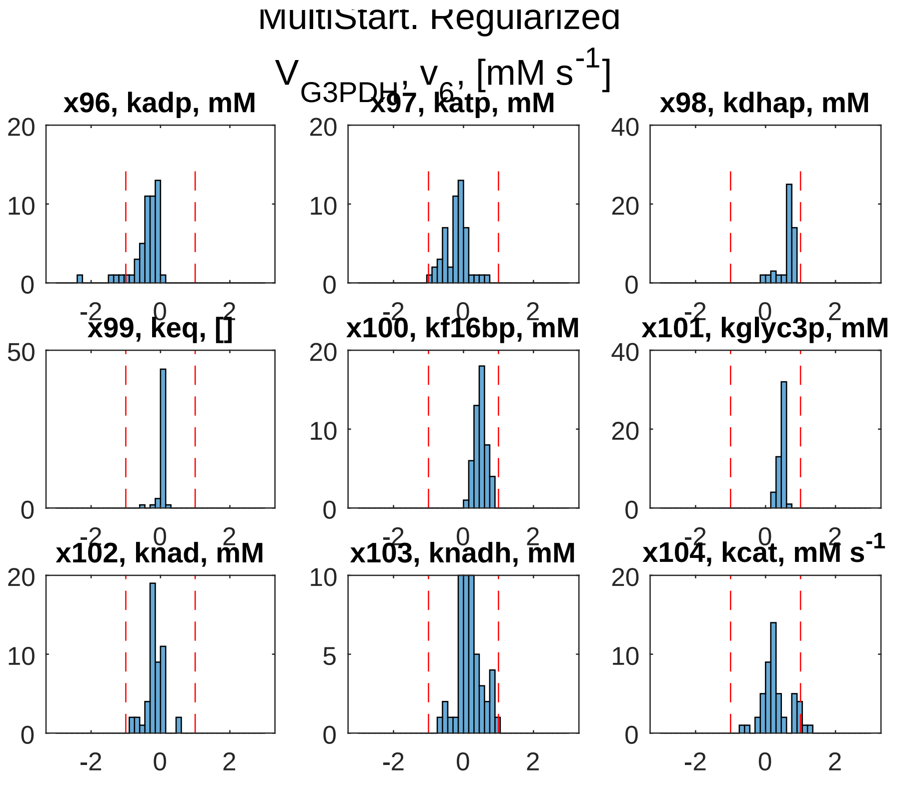


**FIGURE 4** GPD. Global sampling, regularized.

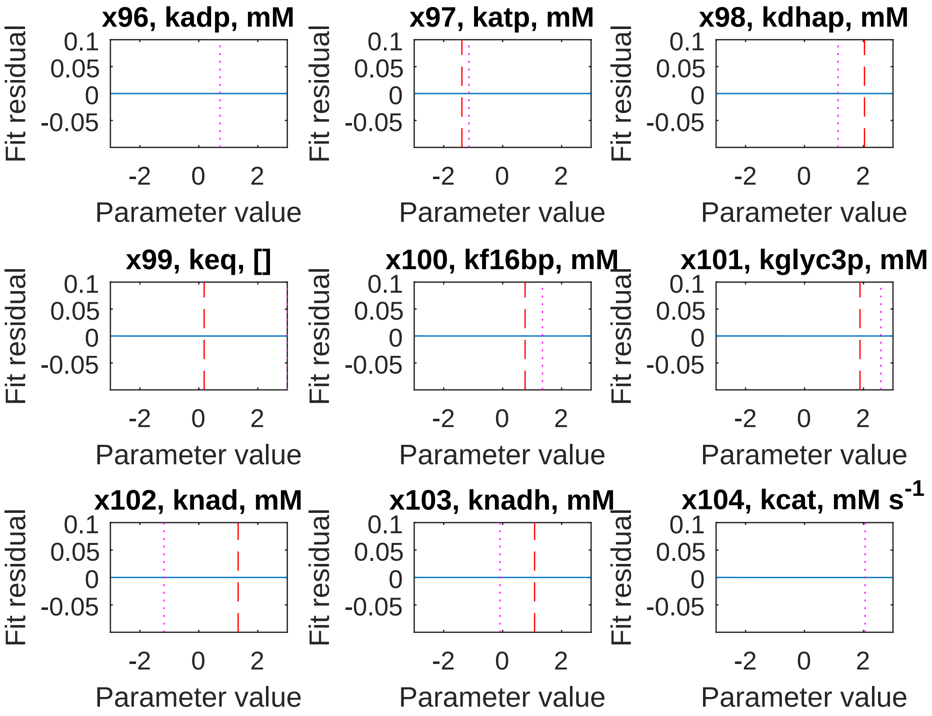


**FIGURE 5** GPD. PLA, non-regularized.

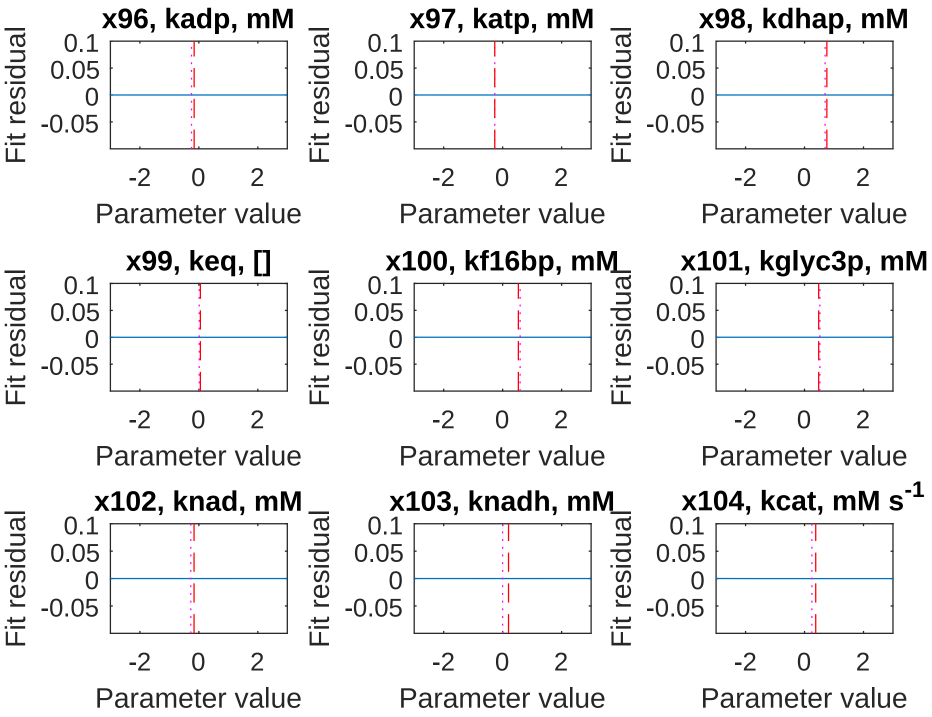


**FIGURE 6** **GPD. PLA, regularized.**
