## Supplementary material for "Glycolysis revisited: from steady state growth to glucose pulses": SM_DandC_PDC_ADH.pdf

supplementary/divide_and_conquer/SM_DandC_PDC_ADH_branches_SScosfunNonreg.pdf

**FIGURE 1** PDC + ADH. Model fit.

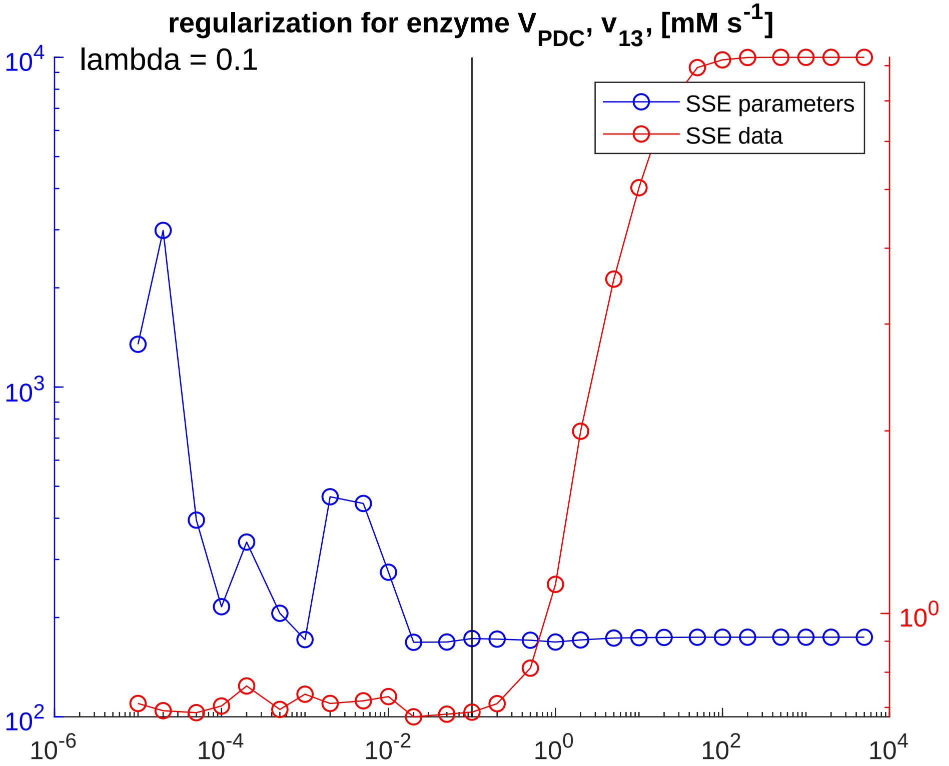


**FIGURE 2** **PDC + ADH. Regularization.**

supplementary/divide_and_conquer/SM_DandC_PDC_ADH_branches_SSMSNonreg.pdf

**FIGURE 3 PDC + ADH. Global sampling, non-regularized.**

supplementary/divide_and_conquer/SM_DandC_PDC_ADH_branches_SSMSreg.pdf

**FIGURE 4** **PDC + ADH. Global sampling, regularized.**

supplementary/divide_and_conquer/SM_DandC_PDC_ADH_branches_PLAnonreg.pdf

**FIGURE 5** **PDC + ADH. PLA, non-regularized.**

supplementary/divide_and_conquer/SM_DandC_PDC_ADH_branches_PLAreg.pdf

**FIGURE 6** **PDC + ADH. PLA, regularized.**
