## Supplementary material for "Glycolysis revisited: from steady state growth to glucose pulses": SM_DandC_PFK.pdf

### Phosphofructokinase

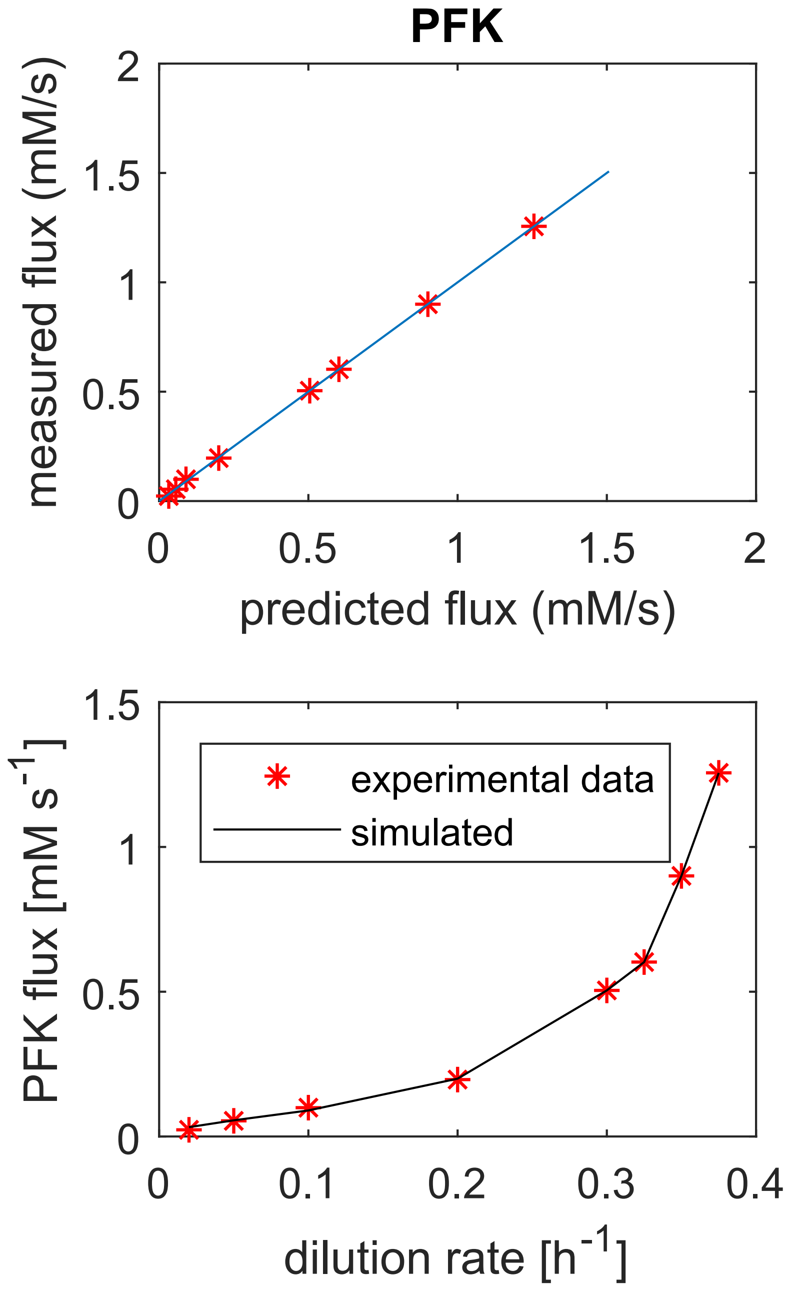


**FIGURE 1** Phosphofructokinase. Model fit.

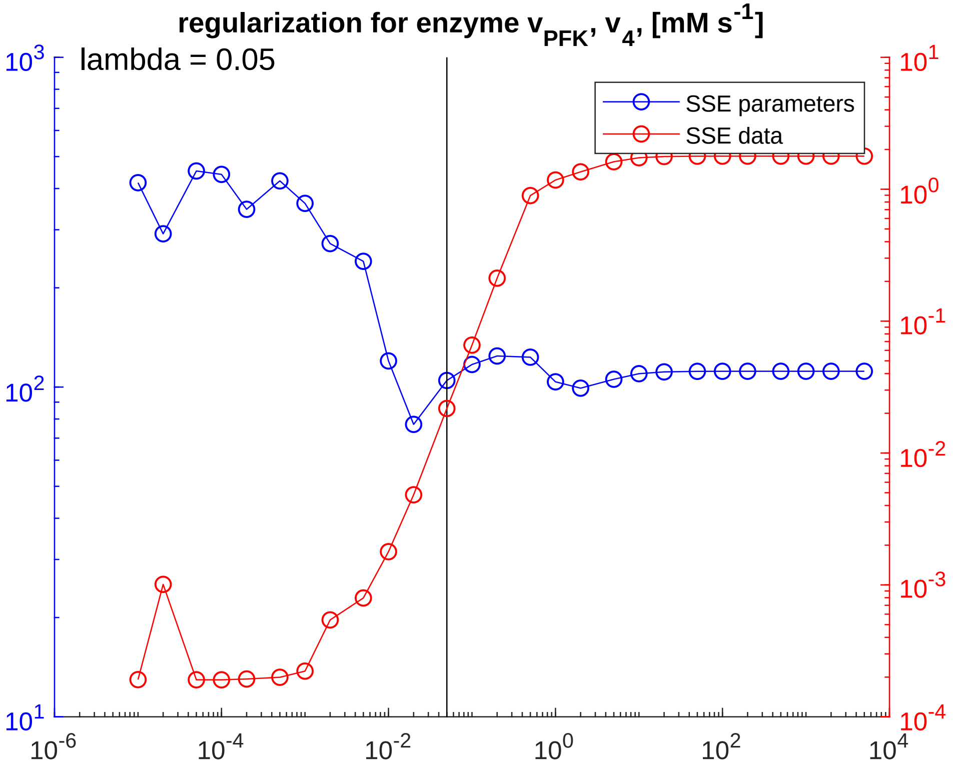


**FIGURE 2** **Phosphofructokinase. Regularization.**

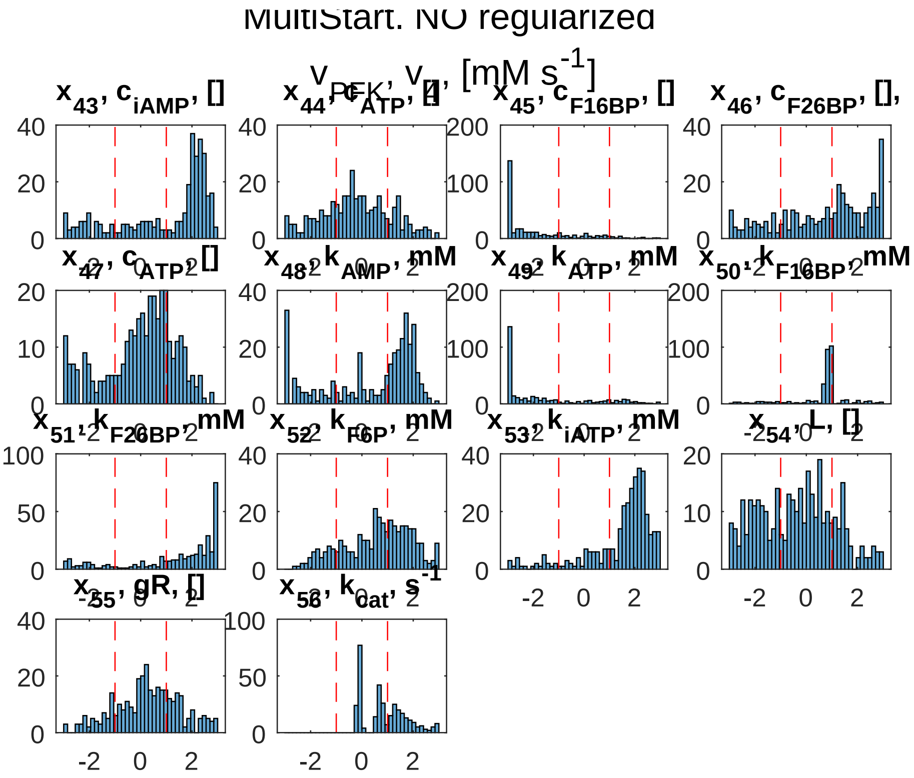


**FIGURE 3** Phosphofructokinase. Global sampling, non-regularized.

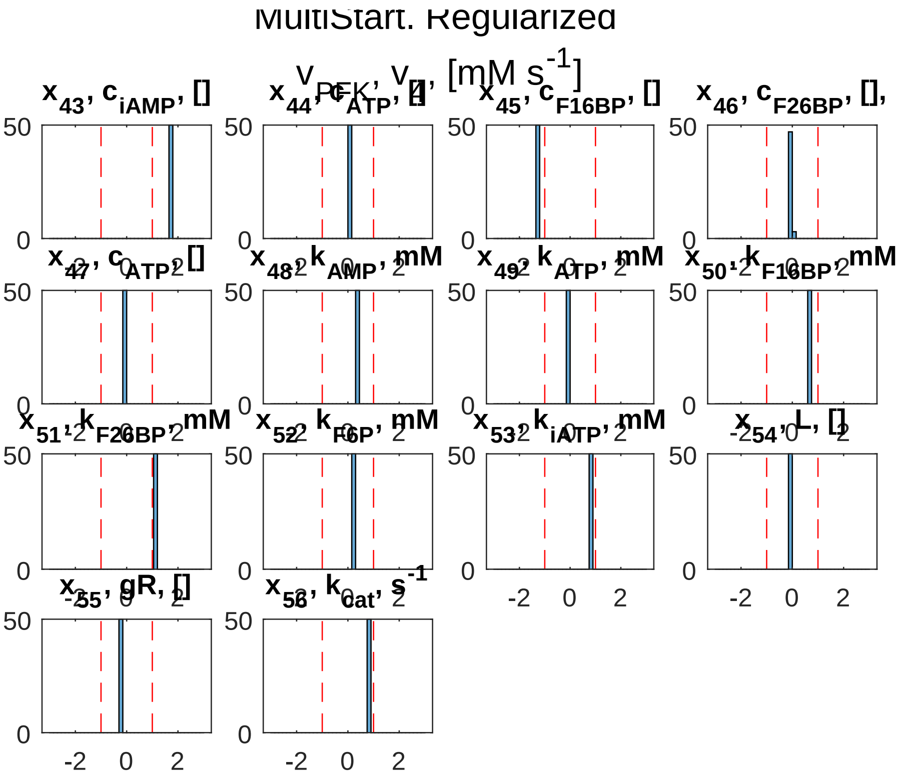


**FIGURE 4** Phosphofructokinase. Global sampling, regularized.

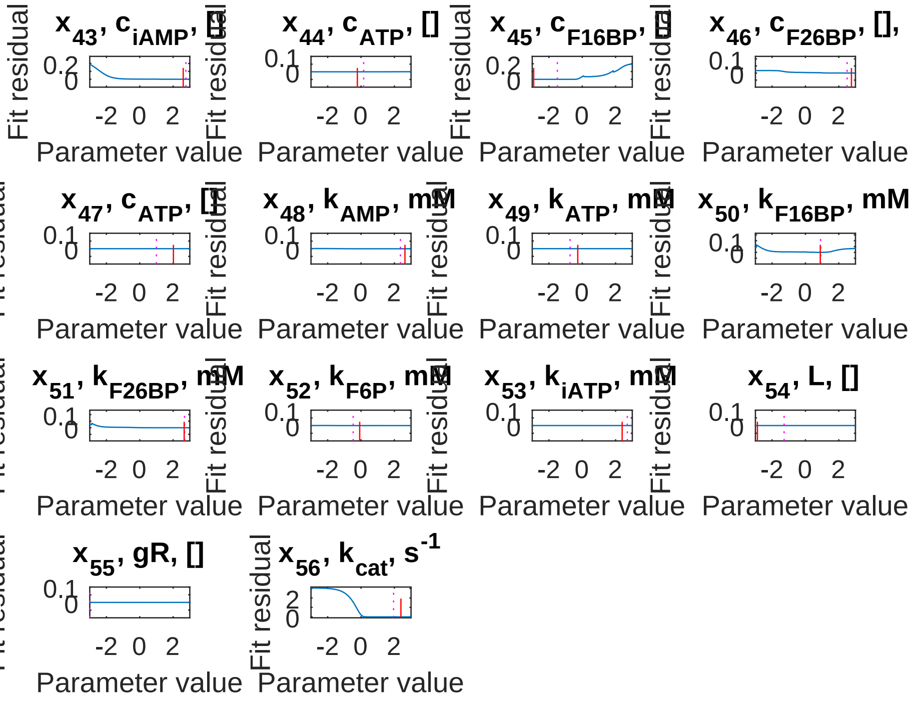


**FIGURE 5** **Phosphofructokinase. PLA, non-regularized.**

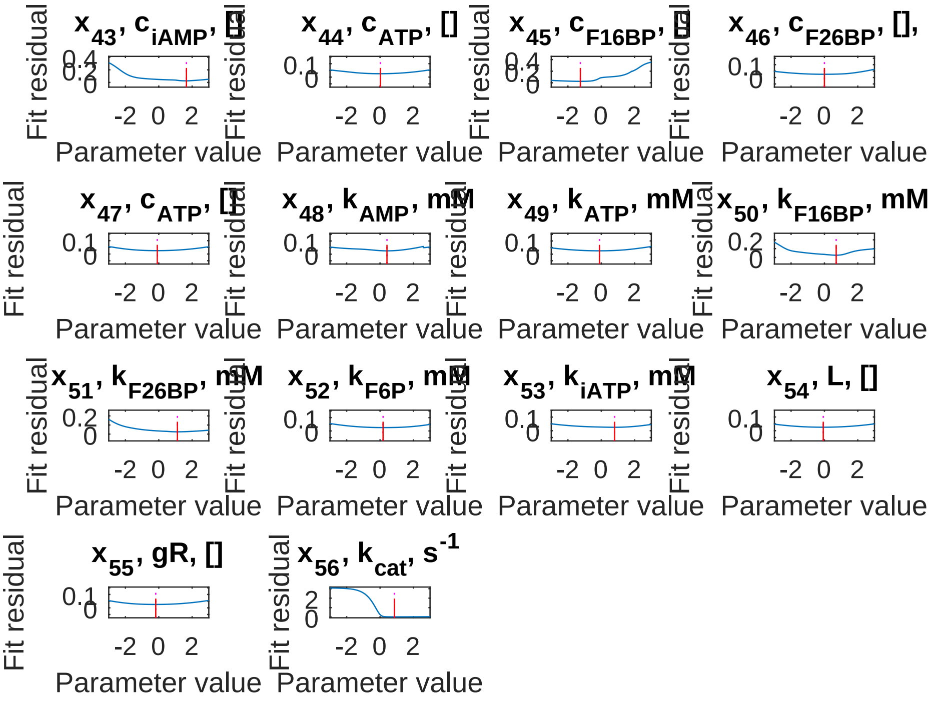


**FIGURE 6** **Phosphofructokinase. PLA, regularized.**
