## Supplementary material for "Glycolysis revisited: from steady state growth to glucose pulses": SM_DandC_PGI.pdf

### Phosphoglucoisomerase

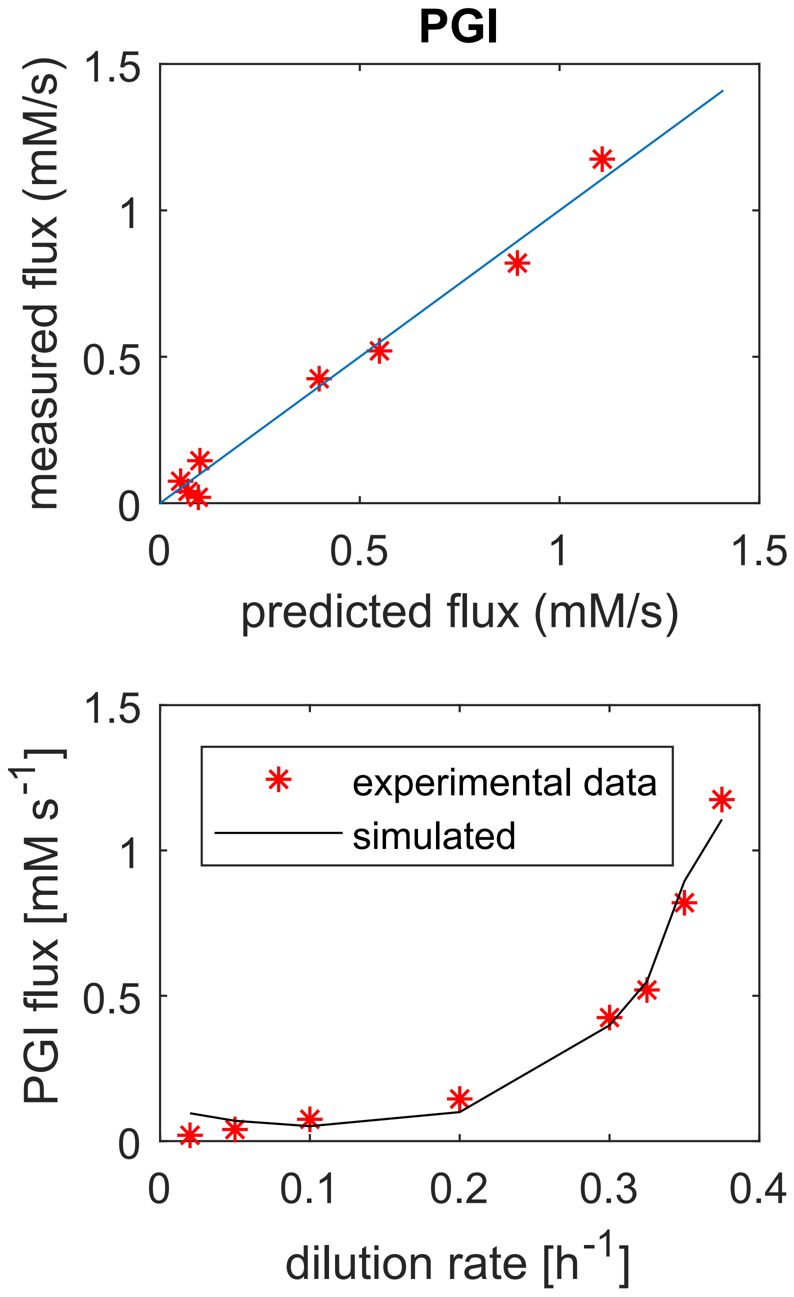


**FIGURE 1** Phosphoglucoisomerase. Model fit.

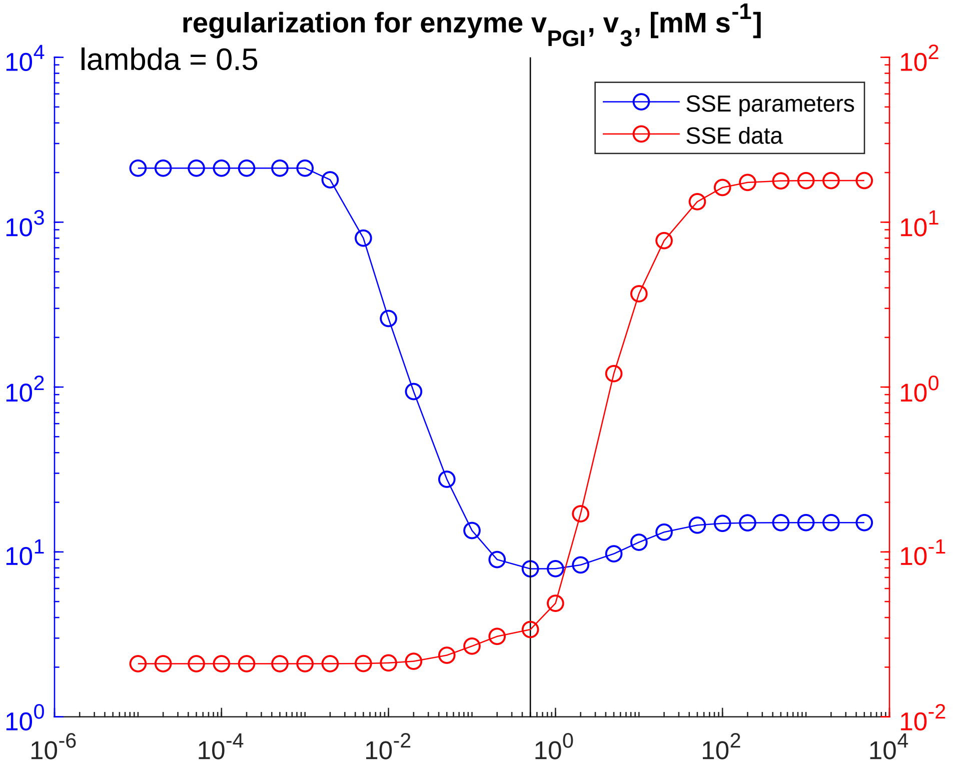


**FIGURE 2** **Phosphoglucoisomerase. Regularization.**

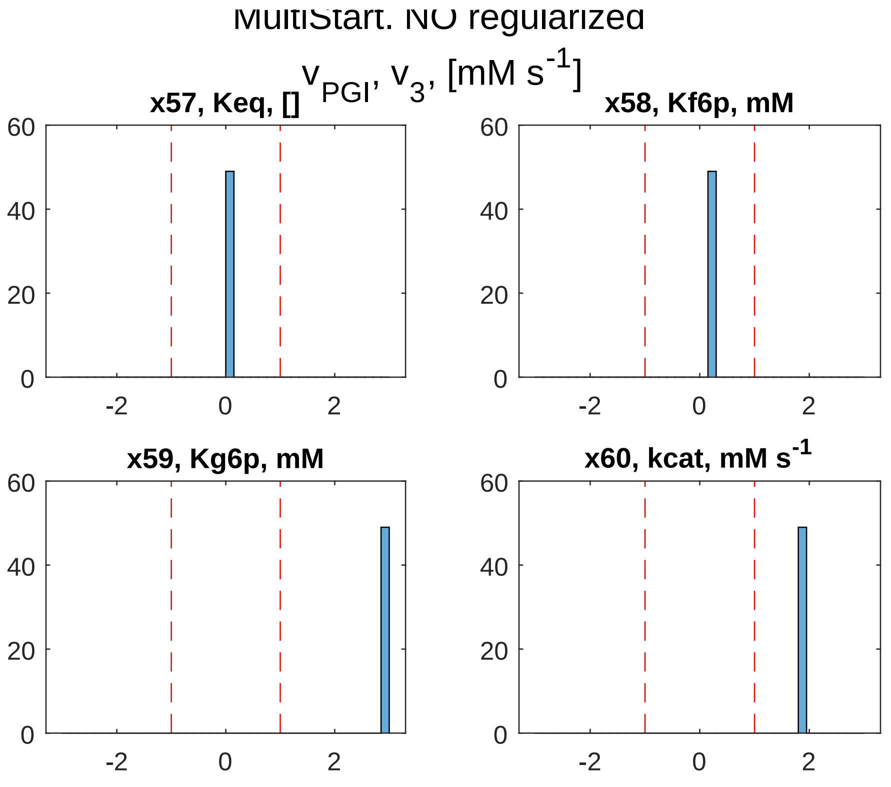


**FIGURE 3** Phosphoglucoisomerase. Global sampling, non-regularized.

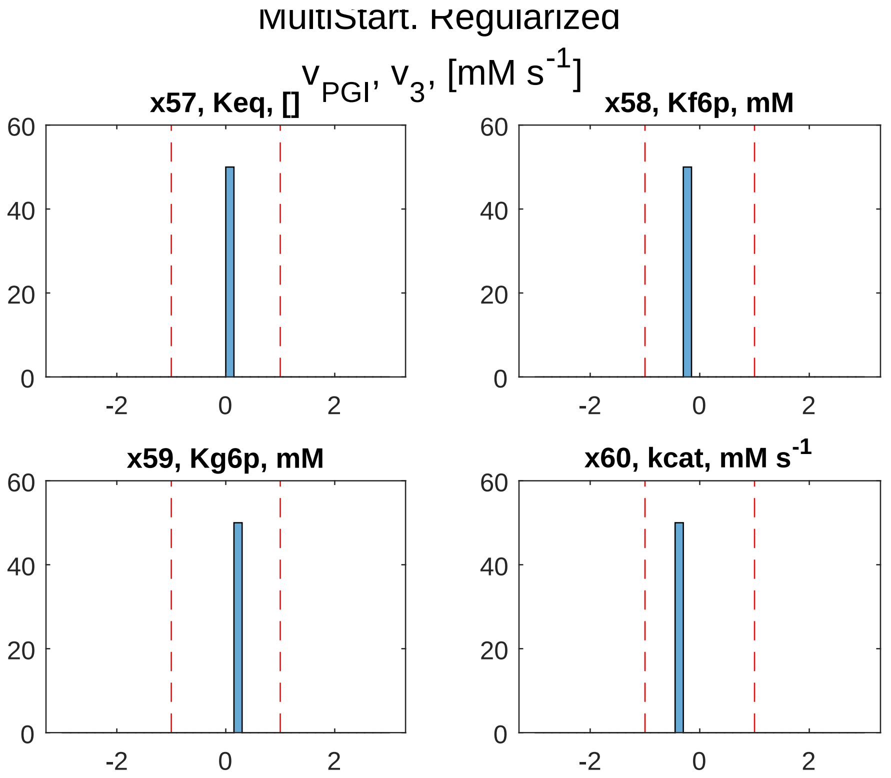


**FIGURE 4** Phosphoglucoisomerase. Global sampling, regularized.

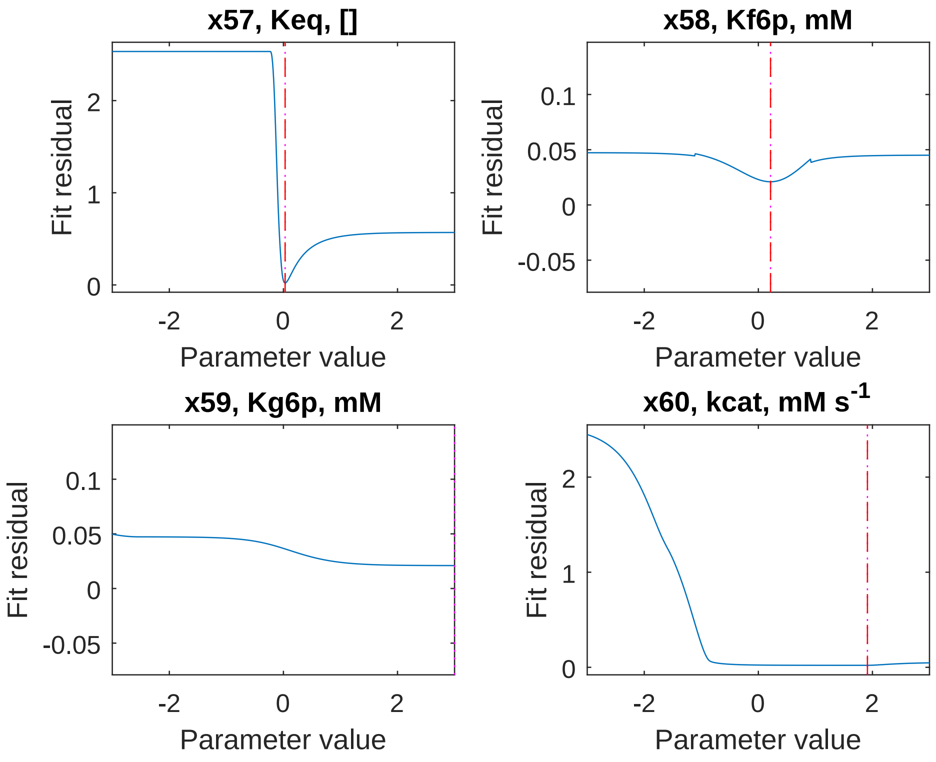


**FIGURE 5** Phosphoglucoisomerase. PLA, non-regularized.

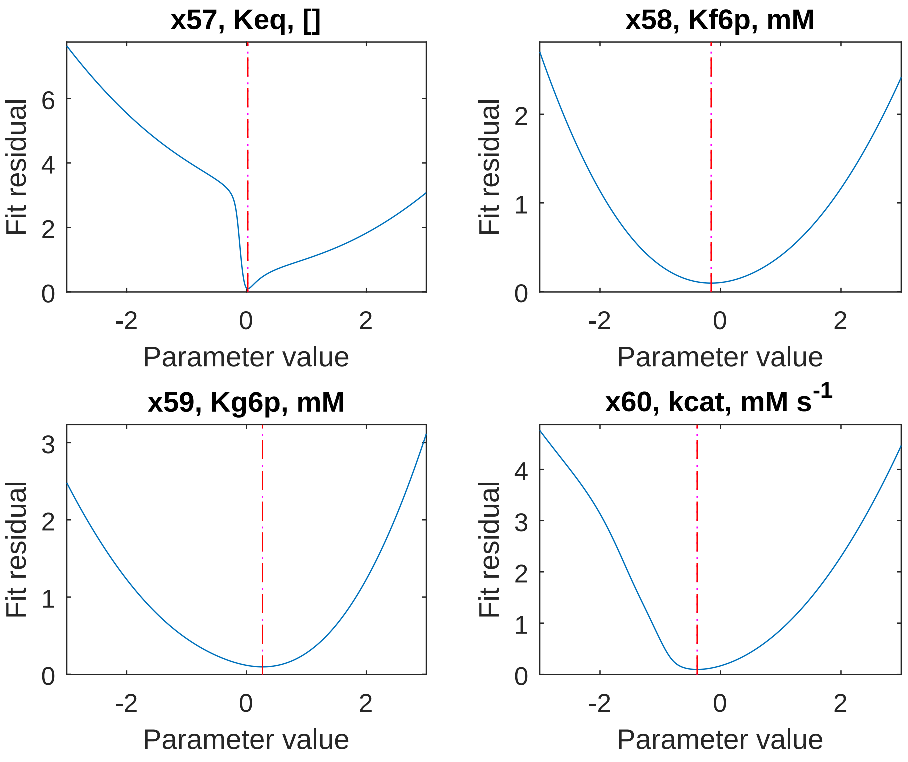


**FIGURE 6** Phosphoglucoisomerase. PLA, regularized.
