## Supplementary material for "Glycolysis revisited: from steady state growth to glucose pulses": SM_DandC_PGM.pdf

### Phosphoglucomutase

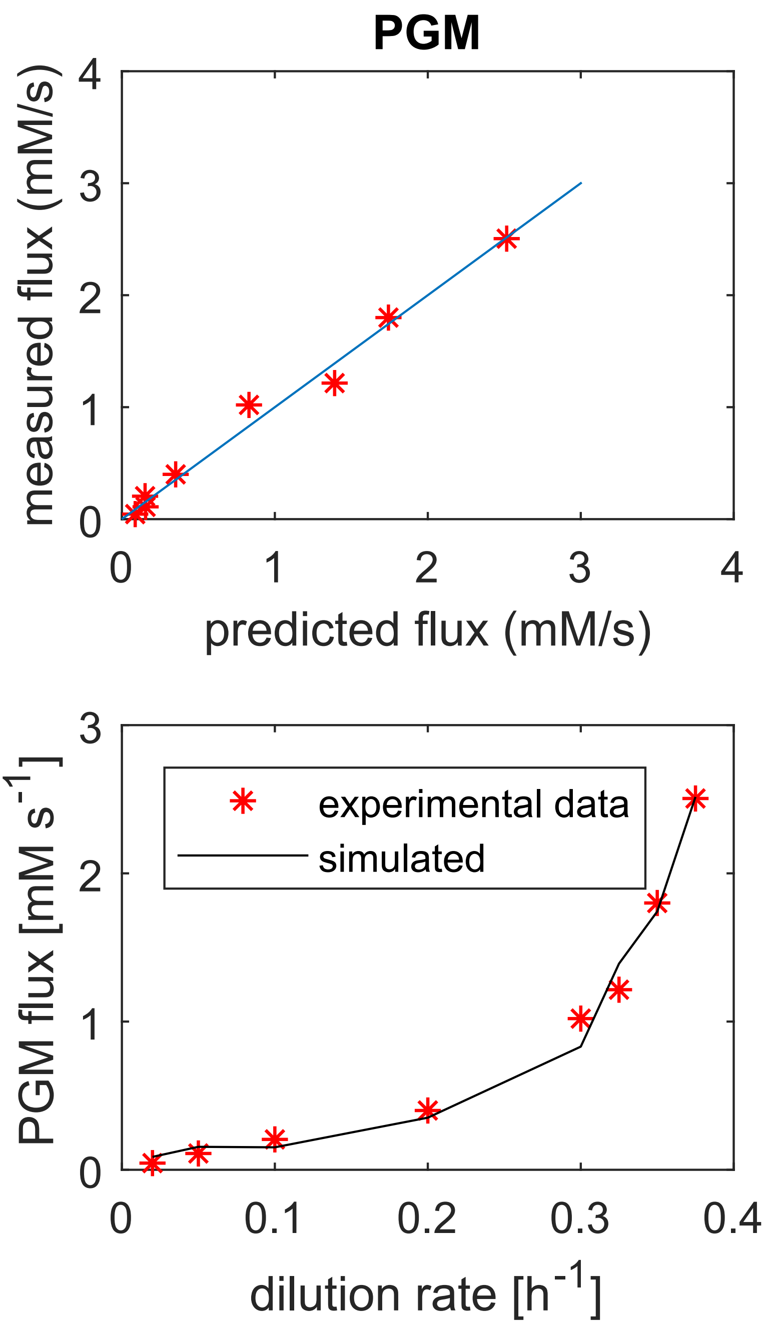


**FIGURE 1** Phosphoglucomutase. Model fit.

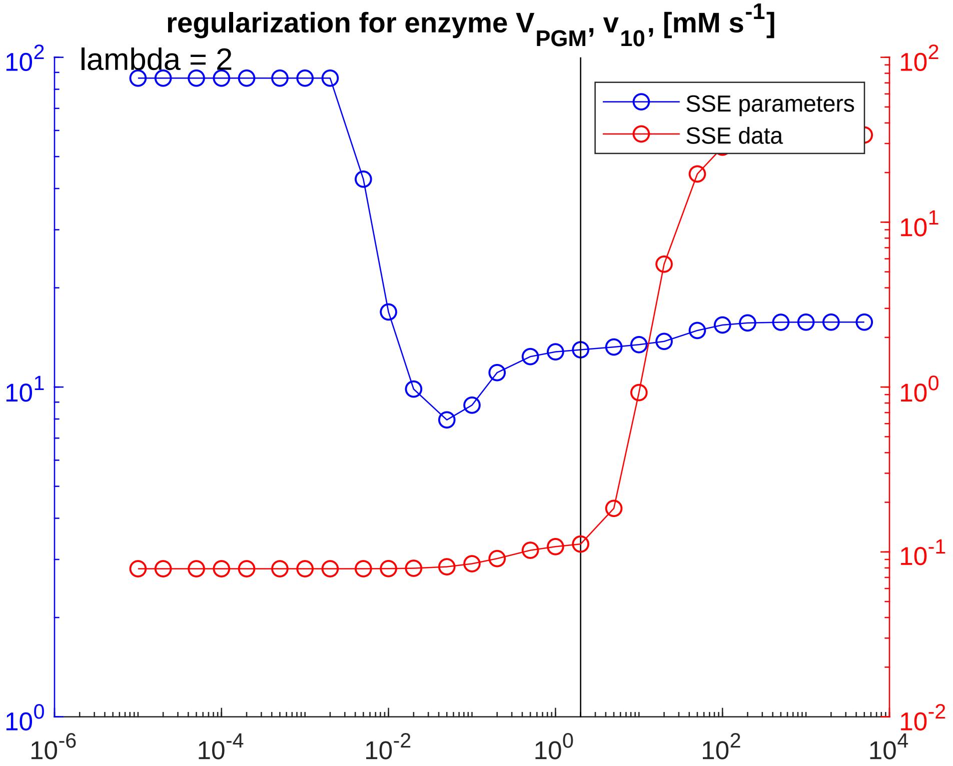


**FIGURE 2** Phosphoglucomutase. Regularization.

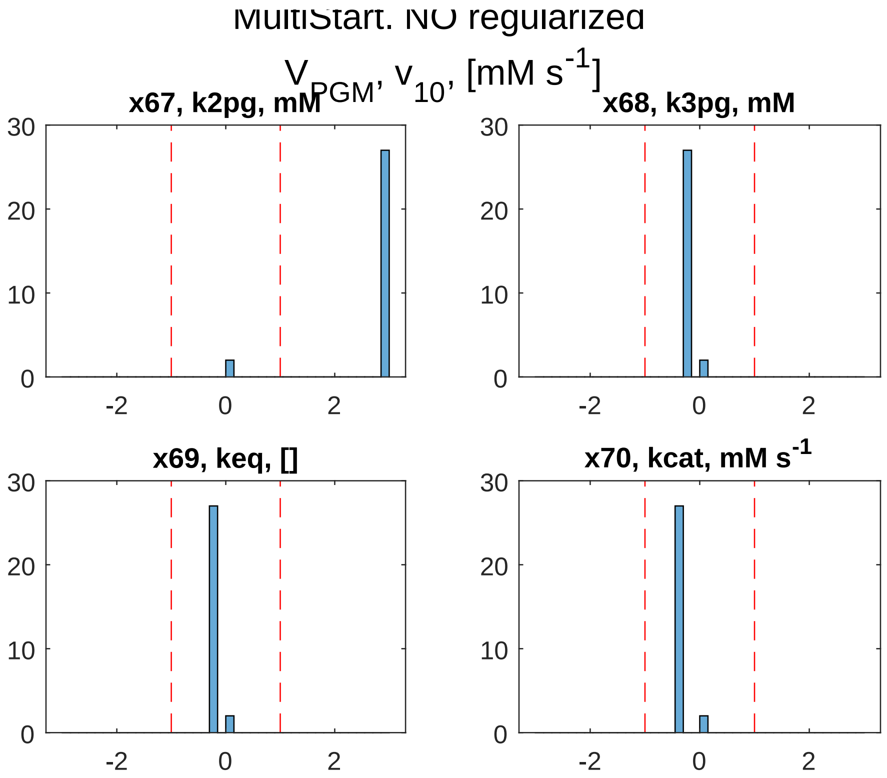


**FIGURE 3** Phosphoglucomutase. Global sampling, non-regularized.

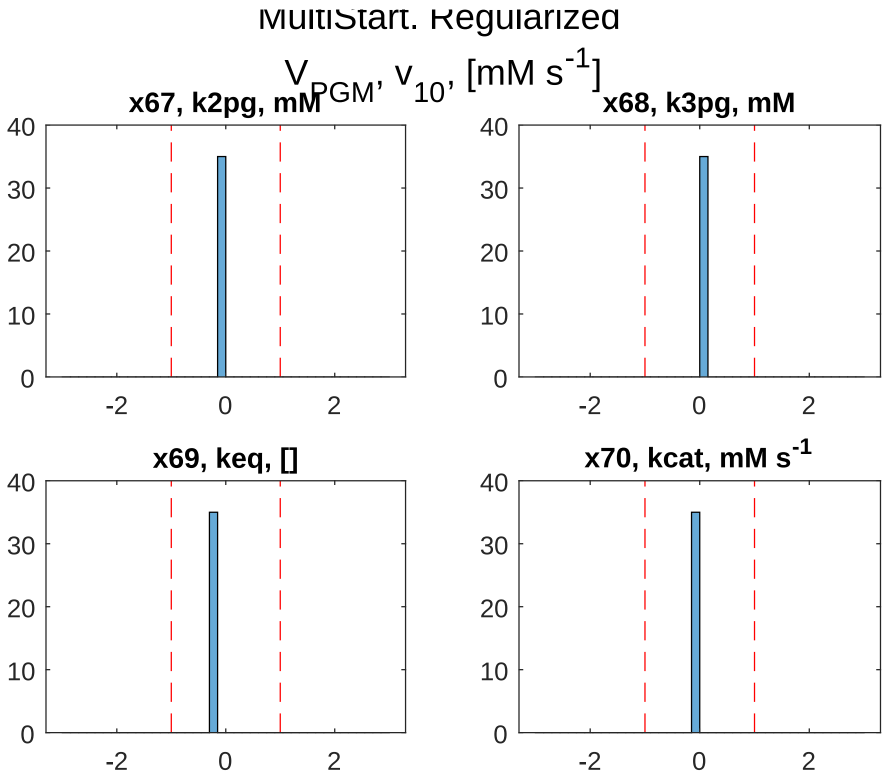


**FIGURE 4** **Phosphoglucomutase. Global sampling, regularized.**

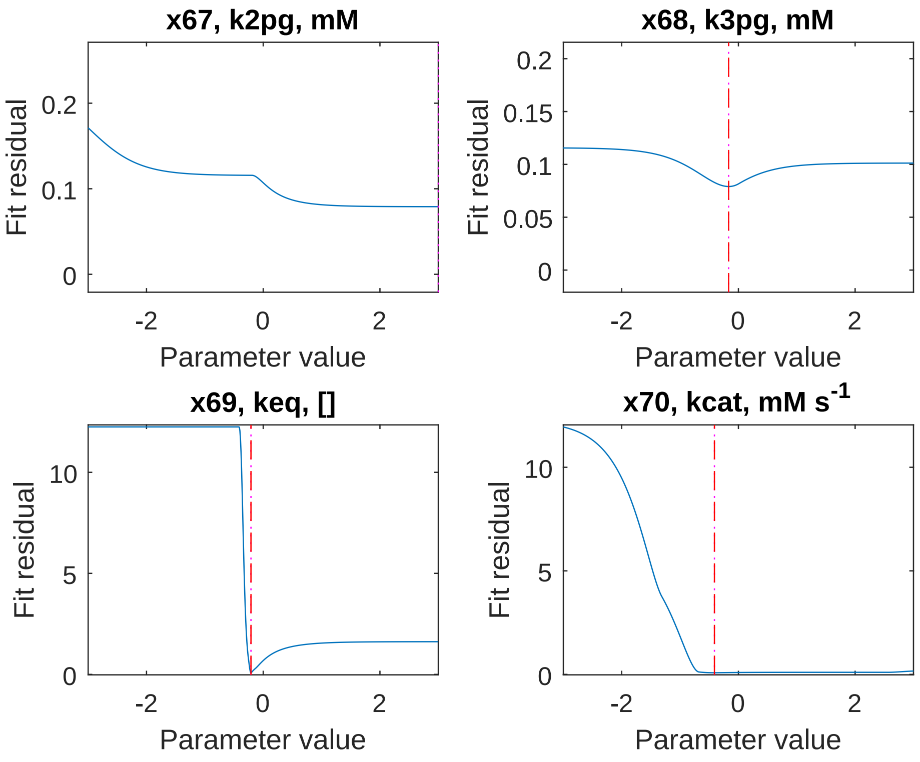


**FIGURE 5** Phosphoglucomutase. PLA, non-regularized.

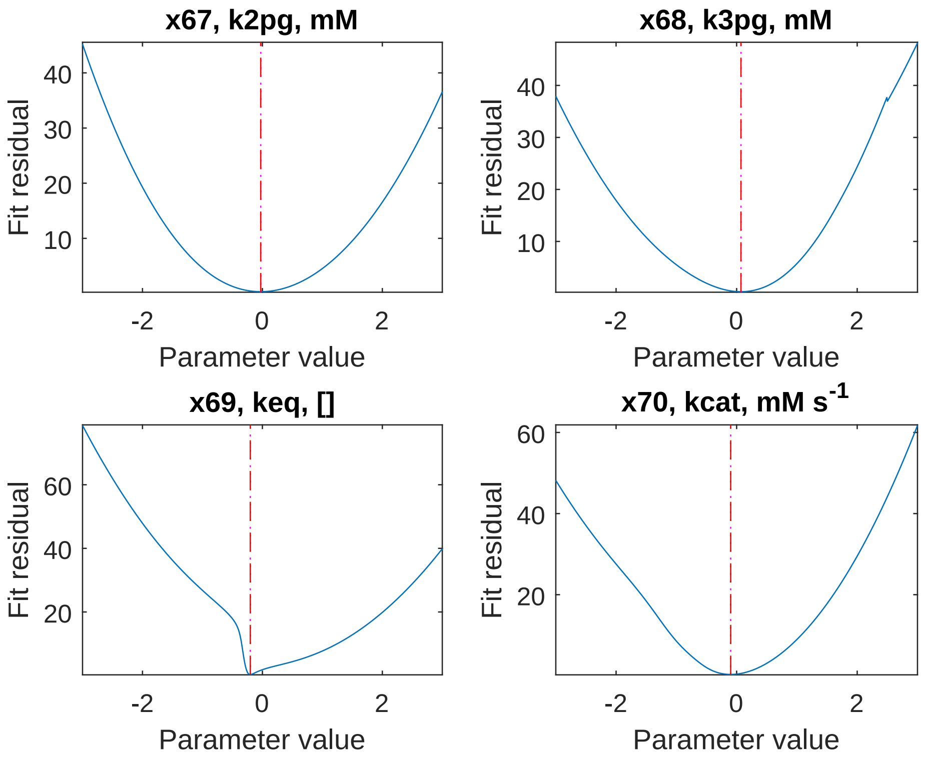


**FIGURE 6** **Phosphoglucomutase. PLA, regularized.**
