## Supplementary material for "Glycolysis revisited: from steady state growth to glucose pulses": SM_DandC_PYK.pdf

### Pyruvate kinase

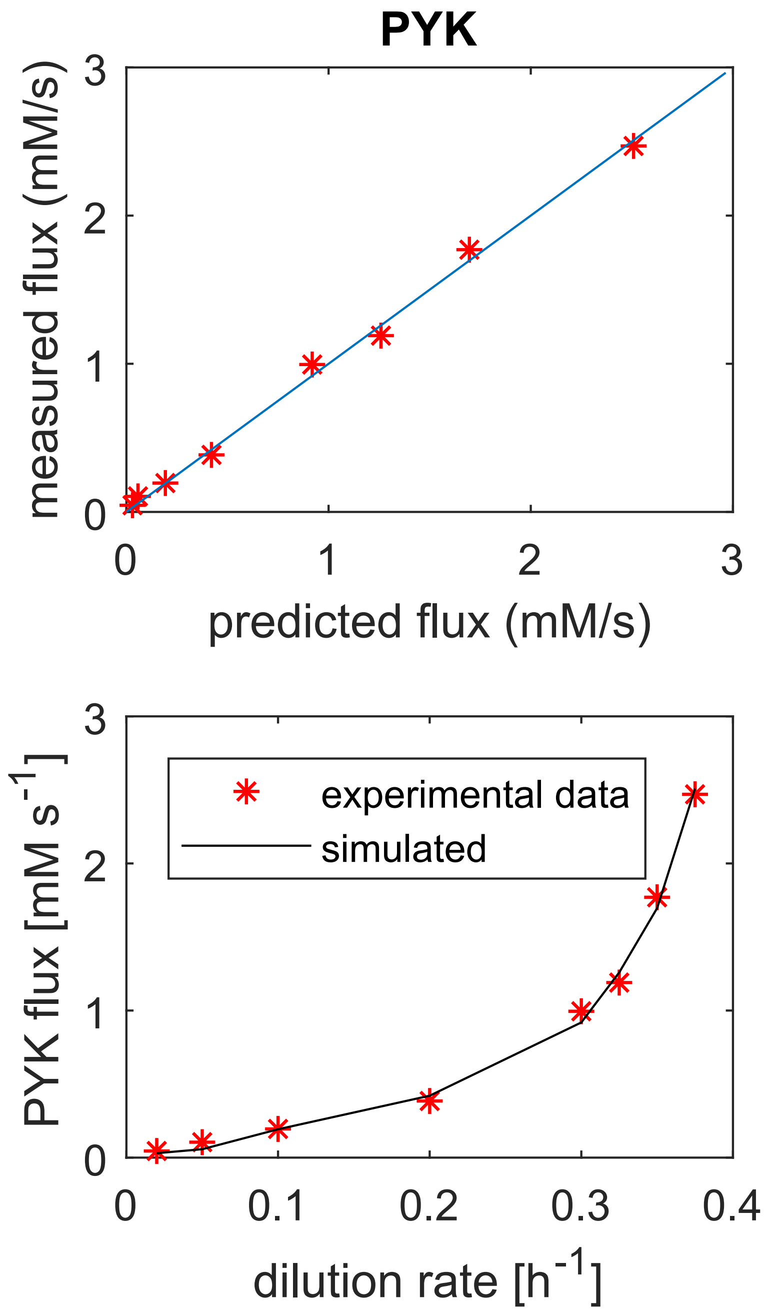


**FIGURE 1** Pyruvate kinase. Model fit.

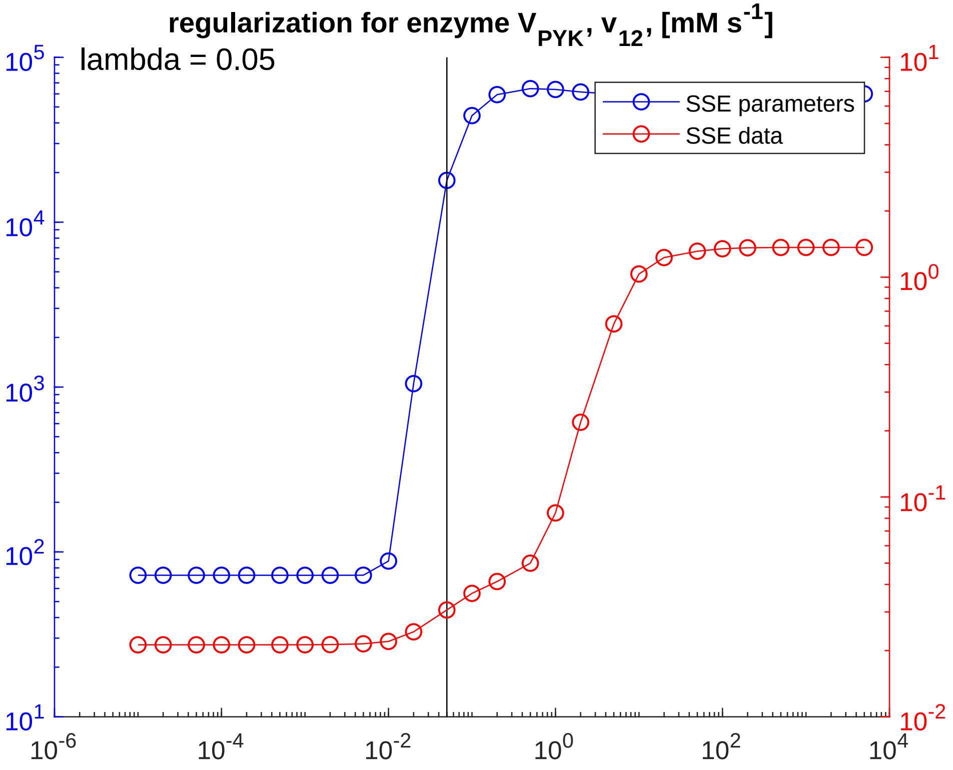


**FIGURE 2** **Pyruvate kinase. Regularization.**

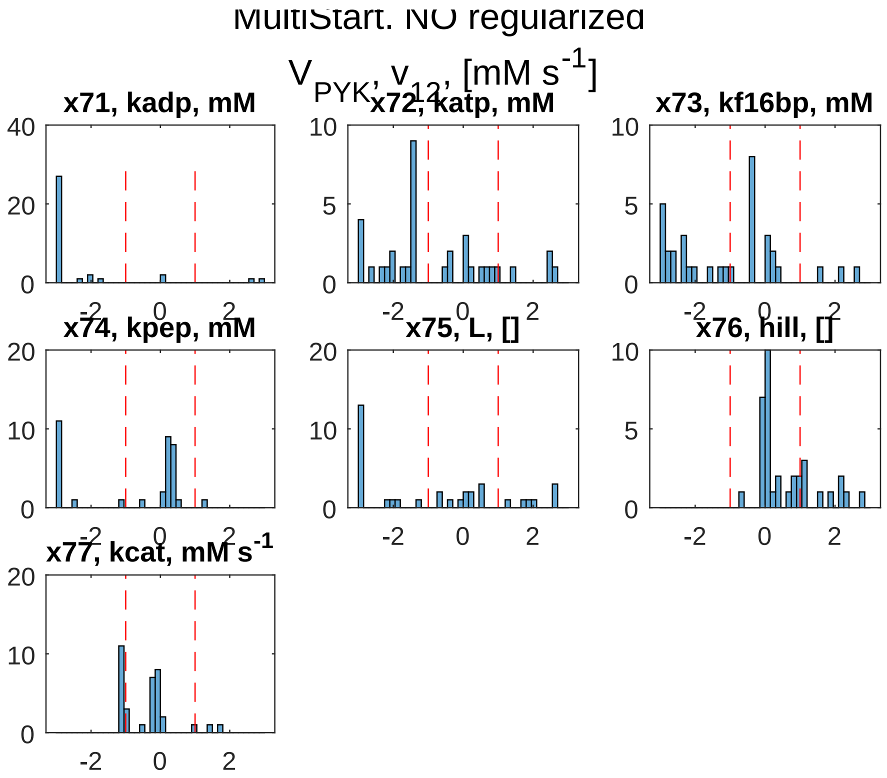


**FIGURE 3** Pyruvate kinase. Global sampling, non-regularized.

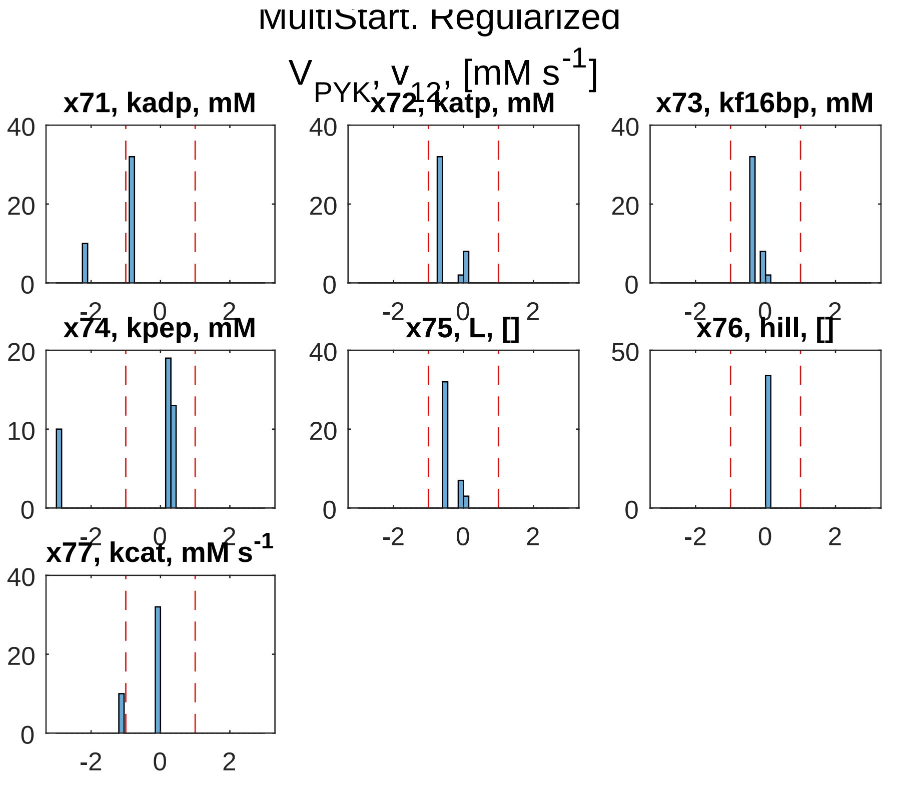


**FIGURE 4** Pyruvate kinase. Global sampling, regularized.

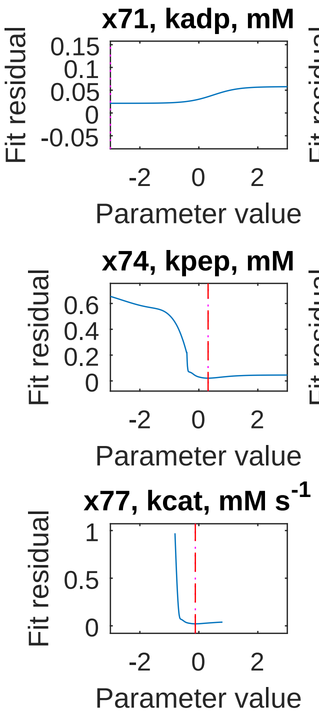


**FIGURE 5** **Pyruvate kinase. PLA, non-regularized.**

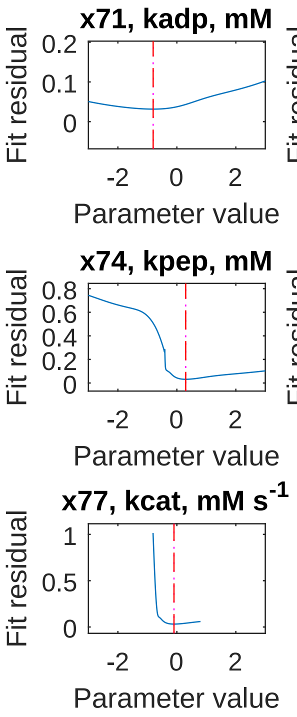


**FIGURE 6** Pyruvate kinase. PLA, regularized.
