## Supplementary material for "Glycolysis revisited: from steady state growth to glucose pulses": SM_DandC_TPI.pdf

### Triosephosphate isomerase

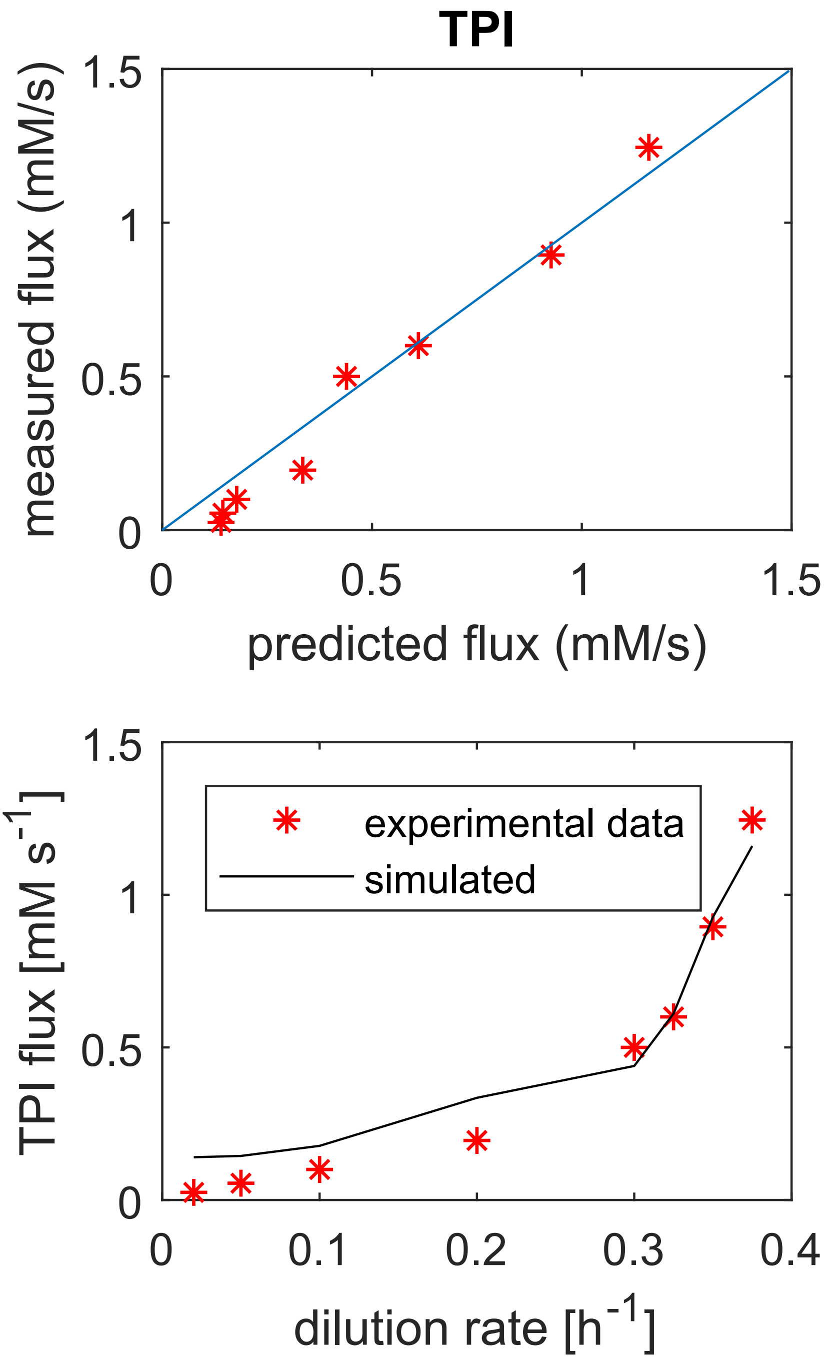


**FIGURE 1** Triosephosphate isomerase. Model fit.

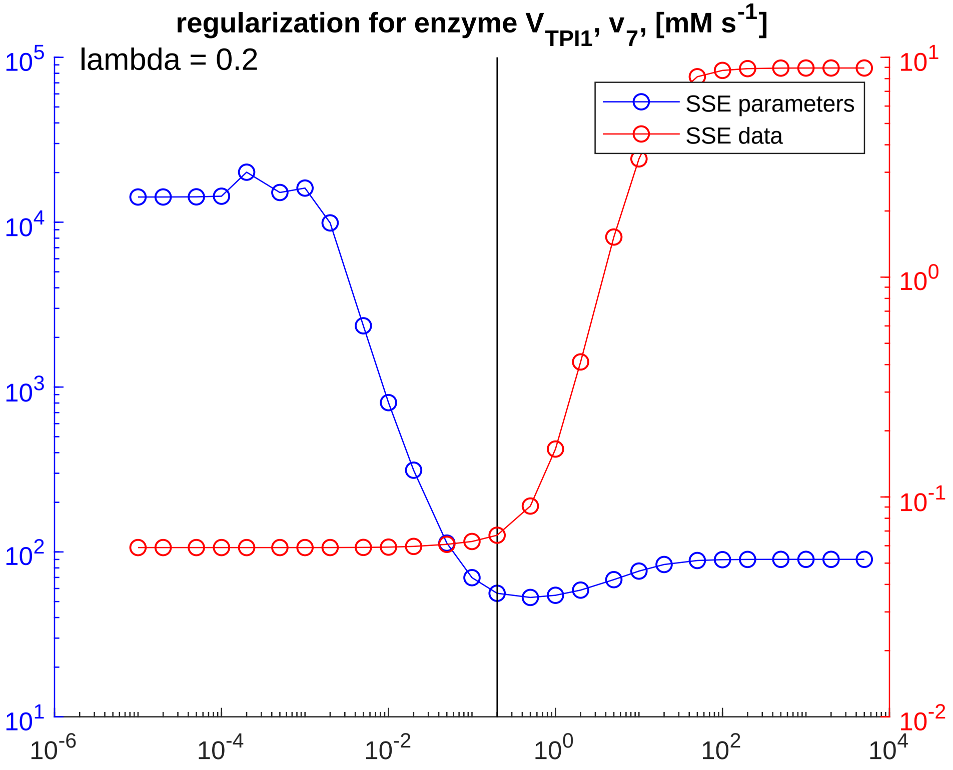


**FIGURE 2** **Triosephosphate isomerase. Regularization.**

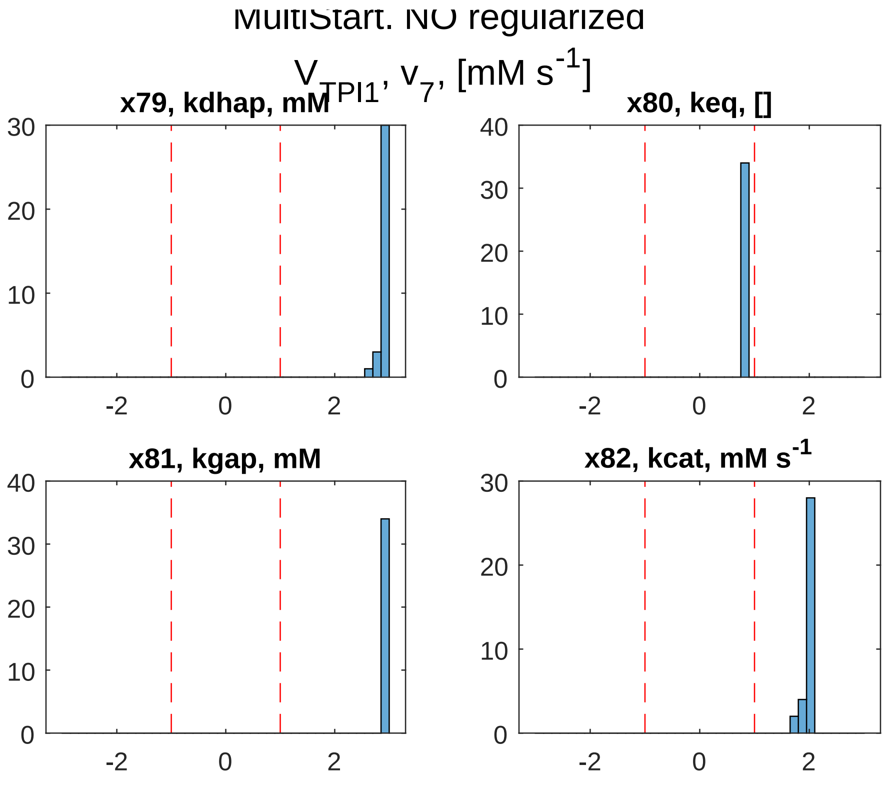


**FIGURE 3** **Triosephosphate isomerase. Global sampling, non-regularized.**

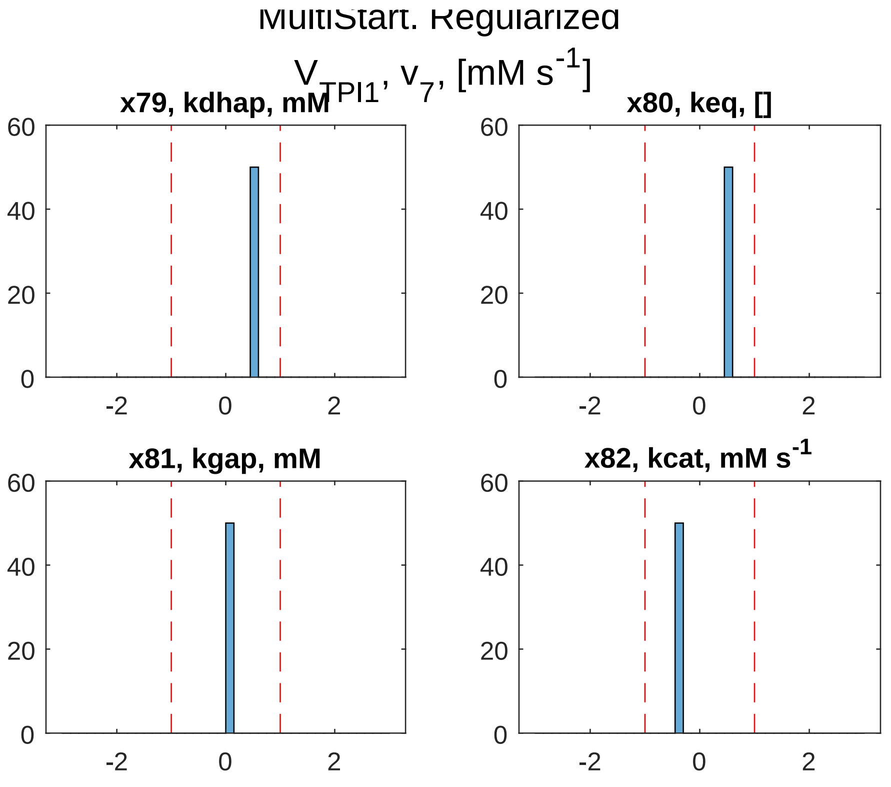


**FIGURE 4** Triosephosphate isomerase. Global sampling, regularized.

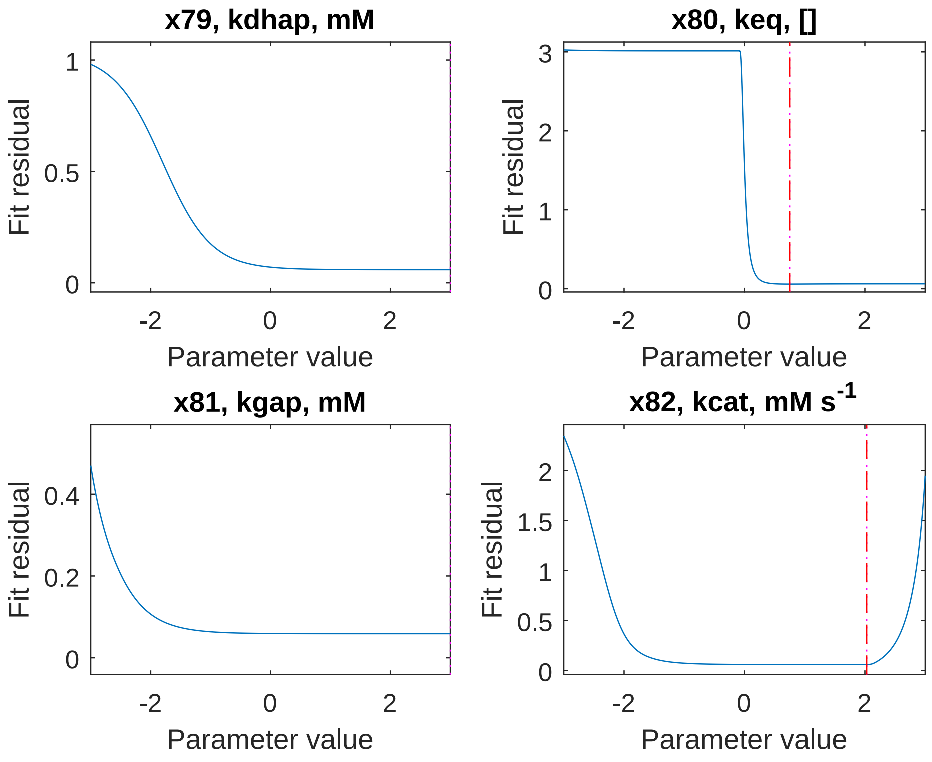


**FIGURE 5** **Triosephosphate isomerase. PLA, non-regularized.**

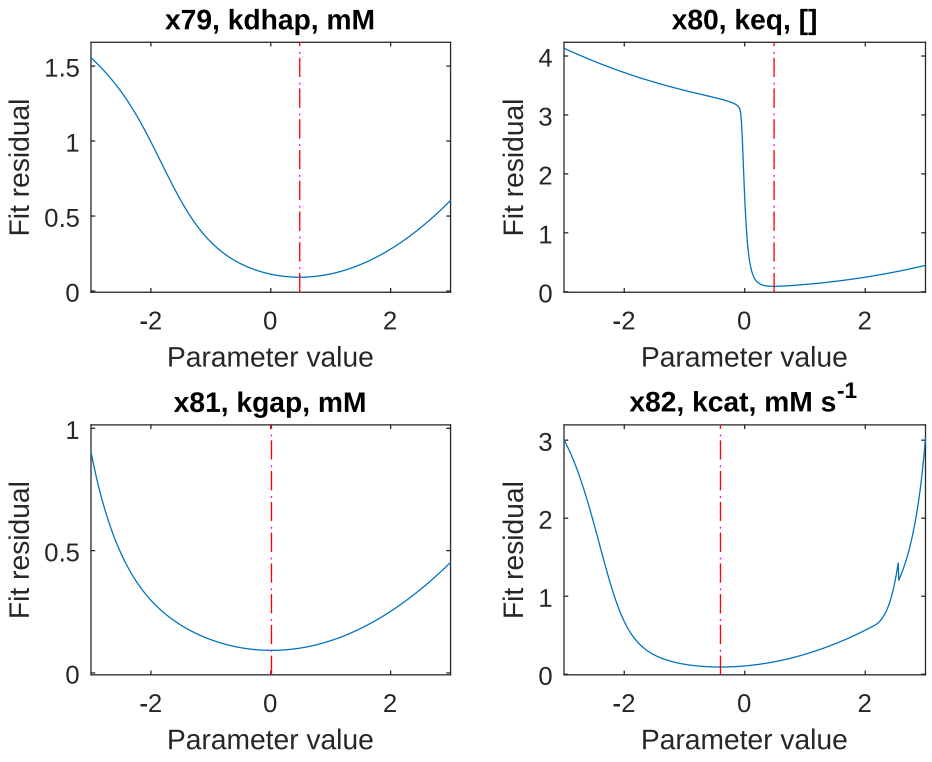


**FIGURE 6** **Triosephosphate isomerase. PLA, regularized.**
