## Supplementary material for "Glycolysis revisited: from steady state growth to glucose pulses": SM_complete_simulation.pdf

### Standard style simulation

The complete simulation of all model metabolites (states) and reaction rates (determining fluxes) can be seen in the following figures:

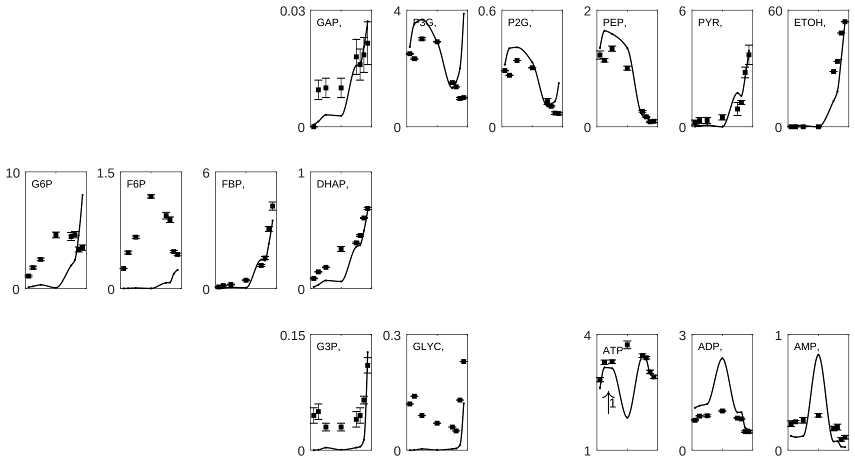


**FIGURE 1** **Steady state concentrations.** Concentrations are plotted in the Y-axis (in mM) and dilution rate in the X-axis (in $h^{-1}$).

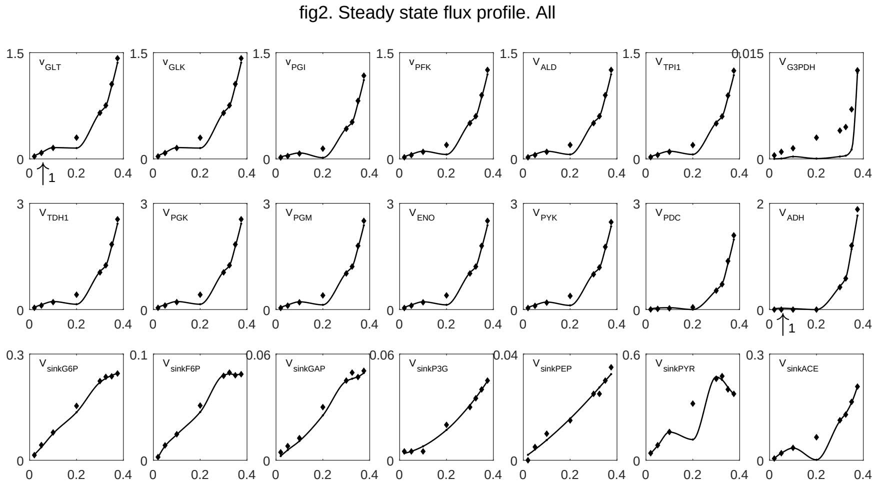


**FIGURE 2 Steady state reaction rates.** Rates are plotted in the Y-axis (in mM $s^{-1}$) and dilution rate in the X-axis (in $h^{-1}$).

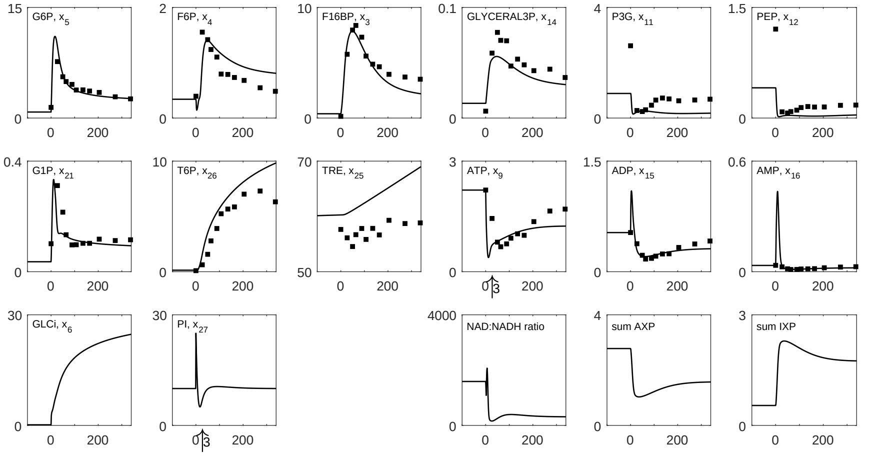


**FIGURE 3 Glucose perturbation concentrations.** Concentrations are plotted in the Y-axis (in mM) and time in the X-axis (in s).

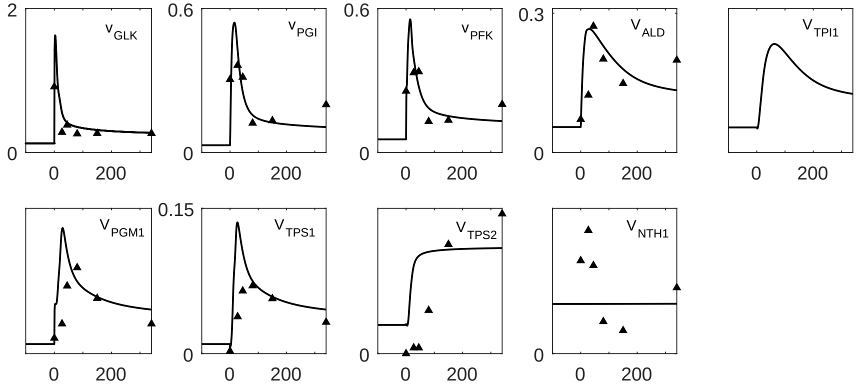


**FIGURE 4 Glucose perturbation reaction rates.** Rates are plotted in the Y-axis (in mM $s^{-1}$) and time in the X-axis (in s).
