## Supplementary material for "Glycolysis revisited: from steady state growth to glucose pulses": SM_hxt_d_dependency.pdf

### HXT adjustment. Vmax or Km

The activity of the glucose transporter could be adjusted in a dilution rate dependent manner. In line with (Diderich *et al*, 1999), where changes were observed in isoenzyme concentrations, these changes were made in the $V_{max}$ of the glucose transporter reaction. Nonetheless, changes in isoenzymes do also alter the affinity constants (Bosdriesz *et al*, 2018; Maier *et al*, 2002; Reifenberger *et al*, 1997). Fig 1 shows how indeed, the observed changes in the hexose tranporter activity could also be due to changes in $V_{max}$ as well as $K_m$.

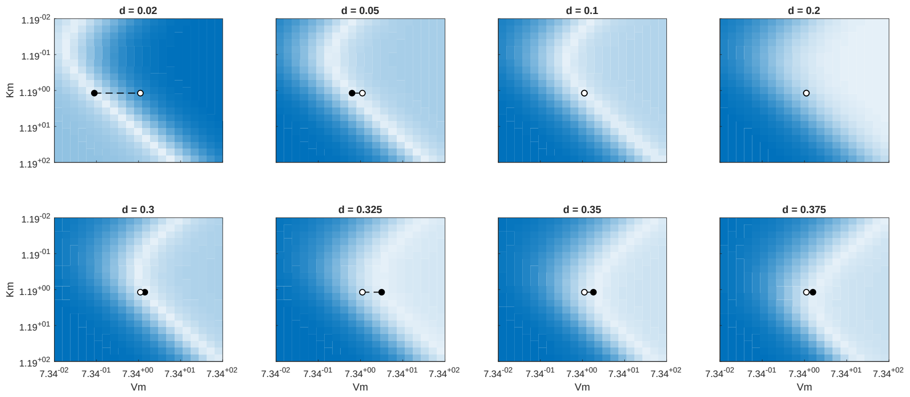


**FIGURE 1 HXT activity changes can be explained by both changes in the $V_{max}$ as well as $K_m$.** Lighter colors show the regions with low error and high agreement between simulations and experimental data. Darker regions show higher error. Each plot shows a dilution rate data point. The Y-axis shows the $K_m$ (in mM), and the X-axis the $V_{max}$ (in mM $s^{-1}$).
