## Supplementary material for "Glycolysis revisited: from steady state growth to glucose pulses": SM_maintenance_mitochondria.pdf

### Dilution rate dependent cofactor kinetics: maintenance and mitochondrial reactions

Cofactor metabolism is affected by dilution rate. The higher the dilution rate, the higher the growth rate. Depending on the growth rate, ATP maintenance and mitochondrial activity changes.

During the steady state simulations, the growth rate-dependent ATP maintenance was considered by setting the reaction mass action kinetic constant. This value was derived from the Growth Associated Maintenance (GAM) and Non Growth Association Maintenance (NGAM). This already made the model run smoothly. To fit the experimental steady state concentrations properly, the ratio between the experimental ADP and ATP concentrations was also included. During the single glucose perturbation simulation, the reaction constant value had to be increased compared to the steady state value at growth rate of $0.1^{-1}$. This could derive from the higher maintenance needs experienced during the strong glucose perturbation, not covered by the reactions in the model.

$$V_{\mathrm{ATPase}} = \frac{\mathrm{ATP}\,K}{\mathrm{ADP}} \tag{1}$$

$$K_{SS} = (NGAM + GAM * d)\frac{\mathrm{ADP_{exp}}}{\mathrm{ATP_{exp}}} \tag{2}$$

Mitochondrial activity is also growth rate dependent. This determines how much NADH is recycled and ATP synthesized by respiration. A direct experimental quantification of this activity was not available, but the sink reaction of pyruvate, whose product enters the mitochondria and is respired, can be used as a proxy of this activity. This flux increased with growth rate, and then decreased as fermentation became predominant (see Fig 1):

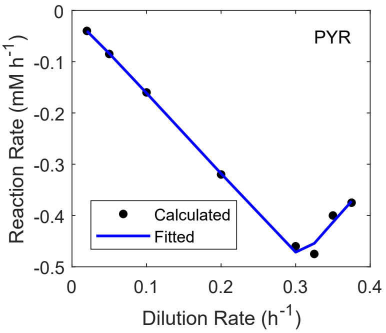


**FIGURE 1 Pyruvate Sink reaction rate.** Reaction rate is plotted against dilution rate. Fits are shown as a blue line and experimental data points as black dots.

Therefore, to calculate the steady state mitochondrial ATP synthesis, first the $q_{CO2}$ was calculated adding together the carbon mols from the sink of pyruvate and the PDC reaction rate. The $q_{O2}$ was then calculated considering the experimental RQ ratio. The synthesis of mitochondrial ATP was then derivated by multiplying by the PO ratio (Verduyn *et al*, 1991).

$$q_{CO2} = -v_{sinkPYR} * 3 + v_{PDC} * 1 \tag{3}$$

$$q_{O2} = q_{CO2} * RQ_{ratio} \tag{4}$$

$$V_{mito.SS} = q_{O2} * PO_{ratio} \tag{5}$$

During the glucose perturbation, standard format Michaelis Menten kinetics were assumed:

$$V_{\text{mito.GP}} = \frac{\text{ADP PI } V_m}{\left(\text{ADP} + K_{m,\text{ADP}}\right)\left(K_{m,\text{PI}} + \text{PI}\right)} \tag{6}$$

To account for the growth rate dependency in the NADH mitochondrial recycle, the maximum reaction rate of the reaction was adjusted accorded to how much the pyruvate sink flux would change respect to 0.1 $h^{-1}$ (this dilution rate value was taken just to have a reference). A constant C was added to modulate this effect, and set to a value where simulations would not be negatively affected (C = 2).

$$V_{m,d} = V_{m,0.1} * 10^{C*(V_{sinkPYR,d} - V_{sinkPYR,0.1})} \tag{7}$$

$$V_{\text{mitoNADH}} = \frac{\text{NADH } V_{m,d}}{K_m + \text{NADH}} \tag{8}$$
