## Supplementary material for "Glycolysis revisited: from steady state growth to glucose pulses": SM_parameters.pdf

### Parameter estimates comparison to the literature

The parameters estimated in this work can be compared with the literature. In the following plots, the ratio between estimated and literature parameter is displayed, parameter by parameter. Fig 1) shows the comparison for all parameters, Fig 2), 3) and 4) show the comparison, bu only for $V_{max}$, $K_m$ and $K_{eq}$.

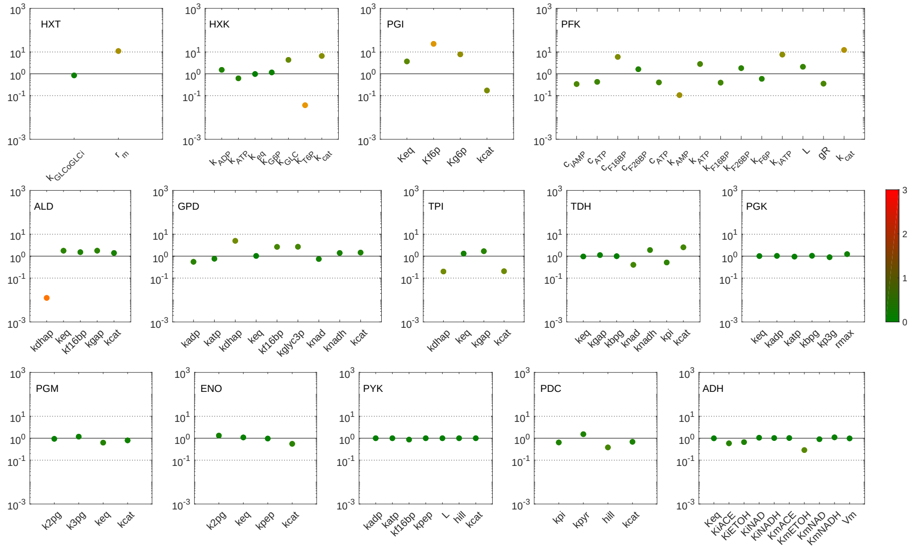


**FIGURE 1** **Estimated vs Literature parameters: all parameters.**

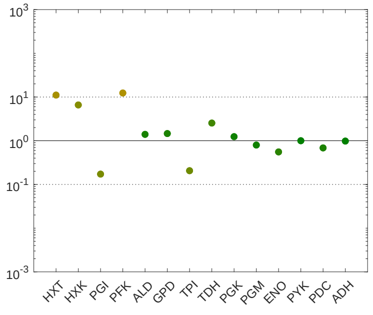


**FIGURE 2** **Estimated vs Literature parameters: $V_{max}$.**

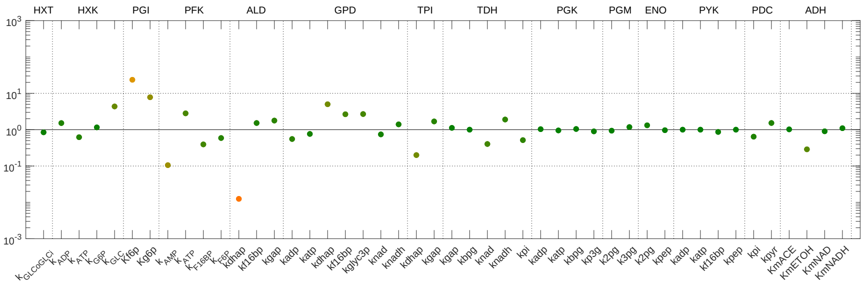


**FIGURE 3** **Estimated vs Literature parameters: $K_m$.**

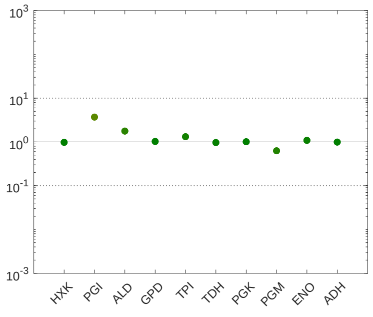


**FIGURE 4** **Estimated vs Literature parameters: $K_{eq}$.**
