## Supplementary material for "Glycolysis revisited: from steady state growth to glucose pulses": SM_pfk_d_dependency.pdf

### Fudge factor PFK adjustment

To test that the difference between simulated and experimental steady state (SS) concentrations of fructose 6-phosphate (F6P) could be explained by changes in phosphofructokinase (PFK) a dilution rate-dependent fudge factor was added to the reaction kinetics of this enzyme. The followign equation shows how the fudge factor was implemented. Fig 1 shows how the model fit could be improved. The fudge factor values are shown in Table 1.

$$v_{\text{PFK,d-dependent}} = \text{fudge}_{d-dependent}\, v_{\text{PFK}} \tag{1}$$

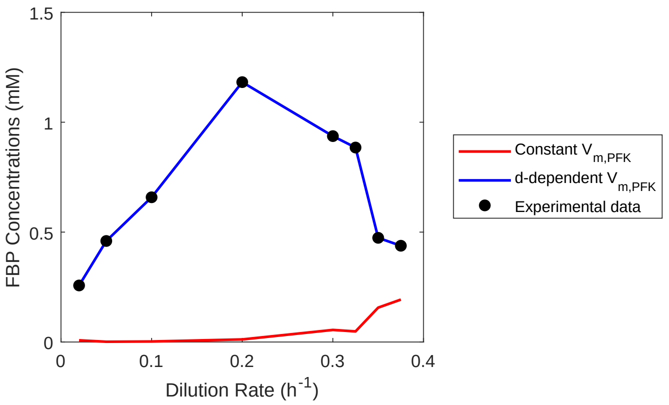


**FIGURE 1 A dilution rate-dependent fudge factor on PFK kinetics improved model fit for SS concentrations of F6P.** The red and blue lines show the model simulations when dilution rate dependency is considered or not, respectively. The black dots show the experimental data. F6P concentrations are plotted in the Y-axis (mM) and dilution rate in the X-axis ($h^{-1}$).

**TABLE 1 Dilution rate-dependent fudge factor values.**

| Dilution rate ($h^{-1}$) | Fudge factor (unitless, log scale) | Fudge factor (unitless) |
|---|---|---|
| 0.0200 | -1.78 | 0.0167 |
| 0.0500 | -2.11 | 0.0078 |
| 0.1000 | -1.98 | 0.0106 |
| 0.2000 | -1.67 | 0.0212 |
| 0.3000 | -1.25 | 0.0564 |
| 0.3250 | -1.25 | 0.0557 |
| 0.3500 | -0.71 | 0.1950 |
| 0.3750 | -0.55 | 0.2821 |
