## Supplementary material for "Glycolysis revisited: from steady state growth to glucose pulses": SM_proteomics.pdf

### Implementation of dilution rate-dependent glycolytic enzyme activity

The growth rate dependent change in protein activity was considered in this work. We used the experimental glycolytic enzymes activity quantification between dilution rates of 0.025-0.4 $h^{-1}$ (van Hoek *et al*, 2000). The experimental setup was also chemostats as in the experimental steady sate data used in this work (Canelas *et al*, 2011), making this comparison more reliable. The experimental data from (van Hoek *et al*, 2000) is plotted in Fig 1:

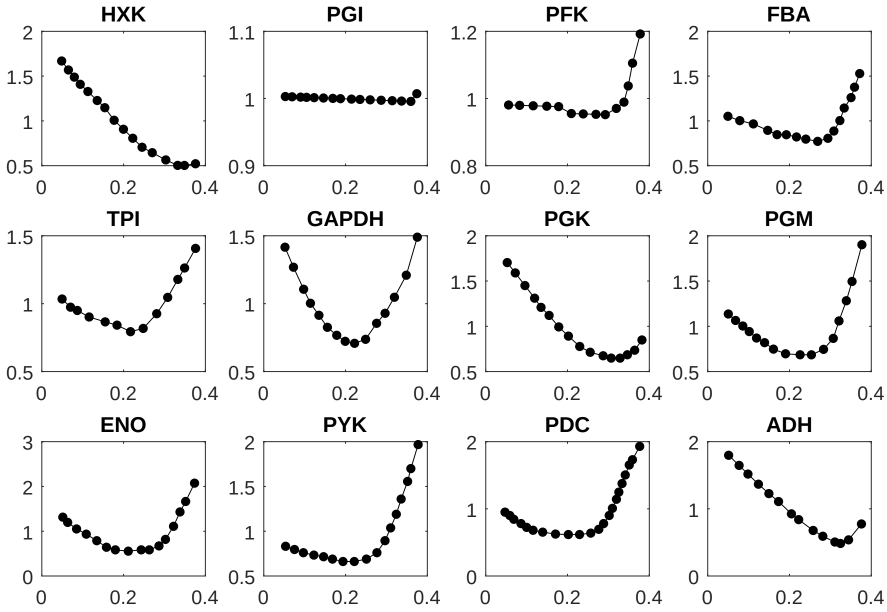


**FIGURE 1 Experimental protein activity in the (van Hoek *et al*, 2000) dataset.** Activity is plotted in the Y-axis (unitless) and dilution rate in the X-axis (in $h^{-1}$).

For all the glycolytic enzymes for which the experimental data was available, enzymatic activity was interpolated at a given dilution rate. A ratio was then calculated between this interpolated value and the experimental value at 0.1 $h^{-1}$ accounting for a relative change in activity. The value at 0.1 $h^{-1}$ was taken as a reference, since the kinetic parameters in the model and the single glucose perturbation simulations were performed in this dilution rate. This ratio was then use to alter the $V_{max}$ for each glycolytic enzyme. The example below shows the case for HXK reaction rate:

$$\mathrm{HXK}_{\mathrm{cor,d}} = \frac{\mathrm{activity}_{\mathrm{HXK,d}}}{\mathrm{activity}_{\mathrm{HXK,0.1}}} \tag{1}$$

$$v_{\mathrm{HXK}} = \frac{\mathrm{HXK}_{\mathrm{cor,d}}\, V_m \left(\mathrm{ATP\,GLCi} - \frac{\mathrm{ADP\,G6P}}{K_{\mathrm{eq}}}\right)}{K_{m,\mathrm{glc}}\, K_{m,\mathrm{atp}} \left(\frac{\mathrm{ADP}}{K_{m,\mathrm{adp}}} + \frac{\mathrm{ATP}}{K_{m,\mathrm{atp}}} + 1\right) \left(\frac{\mathrm{G6P}}{K_{m,\mathrm{g6p}}} + \frac{\mathrm{GLCi}}{K_{m,\mathrm{glc}}} + \frac{\mathrm{T6P}}{K_{i,\mathrm{t6p}}} + 1\right)} \tag{2}$$
