## Supplementary material for "Glycolysis revisited: from steady state growth to glucose pulses": SM_rate_equations.pdf

The following kinetic expressions were used:

$$v_{\mathrm{GLT}} = \frac{V_m \left(\mathrm{GLCo} - \frac{\mathrm{GLCi}}{K_{\mathrm{eq}}}\right)}{K_m \left(\frac{\mathrm{GLCi}}{K_m} + \frac{\mathrm{GLCo}}{K_m} + \frac{\mathrm{GLCi}\,\mathrm{GLCo}\,K_i}{{K_m}^2} + 1\right)} \tag{1}$$

$$v_{\mathrm{HXK}} = \frac{\mathrm{HXK_{cor}}\,V_m \left(\mathrm{ATP}\,\mathrm{GLCi} - \frac{\mathrm{ADP}\,\mathrm{G6P}}{K_{\mathrm{eq}}}\right)}{K_{m,\mathrm{glc}}\,K_{m,\mathrm{atp}} \left(\frac{\mathrm{ADP}}{K_{m,\mathrm{adp}}} + \frac{\mathrm{ATP}}{K_{m,\mathrm{atp}}} + 1\right) \left(\frac{\mathrm{G6P}}{K_{m,\mathrm{g6p}}} + \frac{\mathrm{GLCi}}{K_{m,\mathrm{glc}}} + \frac{\mathrm{T6P}}{K_{i,\mathrm{t6p}}} + 1\right)} \tag{2}$$

$$v_{\mathrm{PGM1}} = \frac{V_m \left(\mathrm{G1P} - \frac{\mathrm{G6P}}{K_{\mathrm{eq}}}\right)}{K_{m,\mathrm{G1P}} \left(\frac{\mathrm{G1P}}{K_{m,\mathrm{G1P}}} + \frac{\mathrm{G6P}}{K_{m,\mathrm{G6P}}} + 1\right)} \tag{3}$$

$$v_{\mathrm{TPS1}} = \frac{\mathrm{F6P}\,\mathrm{G6P}\,\mathrm{UDPGLC}\,V_m}{K_{m,\mathrm{G6P}}\,K_{m,\mathrm{UDPGLC}} \left(\frac{\mathrm{G6P}}{K_{m,\mathrm{G6P}}} + 1\right) \left(\frac{\mathrm{PI}}{K_{i,\mathrm{PI}}} + 1\right) \left(\frac{\mathrm{UDPGLC}}{K_{m,\mathrm{UDPGLC}}} + 1\right) \left(\mathrm{F6P} + K_{m,\mathrm{F6P}}\right)} \tag{4}$$

$$v_{\mathrm{TPS2}} = \frac{\mathrm{PI}\,\mathrm{T6P}\,V_m}{\mathrm{PI}\,\left(K_{m,\mathrm{T6P}} + \mathrm{T6P}\right) + K_{i,\mathrm{PI}}\,K_{m,\mathrm{T6P}}} \tag{5}$$

$$v_{\mathrm{NTH1}} = \frac{\mathrm{TRE}\,V_m}{K_{m,\mathrm{TRE}} \left(\frac{\mathrm{TRE}}{K_{m,\mathrm{TRE}}} + 1\right)} \tag{6}$$

$$v_{\mathrm{PGI}} = \frac{\mathrm{PGI_{cor}}\,V_m \left(\mathrm{G6P} - \frac{\mathrm{F6P}}{K_{\mathrm{eq}}}\right)}{K_{m,\mathrm{g6p}} \left(\frac{\mathrm{F6P}}{K_{m,\mathrm{f6p}}} + \frac{\mathrm{G6P}}{K_{m,\mathrm{g6p}}} + 1\right)} \tag{7}$$

$$v_{\mathrm{PFK}} = \frac{\mathrm{PFK_{cor}}\,R\,g_R\,\mathrm{lam}_1\,\mathrm{lam}_2\,v_m}{R^2 + L\,T^2} \tag{8}$$

$$\mathrm{lam}_1 = \frac{\mathrm{F6P}}{K_{R,\mathrm{F6P}}} \tag{9}$$

$$\mathrm{lam}_2 = \frac{\mathrm{ATP}}{K_{R,\mathrm{ATP}}} \tag{10}$$

$$R = \mathrm{lam}_1\,\mathrm{lam}_2 + g_R\,\mathrm{lam}_1\,\mathrm{lam}_2 + 1 \tag{11}$$

$$T = c_{\mathrm{ATP}}\,\mathrm{lam}_2 + 1 \tag{12}$$

$$L = \frac{L_0\left(\frac{\mathrm{AMP}\,c_{i,\mathrm{AMP}}}{K_{\mathrm{AMP}}} + 1\right)^2\left(\frac{\mathrm{ATP}\,c_{i,\mathrm{ATP}}}{K_{\mathrm{ATP}}} + 1\right)^2\left(\frac{\mathrm{F26bP}\,c_{i,\mathrm{F26bp}}}{K_{\mathrm{F26bP}}} + \frac{\mathrm{FBP}\,c_{i,\mathrm{FBP}}}{K_{\mathrm{FBP}}} + 1\right)}{\left(\frac{\mathrm{AMP}}{K_{\mathrm{AMP}}} + 1\right)^2\left(\frac{\mathrm{ATP}}{K_{\mathrm{ATP}}} + 1\right)^2\left(\frac{\mathrm{F26bP}}{K_{\mathrm{F26bP}}} + \frac{\mathrm{FBP}}{K_{\mathrm{FBP}}} + 1\right)} \tag{13}$$

$$v_{\mathrm{ALD}} = \frac{\mathrm{FBA_{cor}}\,V_m\left(\mathrm{FBP} - \frac{\mathrm{DHAP\,GAP}}{K_{\mathrm{eq}}}\right)}{K_{m,\mathrm{FBP}}\left(\frac{\mathrm{FBP}}{K_{m,\mathrm{FBP}}} + \left(\frac{\mathrm{DHAP}}{K_{m,\mathrm{dhap}}} + 1\right)\left(\frac{\mathrm{GAP}}{K_{m,\mathrm{gap}}} + 1\right)\right)} \tag{14}$$

$$v_{\mathrm{TPI}} = \frac{\mathrm{TPI_{cor}}\,V_m\left(\mathrm{DHAP} - \frac{\mathrm{GAP}}{K_{\mathrm{eq}}}\right)}{K_{m,\mathrm{dhap}}\left(\frac{\mathrm{DHAP}}{K_{m,\mathrm{dhap}}} + \frac{\mathrm{GAP}}{K_{m,\mathrm{gap}}} + 1\right)} \tag{15}$$

$$v_{\mathrm{GPD}} = \frac{V_m\left(\mathrm{DHAP\,NADH} - \frac{\mathrm{G3P\,NAD}}{K_{\mathrm{eq}}}\right)}{K_{m,\mathrm{NADH}}\,K_{m,\mathrm{DHAP}}\left(\frac{\mathrm{NAD}}{K_{m,\mathrm{NAD}}} + \frac{\mathrm{NADH}}{K_{m,\mathrm{NADH}}} + 1\right)\left(\frac{\mathrm{DHAP}}{K_{m,\mathrm{DHAP}}} + \frac{\mathrm{G3P}}{K_{m,\mathrm{G3P}}} + 1\right)\left(\frac{\mathrm{ADP}}{K_{i,\mathrm{ADP}}} + \frac{\mathrm{ATP}}{K_{i,\mathrm{ATP}}} + \frac{\mathrm{FBP}}{K_{i,\mathrm{FBP}}} + 1\right)} \tag{16}$$

$$v_{\mathrm{HOR2}} = \frac{\mathrm{G3P}\,V_m}{K_{m,\mathrm{G3P}}\left(\frac{\mathrm{G3P}}{K_{m,\mathrm{G3P}}} + 1\right)\left(\frac{\mathrm{PI}}{K_{i,\mathrm{PI}}} + 1\right)} \tag{17}$$

$$V_{\mathrm{GLYCt}} = K_{\mathrm{GLYCt}}\,(\mathrm{GLYC} - \mathrm{GLYCe}) \tag{18}$$

$$v_{\mathrm{GAPDH}} = \frac{\mathrm{GAPDH_{cor}}\,V_m\left(\mathrm{GAP\,NAD\,PI} - \frac{\mathrm{BPG\,NADH}}{K_{\mathrm{eq}}}\right)}{K_{m,\pi}\,K_{m,\mathrm{nad}}\,K_{m,\mathrm{gap}}\left(\left(\frac{\mathrm{BPG}}{K_{m,\mathrm{bpg}}} + 1\right)\left(\frac{\mathrm{NADH}}{K_{m,\mathrm{nadh}}} + 1\right) + \left(\frac{\mathrm{GAP}}{K_{m,\mathrm{gap}}} + 1\right)\left(\frac{\mathrm{NAD}}{K_{m,\mathrm{nad}}} + 1\right)\left(\frac{\mathrm{PI}}{K_{m,\pi}} + 1\right) - 1\right)} \tag{19}$$

$$v_{\mathrm{PGK}} = \frac{\mathrm{PGK_{cor}}\, V_m \, \left(\mathrm{ADP\, BPG}\, K_{\mathrm{eq}} - \mathrm{ATP\, P3G}\right)}{K_{m,\mathrm{P3G}}\, K_{m,\mathrm{ATP}} \left(\frac{\mathrm{BPG}}{K_{m,\mathrm{BPG}}} + \frac{\mathrm{P3G}}{K_{m,\mathrm{P3G}}} + 1\right) \left(\frac{\mathrm{ADP}}{K_{m,\mathrm{ADP}}} + \frac{\mathrm{ATP}}{K_{m,\mathrm{ATP}}} + 1\right)} \tag{20}$$

$$v_{\mathrm{PGM}} = \frac{\mathrm{PGM_{cor}}\, V_m \left(\mathrm{P3G} - \frac{\mathrm{P2G}}{K_{\mathrm{eq}}}\right)}{K_{m,\mathrm{P3G}} \left(\frac{\mathrm{P2G}}{K_{m,\mathrm{P2G}}} + \frac{\mathrm{P3G}}{K_{m,\mathrm{P3G}}} + 1\right)} \tag{21}$$

$$v_{\mathrm{ENO}} = \frac{\mathrm{ENO_{cor}}\, V_m \left(\mathrm{P2G} - \frac{\mathrm{PEP}}{K_{\mathrm{eq}}}\right)}{K_{m,\mathrm{P2G}} \left(\frac{\mathrm{P2G}}{K_{m,\mathrm{P2G}}} + \frac{\mathrm{PEP}}{K_{m,\mathrm{PEP}}} + 1\right)} \tag{22}$$

$$v_{\mathrm{PYK}} = \frac{\mathrm{ADP\, PEP\, PYK_{cor}}\, V_m \left(\frac{\mathrm{PEP}}{K_{m,\mathrm{pep}}} + 1\right)^{\mathrm{hill}}}{K_{m,\mathrm{adp}}\, K_{m,\mathrm{pep}} \left(\left(\frac{\mathrm{PEP}}{K_{m,\mathrm{pep}}} + 1\right)^{\mathrm{hill}} + L \left(\frac{\frac{\mathrm{ATP}}{K_{i,\mathrm{ATP}}} + 1}{\frac{\mathrm{FBP}}{K_{a,\mathrm{FBP}}} + 1}\right)^{\mathrm{hill}}\right) \left(\frac{\mathrm{ADP}}{K_{m,\mathrm{adp}}} + 1\right) \left(\frac{\mathrm{PEP}}{K_{m,\mathrm{pep}}} + 1\right)} \tag{23}$$

$$v_{\mathrm{PDC}} = \frac{\mathrm{PDC_{cor}}\, V_m \left(\frac{\mathrm{PYR}}{K_{m,\mathrm{PYR}}}\right)^{\mathrm{hill}}}{\left(\frac{\mathrm{PYR}}{K_{m,\mathrm{PYR}}}\right)^{\mathrm{hill}} + 1} \tag{24}$$

$$v_{\mathrm{ADH}} = -\frac{\mathrm{ADH_{cor}}\, V_m \left(\mathrm{ETOH\, NAD} - \frac{\mathrm{ACE\, NADH}}{K_{\mathrm{eq}}}\right)}{K_{i,\mathrm{NAD}}\, K_{m,\mathrm{ETOH}} \left(\frac{\mathrm{NAD}}{K_{i,\mathrm{NAD}}} + \frac{\mathrm{NADH}}{K_{i,\mathrm{NADH}}} + \frac{\mathrm{ACE}\, K_{m,\mathrm{NADH}}}{K_{i,\mathrm{NADH}}\, K_{m,\mathrm{ACE}}} + \frac{\mathrm{ACE\, NADH}}{K_{i,\mathrm{NADH}}\, K_{m,\mathrm{ACE}}} + \frac{\mathrm{ETOH}\, K_{m,\mathrm{NAD}}}{K_{i,\mathrm{NAD}}\, K_{m,\mathrm{ETOH}}} + \frac{\mathrm{ETOH\, NAD}}{K_{i,\mathrm{NAD}}\, K_{m,\mathrm{ETOH}}} + \frac{\mathrm{ACE\, ETOH\, NAD}}{K_{i,\mathrm{ACE}}\, K_{i,\mathrm{NAD}}\, K_{m,\mathrm{ET}} [illegible]} [illegible]} \tag{25}$$

$$v_{\mathrm{ETOHt}} = K_{\mathrm{ETOHt}} \left(\mathrm{ETOH} - \mathrm{ETOHe}\right) \tag{26}$$

$$v_{\mathrm{mito}} = \frac{\mathrm{ADP\, PI}\, V_m}{\left(\mathrm{ADP} + K_{m,\mathrm{ADP}}\right) \left(K_{m,\mathrm{PI}} + \mathrm{PI}\right)} \tag{27}$$

$$v_{\mathrm{ATPase}} = \frac{\mathrm{ATP}\, K}{\mathrm{ADP}} \tag{28}$$

$$V_{\mathrm{ADK}} = K \left( \mathrm{ADP}^2 - \frac{\mathrm{AMP}\,\mathrm{ATP}}{K_{\mathrm{eq}}} \right) \tag{29}$$

$$V_{\mathrm{vacPi}} = -K \, (\mathrm{PI} - \mathrm{PIvac}) \tag{30}$$

$$V_{\mathrm{Amd1}} = \frac{\mathrm{AMP}\, V_{\mathrm{Amd1}}}{\mathrm{AMP} + K_{m,\mathrm{AMP}} \left( \frac{\mathrm{PI}}{K_{m,\mathrm{PI}}} + 1 \right)} \tag{31}$$

$$V_{\mathrm{Ade13Ade12}} = \mathrm{IMP}\, K \tag{32}$$

$$V_{\mathrm{Isn1}} = \mathrm{IMP}\, K \tag{33}$$

$$V_{\mathrm{Pnp1}} = \mathrm{INO}\, K \tag{34}$$

$$V_{\mathrm{Hpt1}} = \mathrm{HYP}\, K \tag{35}$$

$$V_{\mathrm{mitoNADH}} = \frac{\mathrm{NADH}\, V_m}{K_m + \mathrm{NADH}} \tag{36}$$

$$V_{\mathrm{sinkG6P}} = \frac{\mathrm{G6P}\, V_m}{\mathrm{G6P} + K_m} \tag{37}$$

$$V_{\mathrm{sinkF6P}} = \frac{\mathrm{F6P}\, V_m}{\mathrm{F6P} + K_m} \tag{38}$$

$$V_{\mathrm{sinkGAP}} = \frac{\mathrm{GAP}\, V_m}{\mathrm{GAP} + K_m} \tag{39}$$

$$V_{\text{sinkP3G}} = \frac{\text{P3G}\, V_m}{K_m + \text{P3G}} \tag{40}$$

$$V_{\text{sinkPEP}} = \frac{\text{PEP}\, V_m}{K_m + \text{PEP}} \tag{41}$$

$$V_{\text{sinkPYR}} = \frac{\text{PYR}\, V_m}{K_m + \text{PYR}} \tag{42}$$

$$V_{\text{sinkACE}} = \frac{\text{ACE}\, V_m}{\text{ACE} + K_m} \tag{43}$$
