## Supplementary material for "Glycolysis revisited: from steady state growth to glucose pulses": SM_sink_reactions.pdf

### Dilution rate dependent sinks

The sink reaction rates were described by phenomenological expressions which closely resembled the experimental data. Michaelis Menten kinetics were used. The dilution rate-dependent $V_max$ was fit to the data using polynomial equations.

The rate equations for the sink reactions were the following:

$$V_{\text{sinkG6P}} = \frac{\text{G6P}\, V_m}{\text{G6P} + K_m} \tag{1}$$

$$V_{\text{sinkF6P}} = \frac{\text{F6P}\, V_m}{\text{F6P} + K_m} \tag{2}$$

$$V_{\text{sinkGAP}} = \frac{\text{GAP}\, V_m}{\text{GAP} + K_m} \tag{3}$$

$$V_{\text{sinkP3G}} = \frac{\text{P3G}\, V_m}{K_m + \text{P3G}} \tag{4}$$

$$V_{\text{sinkPEP}} = \frac{\text{PEP}\, V_m}{K_m + \text{PEP}} \tag{5}$$

$$V_{\text{sinkPYR}} = \frac{\text{PYR}\, V_m}{K_m + \text{PYR}} \tag{6}$$

$$V_{\text{sinkACE}} = \frac{\text{ACE}\, V_m}{\text{ACE} + K_m} \tag{7}$$

The following polynomial fits were derived from the data:

$$V_{m,\text{sinkG6P}} = 3.6854 * d.^3 - 1.4119 * d.^2 - 0.6312 * d - 0.0043 \tag{8}$$

$$V_{m,\text{sinkF6P}} = 519.3740 * d.^6 - 447.7990 * d.^5 + 97.2843 * d.^4 + 8.0698 * d.^3 - 4.4005 * d.^2 + 0.6254 * d - 0.0078 \tag{9}$$

$$V_{m,\text{sinkGAP}} = 170.8447 * d.^6 - 113.2975 * d.^5 + 2.6494 * d.^4 + 10.2461 * d.^3 - 1.8002 * d.^2 + 0.1988 * d + 0.0012 \tag{10}$$

$$V_{m,\mathrm{sinkP3G}} = -0.2381 * d.^2 - 0.0210 * d - 0.0034 \quad (11)$$

$$V_{m,\mathrm{sinkPEP}} = -0.0637 * d.^2 - 0.0617 * d - 0.0008 \quad (12)$$

$$V_{m,\mathrm{sinkPYR}} = -8.4853e+03 * d.^6 + 9.4027e+03 * d.^5 - 3.8027e+03 * d.^4 + 700.5 * d.^3 - 60.26 * d.^2 + 0.711 * d - 0.0356 \quad (13)$$

$$V_{m,\mathrm{sinkACE}} = 118.8562 * d.^6 - 352.3943 * d.^5 + 245.6092 * d.^4 - 75.2550 * d.^3 + 11.1153 * d.^2 - 1.0379 * d + 0.0119 \quad (14)$$

The Michaelis Menten constants were set to the following values:

$$K_{mG6P} = 0.01mM \quad (15)$$

$$K_{mF6P} = 0.0001mM \quad (16)$$

$$K_{mGAP} = 0.0005mM \quad (17)$$

$$K_{mP3G} = 0.001mM \quad (18)$$

$$K_{mPEP} = 0.001mM \quad (19)$$

$$K_{mPYR} = 0.001mM \quad (20)$$

$$K_{mACE} = 0.0001mM \quad (21)$$

Resulting in the fits in (Fig 1):

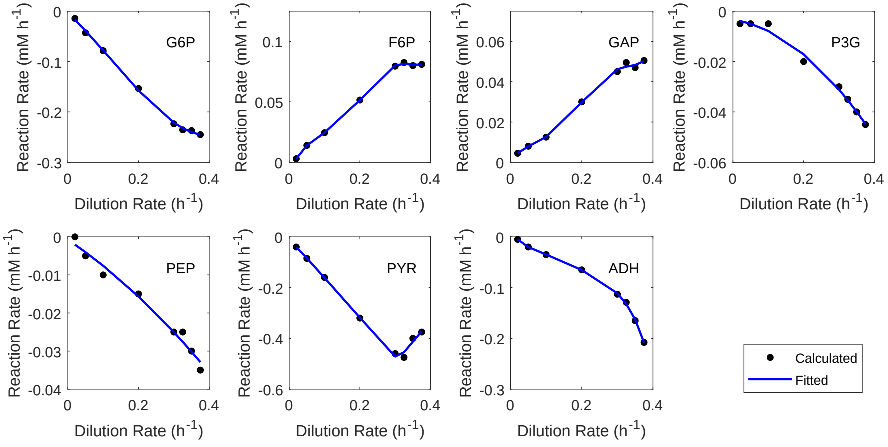


**FIGURE 1 Polynomial fits for the sink reactions.** Each plot contains a sink reaction. Reaction rate is plotted against dilution rate. Fits are shown as a blue line and experimental data points as black dots.
